# Rice soil methane emissions are modulated by microbial cohorts that shape redox succession over the growing season

**DOI:** 10.64898/2026.08.10.744021

**Authors:** Bethany C. Kolody, Junhyeong J. Kim, Rohan Sachdeva, Ling-Dong Shi, Bruce A. Linquist, Jillian F. Banfield

## Abstract

Rice paddies are major anthropogenic methane sources, yet the community interactions regulating emissions remain poorly resolved. This field experiment combined genome-resolved metagenomics, metatranscriptomics, and biogeochemistry to identify microbial cohorts directly and indirectly involved in methane cycling over a rice growing season. Cohorts routed reducing equivalents to different terminal electron-accepting processes and changed in abundance in response to depth and time-dependent shifts in environmental conditions. Methanogenic activity peaked late in the season in the rhizosphere and surface soil, whereas activity in 2-10 cm bulk soil peaked earlier, indicating that methanogenesis was temporally partitioned across soil compartments. *Geobacter*-like extracellular electron transfer was the most highly expressed alternate sink for reducing equivalents. Methane consumption was vertically partitioned between surface type I methanotrophs and deeper type II methanotrophs and anaerobic methane-oxidizing archaea. We infer that methane emissions reflect seasonal shifts in microbial cohorts that redirect methanogenic substrates through competing redox processes.

## Introduction

Rice cultivation generates around 8% of all anthropogenic methane emissions^1^. Methane is an attractive mitigation target because its high warming potential and short atmospheric lifetime relative to CO₂ mean that emission reductions can rapidly slow warming^2^. In rice paddies, methane is produced by anaerobic methanogenic archaea after flooding depletes more energetically favorable electron-accepting pathways^3^.

Upon flooding, oxygen is depleted within hours to days due to microbial respiration, and anaerobic carbon mineralization proceeds through a thermodynamically ordered succession of terminal electron–accepting processes (TEAPs): nitrate reduction, Mn(IV) reduction, Fe(III) reduction, sulfate reduction, and finally methanogenesis^3,4^. Complex carbon is hydrolyzed into monomers which are then fermented into alcohols, fatty acids, and the precursors for methanogenesis, including acetate, CO_2_, and H_2_. The conversion of the remaining substrates into methanogenic precursors depends on a syntrophic secondary fermentation, in which fermenters rely on growing methanogens to keep H_2_ partial pressures low enough for the reactions to be favorable^5^. Initially, plant residue is the primary source of organic matter, but as rice plants grow, their root exudates, which change in composition by life stage, become an important contributor^3^. Methane emissions rise as plants grow, peak around heading and flowering and decline toward maturity^6,7^. Seasonal methane oxidation profiles are inconsistent^8,9^ and methane fluxes are often most strongly controlled by methane production, which also peaks in later reproductive stages^8–12^.

Microbial communities in paddy soils are spatially partitioned. Oxygen penetrates the top millimeters of the bulk soil^13^ and the zone surrounding rice roots (rhizosphere), which are aerated by the specialized aerenchyma vasculature of the rice plant^14^. Aerobic processes such as methane oxidation by particulate methane monooxygenase (pMMO)-utilizing methanotrophs, nitrification, and aerobic heterotrophic respiration occur in these zones. Immediately below, sharp transitions to anoxia support denitrification and metal reduction^3^, while the bulk soil remains strongly reduced and methanotrophy occurs anaerobically^15^. Less is known about microbial composition and function in the deep bulk soil, but microbial biomass and organic matter both decline with depth, and deep-soil community structure is likely influenced both by changes in carbon substrate availability^16^ and soil geochemistry^17^.

While the microbiology of rice paddies has been intensively investigated^3,18–20^, it remains unclear how interacting microbial populations partition reducing equivalents across the distinct yet interconnected soil habitats that structure paddy ecosystems. Many field-based studies focus on individual soil compartments, e.g. the rhizosphere^21,22^, or on specific functional groups. For example, methanogens are frequently profiled using the *mcrA* marker^23^, whereas other studies target nitrogen^24^ or sulfur^25^ cycling in isolation. Unlike marker gene surveys, genome-resolved metagenomics and metatranscriptomics can determine which metabolic processes are active within the same organism or cohort of interacting organisms. Ultimately, methane fluxes emerge from the collective dynamics of interacting respiratory and fermentative cohorts that partition reducing equivalents across competing TEAPs and methanogenesis.

Studies of the whole microbial community underpinning methane emissions have been hampered by the complexity of soil systems, which makes genome reconstruction difficult^26^. Here, we deeply sequenced the rhizosphere and bulk soil microbial community of a medium-grain rice paddy soil across a growing season in Northern California. In addition to profiling the overall transcriptional activity of TEAPs, we reconstructed the genomes of the most important players and grouped them into cohorts based on shared abundance patterns across soil depth and time, and used metatranscriptomes to infer metabolic activity. We investigated how redox processes, methanogenesis, and methanotrophy are partitioned among cohorts and change over space and time. We evaluated which TEAPs are most important for diverting electrons away from methanogenesis, and how stable interdependencies among microbial populations are across soil compartments. This approach enabled a systems-level view of how interacting microbial cohorts regulate methane emissions in field-grown rice.

## Results and Discussion

### Soil sampling and genome recovery

We conducted a field experiment in a 25 m x 85 m rice paddy planted to a Calrose medium grain variety (M-206) at the Rice Experiment Station (RES) in Biggs, CA. We collected samples for metagenomics/metatranscriptomics, soil chemistry, and trace gas fluxes at seven timepoints between May and October 2021. At six timepoints spanning the seedling to harvest maturity growth stages, we sampled rice rhizospheres from three plant replicates and bulk soil controls from five depths (0 - 2 cm, 2 - 10 cm, 10 - 20 cm, 20 - 30 cm, and 50 - 60 cm) spanning the plowed surface Ap horizon and underlying Bss horizons^27,28^ of adjacent plant-excluded soil (Fig. 1A). In May, before seeding, we sampled unplanted control soil. Trace gas fluxes were measured over rice plants and bulk soil using a Picarro gas analyzer, and contextualized with higher-resolution methane flux measurements from an adjacent basin with the same cultivar and flood regime from 2021–2023 (Fig. S1, Data S1)^29^. The clay-rich, low-permeability Esquon-Neerdobe complex soil promoted strong redox zonation (Fig. S2)^30^. Consistent with this stratification, total carbon, total nitrogen, DTPA-extractable copper, iron, and zinc, and sulfate all decreased with depth in the bulk soil (Fig. 1B, Fig. S3, Data S1).

**Fig. 1.**
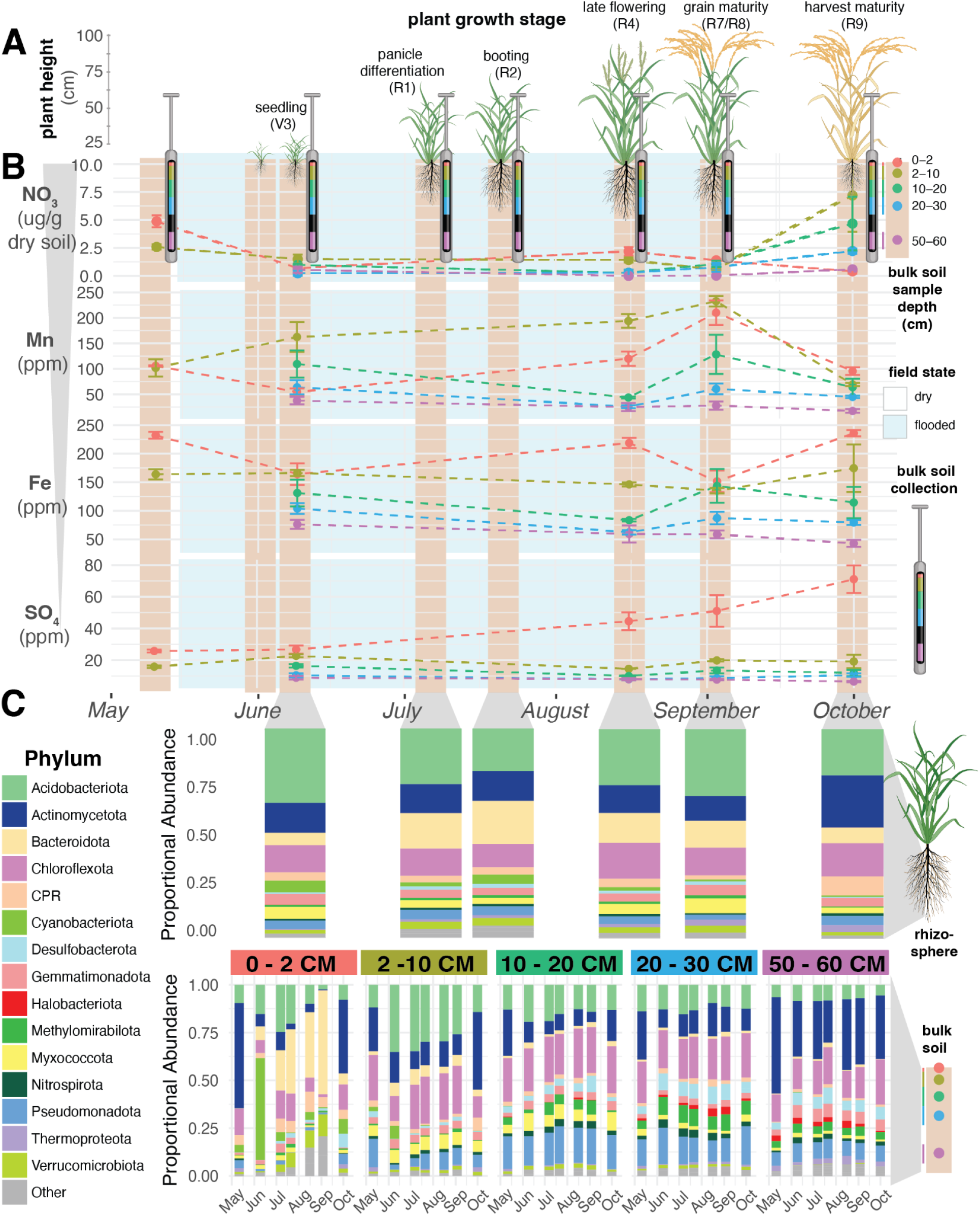
Changes in soil chemistry and the microbial community across the growing season and plant growth stage. Rhizosphere and bulk soil were sampled over the 2021 growing season, starting before seeding and continuing through post-harvest field drying. **(A)** Plant height across the growing season. **(B)** Mean concentrations (± standard error) of electron acceptors shown in order of decreasing redox potential: nitrate, DTPA-extractable manganese and iron, and calcium phosphate-extractable sulfate (labile SO₄²⁻). Electron acceptor concentrations are plotted for each soil depth interval and colored by compartment: red (0–2 cm), olive (2–10 cm), green (10–20 cm), blue (20–30 cm), and purple (50–60 cm). Nitrate was measured via a KCl extract of fresh soil and all other measurements were made on air-dried soil. **(C)** Phylum-level taxonomy of curated genomes from the rhizosphere (top panel) and bulk soil (bottom panel) over time.

We sequenced 122 metagenomic samples, generating 1.65 Tbp of Illumina data and recovering 204 high-quality (> 90% completeness, < 5% contamination) and 525 medium quality (≥ 50% completeness, < 10% contamination) genomes dereplicated at 95% ANI (Data S2). Phylum-level microbial community structure was strongly depth-stratified, with the greatest seasonal variation at the 0-2 cm bulk soil oxic/anoxic interface, where an early-season cyanobacterial bloom was followed by a gradual increase in Bacteroidota. Actinomycetes were most common during dry periods and in deep soil (Fig. 1 C), and *Methanoperedens nitroreducens* sp. anaerobic methanotrophs (Halobacteriota, red) represented a noticeable fraction of the community below 20 cm. We also generated 839 Gb of metatranscriptomic data from rhizosphere samples and the top two bulk soil layers, below which RNA yields were insufficient for sequencing. To capture activity from organisms without recovered genomes, we built a 95% identity dereplicated protein set (n= 64,950,587 proteins) using all metatranscriptomic and metagenomic scaffolds and searched it with HMM markers for dominant TEAPs (Fig. S4).

### Diverse and pervasive methanogens

Methanogens were active across soil compartments at all sampled time points and were highly diverse. We detected expression of 740 genes from methanogenic species-level McrA protein clusters, defined at 95% amino acid identity, yet the top 10 clusters accounted for 21.4% of *mcrA* expression (Extended Data Fig.1A, Data S3). Despite this diversity, *mcrA* expression patterns were highly conserved across clusters, and were more strongly correlated than expression patterns of any other TEAP (Fig. S5). Hydrogenotrophic lineages, particularly *Methanocellales*, accounted for the largest share of *mcrA* activity (Extended Data Fig.1B, Extended Data Fig. 1C). This agrees with recent Northern California rice datasets, suggesting that *Methanocellales*-driven hydrogenotrophic methanogenesis may be a recurring feature of this regional rice-growing system^8,31^.

### Subsurface bulk soil shows an earlier methanogenic activity peak

Years of methane monitoring at the Rice Experiment Station show an initial post-flooding lag, followed by high fluxes through much of the growing season and eventual decline^29^. Our methane measurements followed this trend (Fig. 2A). However, methanogenic activity in the rhizosphere and 0–2 cm bulk soil did not peak until late season, creating an apparent mismatch between methane fluxes and methanogenesis in compartments often considered dominant methane sources^32,33^. Methane consumption does not resolve this discrepancy, because *pmoB* expression was also highest early in the season (Fig. 2B, Fig. 2C). Instead, early fluxes coincided with peak methanogenic *mcrA* expression in the 2–10 cm bulk soil, suggesting that this interval is a major site of early-season methanogenic activity (Fig. 2D). Supporting this interpretation, anaerobic methanotrophs, which could mitigate methane release from depth, had peak activity and abundance late in the growing season (Extended Data Fig. 1C, Fig. S6). Although overall microbial activity declines with soil depth^17^, the outsized volume of subsurface bulk may still make it a major source of early-season methane.

**Fig. 2.**
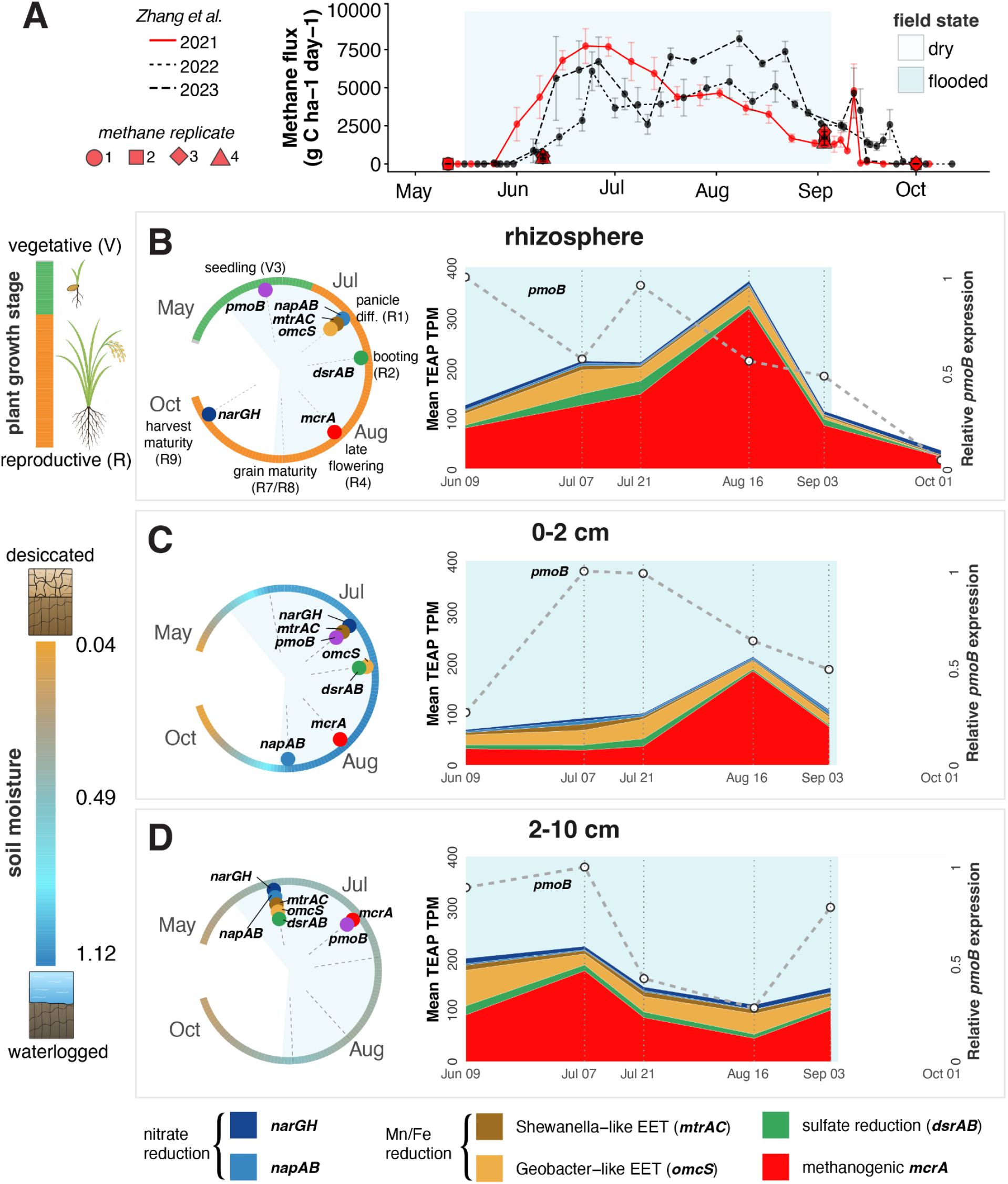
Seasonal methane emissions alongside major pathways generating, competing with, and consuming methane across soil compartments. **(A)** Mean methane fluxes (± standard error) from our rice basin (large red points) are shown alongside methane data from the same cultivar grown in an adjacent basin in 2021 (red), 2022, and 2023^29^. Dates in which the field was flooded are shaded blue. **(B-D)** Left: timing of peak expression of gene markers for methanogenesis, TEAPs competing with methanogenesis, and methanotrophy relative to plant growth stage in the rhizosphere (**B**) and to soil moisture in the 0-2 cm (**C**) and 2-10 cm (**D**) bulk soil. Right: Relative contribution of TEAPs across the growing season by soil compartment (y-axis; left) and aerobic methanotrophy as measured by *pmoB* (y-axis, right). Marker genes were identified from all metatranscriptomic and metagenomic scaffolds, independent of genome binning. Marker abbreviations are: *mcrA*, methyl-coenzyme M reductase subunit A; *narGH*, membrane-bound nitrate reductase; *napAB*, periplasmic nitrate reductase; *mtrAC*, metal-reducing decaheme cytochrome complex; *omcS*, extracellular multiheme cytochrome; *dsrAB*, dissimilatory sulfite reductase; and *pmoB*, particulate methane monooxygenase subunit B.

### Depth-specific TEAP succession shifts the timing of methanogenesis

Methanogens require H_2_ or small organic molecules that can also fuel other anaerobic respiratory processes like nitrate, iron, and sulfate reduction. Across soil compartments, peak methanogenesis expression occurred after peak expression of TEAPs that compete for organic substrates, consistent with a redox succession in which methanogenesis intensifies as alternative terminal electron acceptors are depleted. However, the timing and degree of separation among TEAPs differed sharply among soil compartments (Fig. 2B-D, Extended Data Fig. 2).

In the rhizosphere, expression of the dominant nitrate respiration marker, *narGH*, was high early in the season before nitrate is fully drawn down by anaerobic metabolism and highest during dry down when oxic microsites can regenerate nitrate. Periplasmic nitrate reduction (*napAB*) and manganese and iron reduction (*Geobacter*-like EET, *omcS*, *Shewanella*-like EET, *mtrAC*) gene expression peaked simultaneously in early July during panicle differentiation, followed by sulfate reduction (*dsrAB*) in late July during booting, and methanogenic *mcrA* in mid-August during late flowering (Fig. 2B, Extended Data Fig. 2). This timing is consistent with previous studies that saw peak methanogenesis in mid-late reproductive stages^8,9^.

In the 0-2 cm bulk soil, TEAP expression followed a similar trajectory to that of the rhizosphere, although *Geobacter*-like EET was decoupled from *Shewanella*-like EET and peaked alongside sulfate reduction in late July. As in the rhizosphere, methanogenic *mcrA* expression peaked in mid-August (Fig. 2C, Extended Data Fig. 2).

In 2–10 cm bulk soil, methanogenic mcrA expression peaked earlier, in early July. Unlike in the rhizosphere or 0–2 cm bulk soil, competing TEAPs were synchronized, with marker expression high in mid-June, declining over time, and dipping during peak methanogenesis (Fig. 2D, Extended Data Fig. 2).

The contrast in behavior of the 2-10 cm compared to shallow soil may reflect more persistent moisture and reducing conditions in 2–10 cm soil. Seasonal desiccation of the 0-2 cm soil may regenerate electron acceptor pools and prolong the redox cascade after flooding, whereas 2–10 cm soil likely contains fewer oxidized microsites and transitions to methanogenesis more rapidly. Substrate availability may also differ by depth. Methanogenesis in the rhizosphere and surface bulk soil were predominantly hydrogenotrophic, whereas the early 2–10 cm bulk soil methanogenesis peak was driven largely by Methanotrichales, likely acetoclastic methanogens.

Because straw-incorporation can increase acetoclastic methanogenesis^34^, this depth-specific shift suggests that 2-10 cm methanogens accessed a distinct substrate pool, potentially acetate derived from slow fermentation of rice residues from the previous growing season. The tendency for 2-10 cm subsurface soil to maintain reducing conditions may also partly explain why methane fluxes have been observed to continue after drainage^29^.

### Genome cohorts reveal environmental and functional specialization

To resolve the microbial community dynamics underlying seasonal methane emissions, we recovered genomes for 80% of the top 100 and 62.3% of the top 1000 taxa detected via ribosomal protein S3 (rpS3) single-copy gene marker analysis (Data S4). We then used weighted correlation network analysis (WGCNA) to group genomes with shared abundance patterns across depth and season. Both rpS3 marker profiles and genome-resolved analyses showed that community composition was structured primarily by depth (Fig. S7, Fig. 3A). WGCNA resolved genomes into 26 cohorts, each assigned a color designation (Fig. 3, Fig. S8-S34). Thirteen cohorts were significantly enriched in specific functional genes based on KEGG Orthology enrichment using Fisher’s exact tests, reported as odds ratios (OR; FDR < 0.05; Data S5).

**Fig. 3.**
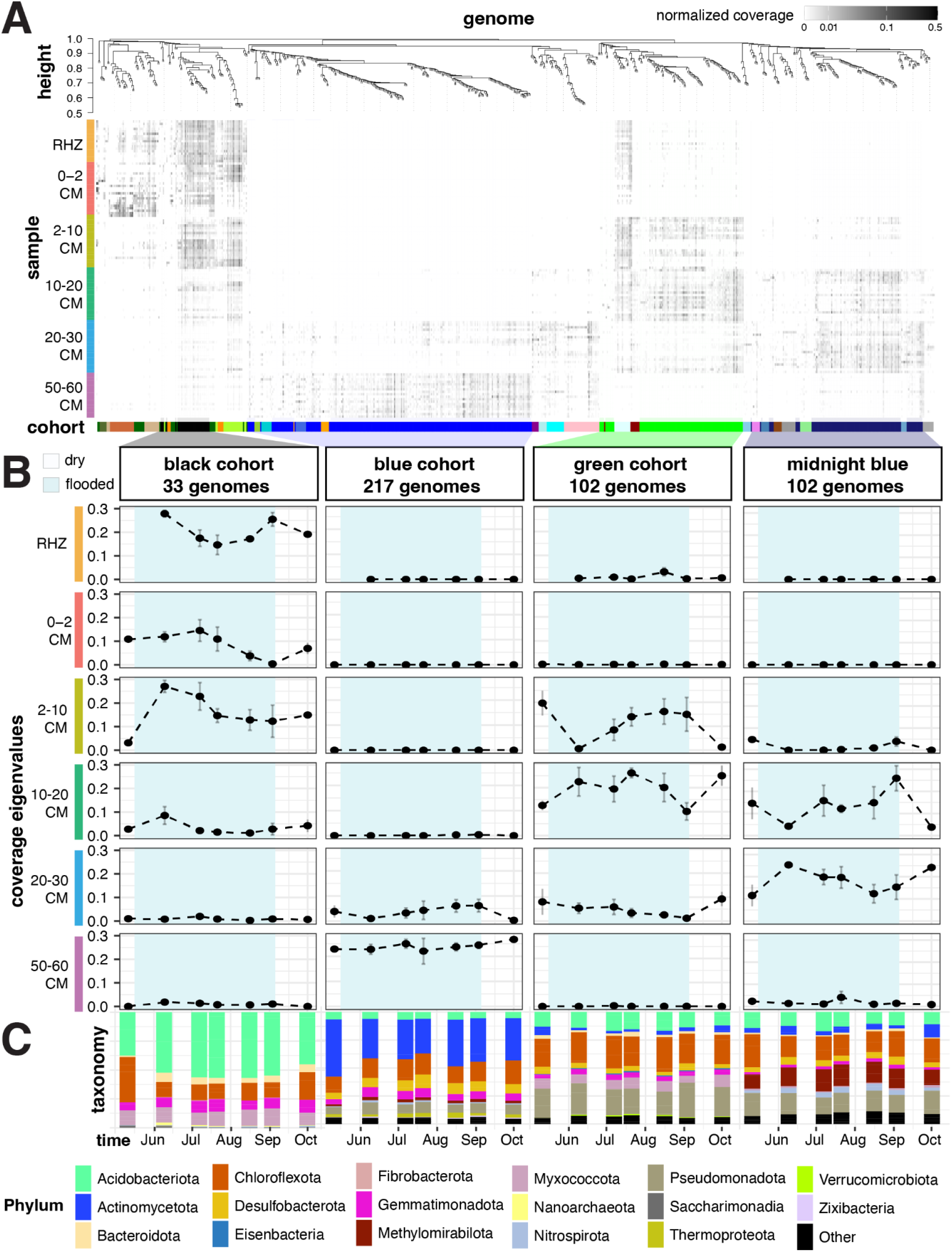
Soil compartments harbor distinct microbial genome signatures that change over time. **(A)** Heatmap of microbial genome abundance. Genomes (x-axis) are grouped according to co-occurance as measured by WGCNA clustering (top dendrogram) and colored by WGCNA cohort membership (bottom). Samples (y-axis) are grouped by soil compartment. **(B)** Average abundance profiles (coverage eigenvalues) across time (mean ± SE) for the four cohorts encompassing the most genomes (columns), shown in separate panels by soil compartment. Shaded blue rectangles indicate flooded period. **(C)** Phylum-level taxonomic composition of genomes pertaining to top four cohorts across the growing season.

#### Soil depth structures genome cohorts

Genome cohorts generally spanned tens of centimeters, crossing multiple soil compartments, but showed replicable seasonal trends that differed by compartment. Eight cohorts were surface-associated, with abundance profiles negatively correlated with depth and pH and positively correlated with organic matter, moisture, and nutrients, whereas the remaining 18 cohorts showed the opposite pattern and were associated with deeper soil (Fig. S35). These trends likely reflect the strong depth dependence of pH, organic matter, and nutrient composition.

#### Drydown selects for desiccation-adapted and sulfur-cycling cohorts

Some cohorts were strongly associated with the seasonal flood regime. Cohorts spanning multiple soil depths were abundant either during flooded periods (e.g. darkgreen, pink, sienna3, tan, and cohorts; Fig. S11, S26, S29, S32) or after drydown (e.g. darkolivegreen, darkorange, and greenyellow cohorts; Fig. S14, S15, S19). For example, the darkorange cohort increased after drydown in the rhizosphere, 0-2 and 2-10 cm bulk soil and consisted of genomically reduced, putatively episymbiotic Saccharibacteria and Paceibacteria (Fig. S15). This is consistent with previous observations that strong shifts in soil conditions can favor Saccharibacteria over other CPR lineages^35^. The darkorange cohort was enriched in functions consistent with osmotic and desiccation stress recovery, including resuscitation-promoting factor B (K21688; OR 32.4) and a large-conductance mechanosensitive channel (K03282; OR 6.8; Data S5), which can relieve hypoosmotic shock. After drydown, these Saccharibacteria consistently expressed type IV pili, which are involved in host attachment^36^, as well as genes involved in DNA replication, cell division, ribosome maturation, and complex carbon degradation (Fig. S36), suggesting that they exploit drydown-associated shifts in host physiology to attach and grow.

The darkolivegreen cohort also rose in abundance after drydown, but only in the surface bulk soil (Fig. S14). This cohort included desiccation-tolerant *Nostoc*, a *Gracilibacteria*, and a 94% complete cable bacteria genome most closely related to *Candidatus* Electronema aureum ENR-cMAG (81.4% rpS3 nucleotide identity; genomes 90, 127, 556, and 245)^37^. Cable bacteria form centimeter-scale conductive filaments that couple sulfide oxidation in reduced soil to electron-accepting processes near oxic interfaces, and a recent greenhouse experiment found that a *Ca.* Electronema enrichment decreased rice methane emissions by 93%^38^. Although *Ca*. Electronema has been detected in rice soils using FISH^39^ and 16S rRNA amplicon sequencing^40^, to our knowledge, genome 245 is the first cable bacteria genome recovered from a rice paddy. It encodes characteristic PilA nanowire proteins and several multi-heme cytochromes (MHCs) with up to 11 heme-binding domains. Like other cable bacteria genomes, it lacks common sulfide-oxidizing genes and instead encodes the dissimilatory sulfur reduction pathway, which is expected to operate in reverse^41^. Genome 245 also encodes *napAB*, which *Ca.* Electronema aureum uses to couple nitrate reduction to sulfide oxidation^42^, and may allow it to perform sulfide oxidation without making contact with fully oxidized soil. The presence of this cable bacteria in a post-drydown surface-soil cohort suggests that oxic microsites may enhance cable bacteria growth, enabling sulfide oxidation and potentially contributing to the observed sulfate accumulation in 0–2 cm bulk soil (Fig. 1B).

#### Subsoil and rhizosphere cohorts encode habitat-specific resource-use strategies

Other cohorts were associated with a single depth across time. For example, the diverse blue cohort (217 genomes) was stably abundant over time at 50 – 60 cm in the lower Bss horizons (Fig. S9). Blue cohort genomes were enriched in mercury resistance genes, including alkylmercury lyase (K00221; OR 11.9) and mercuric reductase (K00520; OR 5.9), as well as sporulation machinery (e.g. K06381; OR 2.4; Data S5), suggesting a persistent deep-subsoil community adapted to metal stress and low-resource conditions.

The black, darkgreen, greenyellow, lightcyan, and tan cohorts were most abundant in the rhizosphere, suggesting that plant-associated carbon inputs may shape their dynamics (Fig. S8, S11, S19, S21, S32). Consistent with this interpretation, the tan cohort was enriched in genes for uptake and metabolism of deoxy sugars found in plant-associated polysaccharides, including fucose (L-fuculokinase, K00879; OR 24.5), which can be abundant in root mucilage glycans^43^, and rhamnose (L-rhamnose-H^+^ transport protein, K02856; OR 9.2; Data S5), a constituent of pectic polysaccharides such as rhamnogalacturonans^44^. Together, these patterns indicate that paddy soils host functionally distinct cohorts shaped by soil geochemistry, flood regime, and plant inputs.

### Stable genome ranks suggest conserved interactions within cohorts

Within each cohort, genome ranks were more stable across soil compartments than expected by chance. For example, organisms that were most abundant within a cohort in the 0-2 cm oxic/anoxic interface often remained most abundant at all other soil depths, even in the 50-60 cm bulk soil (Fig. 4). This rank stability is not required by WGCNA, which clusters genomes based on similar abundance patterns rather than fixed proportions across samples. Conserved genome ranks across chemically and physically distinct soil compartments therefore suggest consistent functional coupling and possible metabolic interdependence within cohorts. However, rank stability varied by cohort and compartment, with more distant soil compartments showing greater rank shuffling and lower correlations (Fig. 4B–D). Some of the strongest rank correlations occurred within the methanogen-containing greenyellow cohort, including between 2–10 cm and 50–60 cm bulk soil (ρ = 0.96; Fig. S37, Fig. S38), and within the methanotroph-containing black cohort, including between the rhizosphere and 0–2 cm bulk soil (ρ = 0.95; Fig. S39). Similar conservation of partner abundance has been observed in syntrophic communities, where metabolic exchange can constrain the relative abundances of interacting taxa. For example, co-cultures of butyrate-oxidizing syntrophs and hydrogenotrophic methanogens converged toward relatively constant cell ratios despite differing starting abundances^45^.

**Fig. 4:**
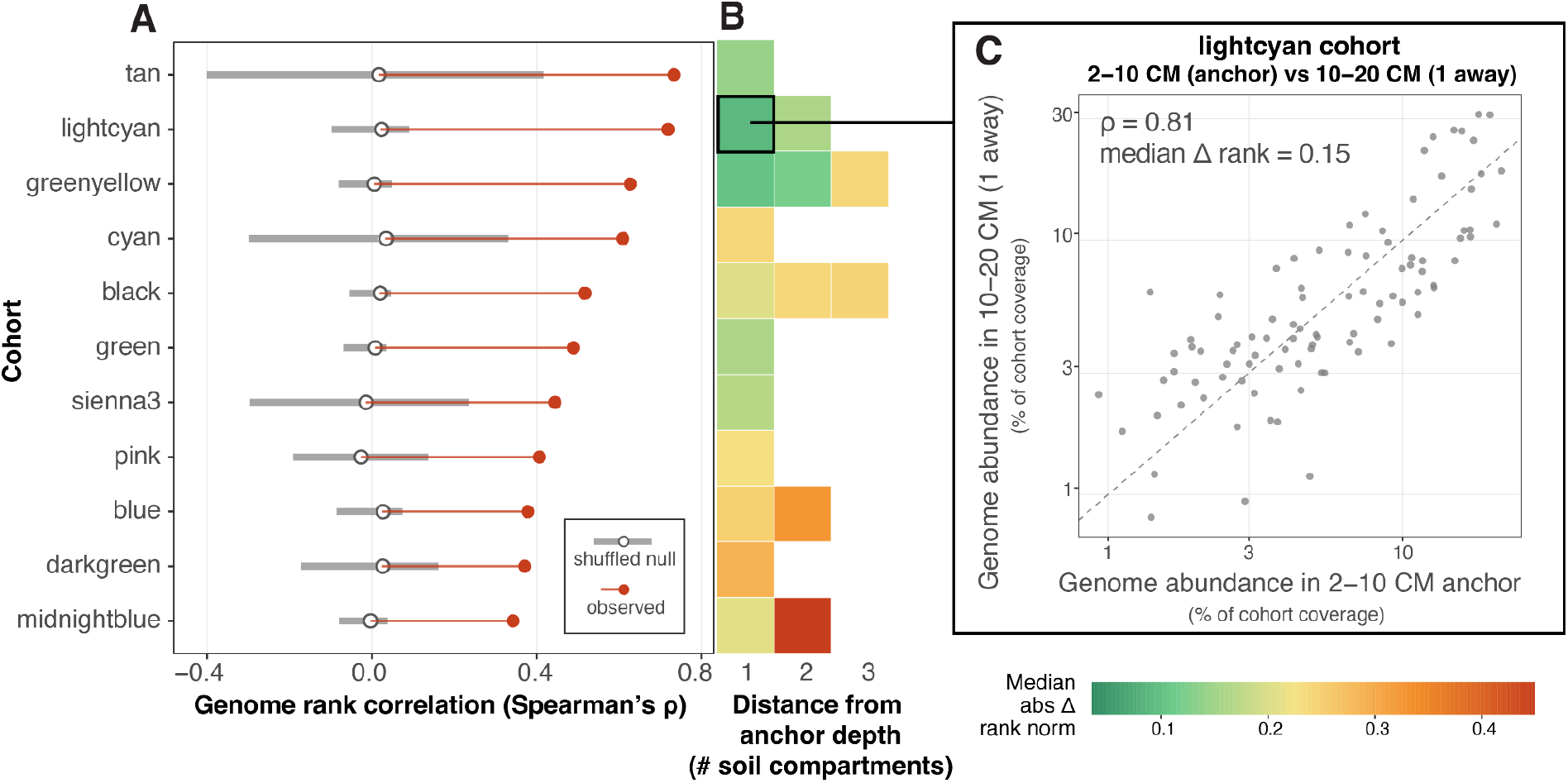
Evidence for persistence of rank within cohorts. **(A)** Within-cohort genome rank conservation relative to a shuffled null model. For each cohort, genomes were ranked within each sampling date and bulk soil compartment by their percent contribution to total cohort abundance and compared across soil compartment pairs. Red points show the observed pooled Spearman correlation (rho) of within-cohort ranks across compartment pairs, open circles show the median null expectation, and grey horizontal bars show the 95% permutation interval after shuffling genome identities within each cohort × date × compartment-pair, thereby preserving cohort size and marginal rank distributions while disrupting cross-compartment correspondence. **(B)** Rank displacement from each cohort’s dominant bulk-soil compartment (rhizosphere excluded). For each cohort, the anchor depth was defined as the bulk-soil compartment with the highest mean cohort abundance. Heatmap values show the median absolute normalized rank shift between the anchor depth and other bulk-soil compartments, grouped by the number of compartments separating them (1 away, 2 away, 3 away). Color scale indicates median absolute delta rank norm, calculated as the absolute difference in within-cohort rank between two compartments, normalized by cohort size and summarized as the median across matched genome-date observations (warmer colors indicate more rank shuffling between soil compartments). This panel includes only cohort-by-depth comparisons in which at least five genomes were ranked within each cohort-date-depth sample, at least five matched genome-date pairs were available for the summarized comparison, and the non-anchor comparison depth had a mean cohort abundance of at least 0.015. **(C)** Exemplar scatterplot comparing each genome’s relative contribution to cohort abundance for anchor-versus-distance comparisons. The dashed line indicates a 1:1 relationship across soil compartments.

### Cohorts differ in how they route substrates through competing redox pathways

Broad gene surveys showed that pathways supporting and competing with methanogenesis were encoded by many cohorts (Extended Data Fig. 3). However, cohort activity changed across the growing season (Fig. 5A), and TEAP expression varied strongly by cohort, consistent with distinct physicochemical niches (Fig. 5). For example, the tan, darkgreen, and sienna3 cohorts all expressed genes for complex carbon degradation, but coupled them to the expression of distinct TEAPs: sulfate reduction, sulfate and iron reduction, and nitrate/nitrite reduction, respectively (Data S6). We conclude that many organisms have the capacity to draw down substrates used as inputs to methane production, but the electron acceptor they use depends on local physicochemical conditions and cohort context.

**Fig. 5:**
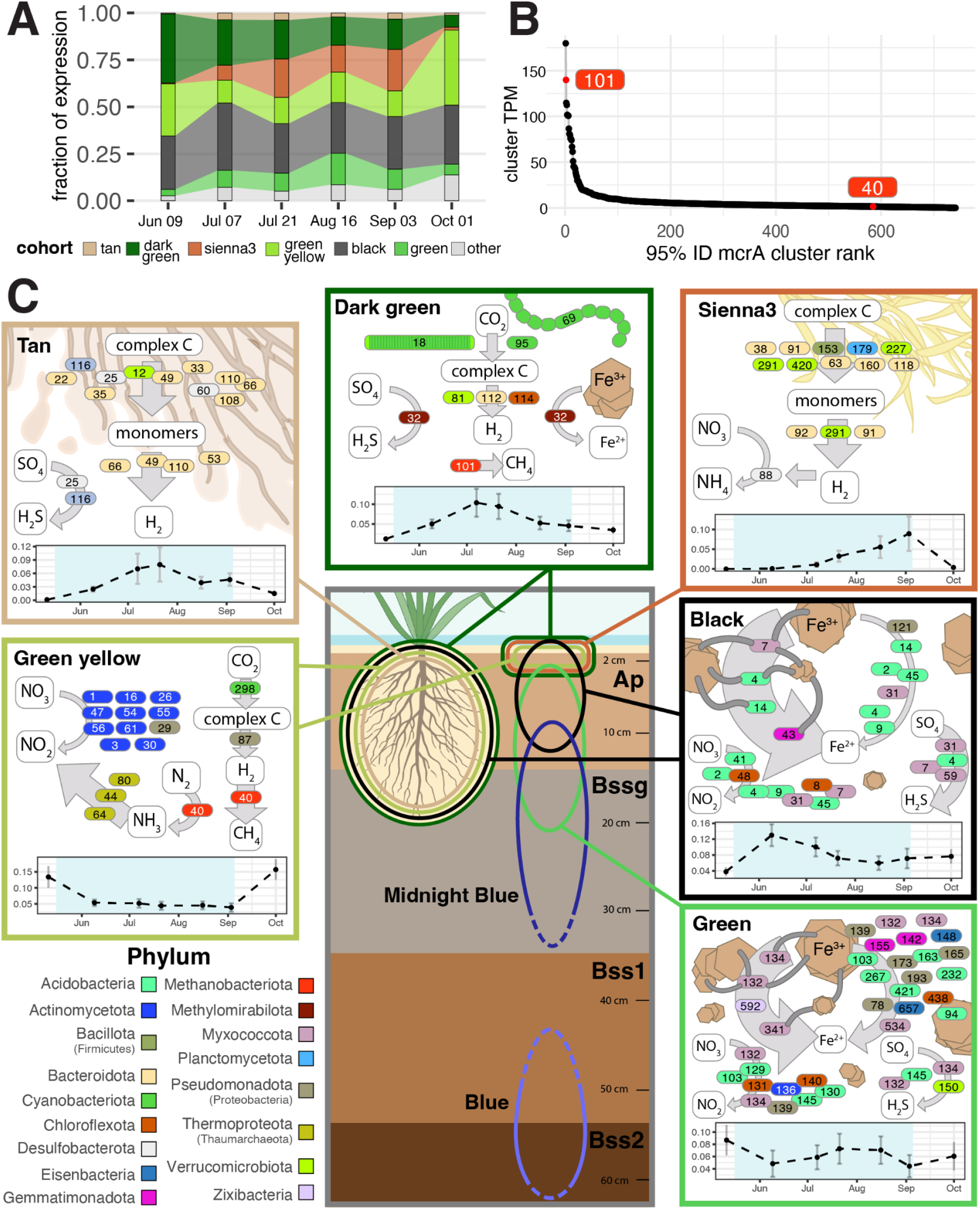
Microbial activities upstream of methanogenesis differ by soil compartment. **(A)** Relative contribution of each cohort to gene expression over time in the rhizosphere and 0 - 10 cm bulk soil. **(B)** Rank-abundance of expressed 95% ID dereplicated protein clusters corresponding to McrA from methanogens (black points). Red dots represent expression from McrB clusters pertaining to binned methanogens, for context. **(C)** Cartoon summary of genomes expressing major processes linked to carbon cycling. Genomes are represented by ovals colored by phylum and numbered (full genome information is in Data S1), and placed alongside arrows according to transcriptomic evidence of various metabolic processes supporting carbon transformations. For example, genomes alongside arrows indicating hydrogen production expressed H_2_-evolving hydrogenases. For iron reduction, arrows with grey nanowire cartoons indicate expression of *Geobacter*-like extracellular electron transfer (EET) machinery and those without indicate expression of *Shewanella*-like EET machinery. Genomes are grouped into pop-out panels by cohort, with the average cohort abundance profile given at the bottom of the panel. The primary location of each cohort in the soil is circled in the center panel. Midnight Blue and Blue cohorts occur below the region in which we recovered RNA, but were included because they harbor many genomes. Dashed lines indicate that these cohorts may continue into unsampled soil depths.

### Methanogens from distinct cohorts converge in seasonal mcr expression

We recovered two medium-quality hydrogenotrophic *Methanobacterium* genomes. Both expressed *mcr* following the same seasonal trends observed in genome-independent McrA analysis across all soil compartments, despite having distinct abundance patterns (Fig. S40, Fig. 2 B-D). Genome 101 was among the most highly expressed methanogens detected (Fig. 5B) and belonged to the darkgreen cohort (Fig. S11), which peaked mid-season and was a principal contributor to complex carbon degradation, including carbon fixed by cyanobacterial members (Fig. 5C). Genome 40 belonged to the greenyellow cohort (Fig. S16), which was dominated by actinomycetes and abundant during dry periods before and after flooding. Previous studies have reported discrepancies between methanogen abundance and *mcr* expression levels^23,46,12^. Our finding that methanogens with distinct abundance dynamics converge in seasonal activity may help explain this disconnect: mcr operon expression may directly reflect H₂ availability ^47^, whereas growth rate depends on substrate competition and physiological responses to local conditions such as desiccation.

### Plant-polymer degrading cohorts produce H_2_ in distinct soil habitats

Because H_2_ was the dominant substrate for methanogenesis, we functionally classified hydrogenases from our genome set using HydDB (Fig. S41)^48^. The most highly expressed H_2_-generating hydrogenases were Group A1 prototypical fermentative [NiFe] hydrogenases, primarily from Bacteroidata (Fig. S42; Data S7). While some of this expression originated from the two cohorts where we recovered methanogens, most came from the non-methanogenic sienna3 and tan cohorts (Fig. S43), suggesting that H₂-producing fermenters can follow distinct spatiotemporal abundance patterns from hydrogenotrophic methanogens. However, syntrophic coupling between fermentative bacteria and methanogens is well established in rice paddy soils^3,5^, and genome-independent marker data indicate that these compartments harbor rare methanogens not represented in recovered cohorts. Consistent with this, the tan cohort includes a Syntrophorhabdaceae genome (genome 111) related to lineages with obligate syntrophic associations with hydrogenotrophic methanogens that maintain favorable H₂ partial pressures^49^. Both the sienna3 and tan cohorts expressed enzymes targeting plant material like starch, hemicellulose, cellulose, chitin, and other complex polysaccharides (Fig. S44, Fig. S45, Data S6). However, their abundance patterns diverged: tan was most abundant in the rhizosphere and peaked mid-season, coinciding with elevated root exudate release in rice^50^, whereas sienna3 was most abundant in 0–2 cm bulk soil, consistent with utilization of rice straw-derived plant material. Thus, H₂ production was not strictly colocalized with methane production, and H₂-producing cohorts may be fueled by distinct plant-carbon pools.

### Redox heterogeneity sustains concurrent TEAP expression across soil compartments

TEAPs are often conceptualized as a spatially ordered sequence proceeding from high- to low-redox-potential electron acceptors with distance from oxic zones. However, markers for nitrate reduction, sulfate reduction, and *Geobacter*- and *Shewanella*-like extracellular electron transfer were concurrently expressed across the rhizosphere, 0–2 cm, and 2–10 cm bulk soil throughout the growing season (Fig. 6; Data S8). Redox pathways were also expressed beyond the soil regions where they are considered favorable. For example, nitrate reduction was expressed throughout the growing season in 2–10 cm bulk soil, well below the millimeter-scale oxic/anoxic interface, implying that high-redox-potential microenvironments persist months into the flooding regime. Soil redox potential is heterogeneous on the microscale^51^, and in fine-textured rice clay, aggregate structure, tortuous diffusion pathways, and spatially variable organic matter may promote steep microscale redox gradients^52,53^.

**Fig. 6.**
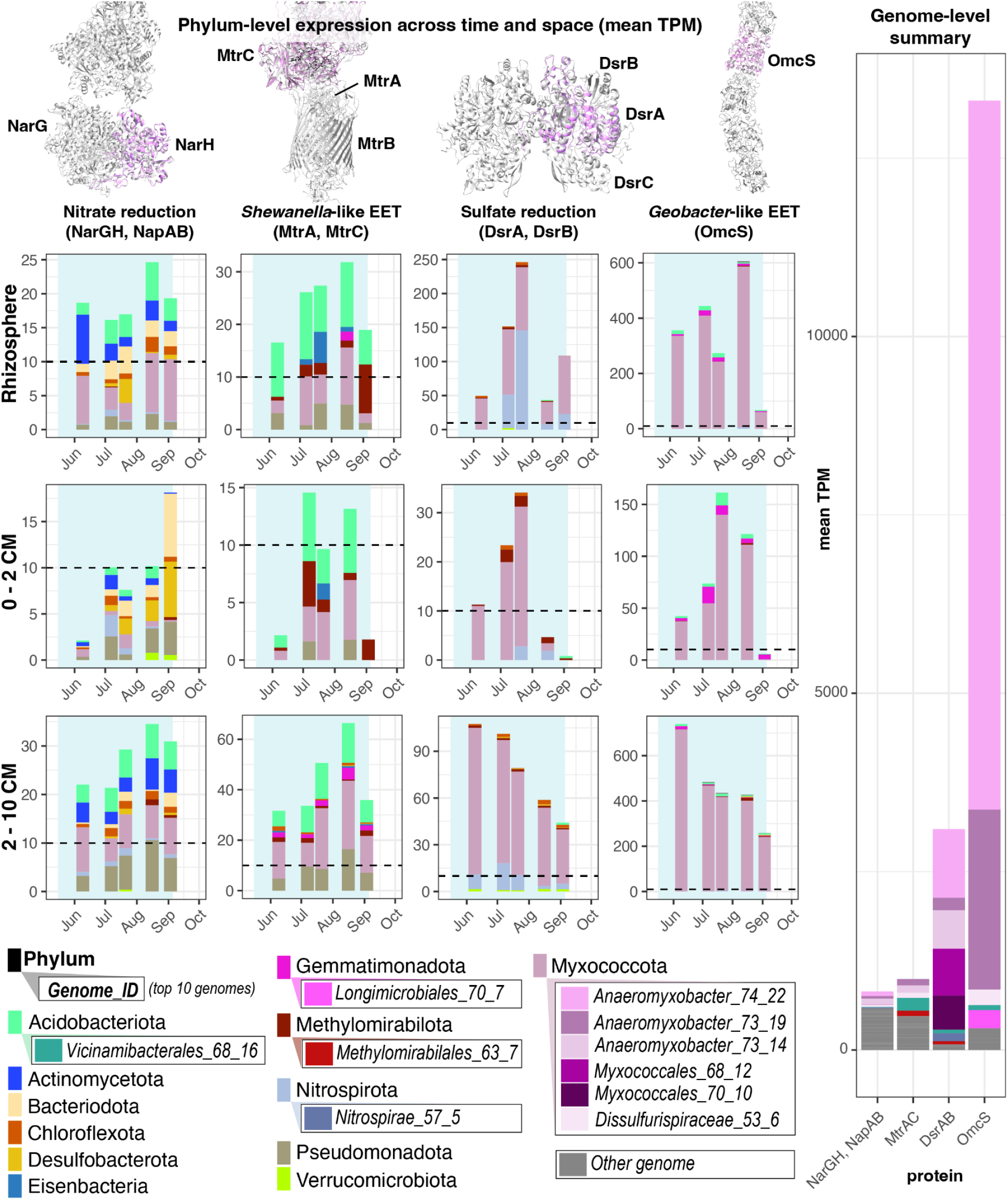
Genome-resolved composition of TEAP expression competing with methanogenesis. Left: Phylum-level comparison of dissimilatory nitrate reduction (NapA, NapB, NarG, and NarH), *Shewanella*-like EET (MtrA and MtrC), dissimilatory sulfate reduction (DsrA, DsrB), and *Geobacter*-like EET (OmcS) across soil compartments (rhizosphere, 0 - 2 cm bulk soil, 2 -10 cm bulk soil) over time. For each sample, transcripts per million (TPM) values were summed across genomes within each phylum and then averaged across replicates. A horizontal dashed line at 10 TPM highlights differences in y-axes scales. AlphaFold-predicted structures of the most highly expressed complete protein from each marker set (pink) are shown aligned to reference structures (grey; 1R27^77^, 6R2Q^78^, 2V4J^79^, and 6EF8^80^). Right: Phylum-level comparison of TEAP activity. TPM values were summed by marker category and averaged across replicates.

### Methanogen competitors are dominated by Geobacter-like EET and TEAP generalists

Overall, *Geobacter*-like EET was the major TEAP competing with methanogenesis (73% of expression). In the rhizosphere, sulfate reduction was more prominent, at times reaching over 50% of competing TEAP expression, but in the bulk soil below the oxic/anoxic interface, *Geobacter*-like EET consistently accounted for 70-80% of competing TEAP expression (Fig. S46). The most active species competing with methanogens expressed multiple TEAPs, likely reflecting the redox state of their local microenvironments. Several of the strongest TEAP-expressing genomes were phylogenetically similar Myxococcota pertaining to different cohorts. For example, 75% of *omcS* expression originated from a single Myxcoccota genome, genome 7, which also generated 31% of *dsrAB* expression and 8% of nitrate reduction marker expression (Fig. 6). The dominance of generalist taxa suggests that fluctuating rice paddy redox conditions favor microbes capable of using diverse soluble and insoluble electron acceptors.

### Methane oxidation is partitioned across oxic and anoxic soil compartments

#### pMMO-containing methanotrophs occupy both oxic and anoxic soil

Aerobic methanotrophs are considered major methane consumers in rice paddies^3^, and our genome-independent analysis recovered expression of 805 unique species-level *pmoB* clusters. We reconstructed pMMO-utilizing bacterial methanotroph genomes from both anoxic and oxic soil compartments. Two *Methylobacter* type I methanotroph genomes (121 and 124) were recovered from oxic soil and were most active and abundant in the rhizosphere and early-season bulk soil, when oxic microsites are expected to be common.

Four *Methylosinus* type II methanotroph genomes (290, 545, 571, and 589) were recovered from anoxic bulk soil. Type II methanotrophs are adapted to lower oxygen and higher methane concentrations than type I methanotrophs, but still employ the oxygen-dependent pMMO methane oxidation mechanism. Direct DNA-based replication metrics and seasonal increases in relative abundance indicate growth of these methanotrophs in anoxic bulk soil (Data S9; see Methods). These populations may operate within oxic microsites or, under O₂ limitation, use alternative terminal electron acceptors to dissipate reducing equivalents generated during methane oxidation. This possibility is supported by reports of pMMO-encoding methanotrophs in apparently anoxic freshwater habitats^54^, as well as enrichment studies showing methane-dependent nitrate reduction and iron oxide reduction by conventionally aerobic methanotrophs^55,56^.

#### Methanoperedens populations express diverse extracellular electron-transfer machinery

We recovered five genomes of anaerobic methane oxidizing archaea from deep bulk soil cohorts (steelblue, pink, and blue). Anaerobic methane oxidation ramped up in mid-July, and was dominated by *Methanoperedens nitroreducens*, which couples methane oxidation to the reduction of alternative electron acceptors including nitrate, iron, manganese^57^ and possibly humic acids^58,59^.

In a Chinese rice paddy, anaerobic oxidation of methane (AOM) was primarily linked to nitrite and nitrate reduction, but also to diverse electron acceptors whose relative importance varied across soil horizons and time^15^. Here, gene expression data suggest that *M. nitroreducens* populations are similarly metabolically versatile. Although RNA yields from the deep soil that *M. nitroreducens* primarily inhabits were insufficient for sequencing, we detected *mcr* expression from three *M. nitroreducens* genomes (167, 593, and 666) in 2-10 cm bulk soil. We detected expression of multiple distinct multi-heme cytochromes (MHCs) pertaining to the same genomes. We did not detect transcript evidence for canonical nitrate-dependent AOM, but did detect expression of two pentaheme cytochromes most similar to formate-dependent nitrite reductase (68.0% and 73.2% nucleotide identity). Four additional expressed MHCs were predicted to bind 8-25 heme-c ligands, localize extracellularly, and have homology to iron-reduction systems like OmcZ, DFE_0462, and DFE_0449 (Table S1). Together, these data suggest that *M. nitroreducens* may use flexible extracellular electron-transfer pathways to couple methane oxidation to locally available electron acceptors, including metal oxides, nitrite, sulfite^60^, or other redox-active substrates.

#### Nitrogen-cycling populations may regenerate electron acceptors for anaerobic methanotrophy

In other rice paddy systems, *M. nitroreducens* is associated with NC10/Methylomirabilota bacteria, which can use nitrite generated by *M. nitroreducens* for intra-aerobic methane oxidation via nitric oxide dismutase (NOD)^61^. We did not detect NC10/Methylomirabilota bacteria, but identified four putative NODs outside this lineage (Fig. S47), suggesting that intra-aerobic metabolism may also occur in anaerobic microbes with other metabolic strategies^62^.

To further resolve community members potentially supporting anaerobic methanotrophy, we deeply sequenced a 20–30 cm bulk soil sample with high *M. nitroreducens* abundance using PacBio HiFi sequencing. We recovered a putatively circular *M. nitroreducens* genome (160_C_Methanosarcinia), along with 26 putatively circular bacterial and archaeal genomes encoding nitrogen-cycling functions that may influence nitrate availability to *M. nitroreducens* (Extended Data Fig. 5; Data S10).

*M. nitroreducens* co-occurred with low-oxygen-adapted Nitrososphaeraceae ammonia oxidizers, whose genomes were also detected only below 10 cm in bulk soil (Extended Data Fig. 4). These Nitrososphaeraceae may oxidize the ammonia supplied during fertilization (Table S2). If the *M. nitroreducens* nitrite-reductase-like pentaheme cytochromes discussed above reduce the resulting nitrite to ammonium, this co-occurrence raises the possibility of a cryptic ammonium–nitrite cycle (Extended Data Fig. 4). In this scenario, methane would provide the reducing power, while nitrogen species would act as recyclable intermediates rather than terminal energy sources. Alternatively, Nitrososphaeraceae-derived nitrite could be oxidized to nitrate by co-occurring nitrite oxidoreductase (*nxrAB*)-encoding Acidobacteriota, Chloroflexota, and Planctomycetota, providing a potential source of nitrate for *M. nitroreducens* (Extended Data Fig. 5).

Putatively circular genomes allowed us to assess not only which nitrogen transformations were encoded by individual community members, but also which steps were absent from each genome. This genome-resolved inference builds on prior work in complex subsurface communities showing that sequential redox pathways are often partitioned across coexisting organisms^63,64^. Together, these data suggest that deep-soil nitrogen cycling depends on metabolic handoffs among multiple interacting populations, with nitrate availability to anaerobic methanotrophs shaped by a broader network of ammonia- and nitrite-oxidizing organisms.

## Conclusions

Our genome-resolved approach showed that methane cycling is embedded within environmentally-structured microbial cohorts rather than controlled by isolated taxa. These cohorts were associated with distinct drivers, including soil depth, pH, organic matter availability, flooding state, and plant-associated compartments. Within several cohorts, genome ranks were conserved across chemically and physically distinct soil compartments, consistent with stable ecological associations and potential functional coupling among cohort members. This structure was especially pronounced in methanogen-containing cohorts, suggesting that methane-producing populations are embedded in particularly stable networks of interacting organisms.

Cohort-level genome content indicated broad capacity for terminal electron-accepting processes across the redox spectrum, but metatranscriptomic data showed that TEAP activity was concentrated in a subset of cohorts. By linking expressed TEAPs to genomes, we found that phylogenetically similar, metabolically flexible Myxococcota from different cohorts were the most active competitors for methanogenic substrates. Thus, diversion of substrates away from methanogenesis in rice soils is not simply governed by the presence of alternative electron acceptors, but by which microbial cohorts are positioned to exploit them as redox conditions shift.

By sampling across soil depth and over the growing season, we found that methanogenic activity was temporally partitioned among soil compartments. Subsurface bulk soil showed evidence of an earlier methanogenic peak than the rhizosphere and surface bulk soil, suggesting that it becomes methanogenic sooner because it is less directly affected by oxygenation and seasonal replenishment of high-redox-potential electron acceptors. This depth-specific timing implies that methane mitigation strategies should target not only late-season rhizosphere-associated methanogenesis, but also early-season methane production from subsurface bulk soil. Practices that alter deep-soil redox dynamics, such as modified flooding regimes, root-zone oxygenation, or rotations with deeper-rooted crops, represent testable strategies for disrupting this early methane source.

Cohort analysis generated additional hypotheses for manipulating methane cycling through management of microbial community interactions. For example, *M. nitroreducens* co-occurred with low-oxygen-adapted ammonia-oxidizing Nitrososphaeraceae archaea in deep bulk soil, suggesting that nitrogen transformations may influence the availability of electron acceptors supporting anaerobic methane oxidation. This raises the testable possibility that targeted nitrogen delivery to the depths where *M. nitroreducens* is most abundant (20-30 cm) could stimulate methane oxidation to mitigate methane emissions that apparently originate from deep soil early in the growing season. More broadly, our results show that microbial functions relevant to greenhouse gas emissions cannot always be predicted from individual genomes or taxa alone. Instead, realized metabolism depends on the local ecological context, including cohorts that supply substrates, remove products, compete for electron donors, or regenerate electron acceptors. Together, these results establish a community-scale framework for understanding methane cycling in rice agriculture.

## Materials and Methods

The soil microbiome of M-206 rice was observed at the Rice Experiment Station (Biggs, CA; 39°27’30.07” N, -121°44’20.06” W) throughout the 2021 growing season (May – September), where the soil is characterized as Esquon-Neerdobe complex (Simmonds et al., 2015) and averaged 45.25% clay, 27.5% silt and 27.25% sand (Data S1). The experimental basin has been in rice for decades. In the previous year, after rice was harvested, the rice straw was chopped and incorporated into the soil and the field flooded as is standard practice^65^.

### Field treatments

Standard management practices for water-seeded systems were employed for the test basin (∼0.21 ha). Tillage to ∼15 cm began in late April of 2021 to prepare a seedbed. Nitrogen fertilizer was applied at a rate of 168 kg/ha as aqueous-ammonia injected 5-10 cm below the soil surface on 11 May. Rice was seeded at a rate of 168 kg/ha on 13 May 2021 and flooded by 16 May 2021. The field remained flooded until it was drained late in the season (5 September 2021) in preparation for harvest. On June 10, 45 kg/ha P_2_O_5_, 39 kg/ha K_2_O, and 17 kg/ha Zn were applied to ensure these nutrients did not limit plant growth. Pests were controlled with the application of insecticide (0.28 L/ha Lambda-Cy AG on 23 May 2021), herbicide (8.4 kg/ha BUTTE® on 25 May 2021, 14 L/ha SuperWHAM!® CA and 0.29 L/ha Grandstand® CA on 25 June 2021), and fungicide (0.88 L/ha Quadris® on 4 Aug 2021; Table S2).

### Soil sampling

At each of 7 timepoints, replicate samples were collected along a boardwalk from 20’-35’ into the field. Rhizosphere metagenomes and metatranscriptomes were collected in triplicate on the left side of the boardwalk. Replicates were collected moving inward, with replicate 1 closest to the perimeter of the paddy. On the right side of the boardwalk, a tarp was used to prevent seeding and expose a bulk soil control. Triplicate bulk soil cores (1” diameter) were taken directly across from sampled plants, and divided into 2-10 cm, 10-20 cm, 20-30 cm, and 50-60 cm sections. Oxic samples were also taken at the location of each core by collecting the top ∼2 cm of sediment with a trowel. Three cores from each replicate were dried and used for total nitrogen, total carbon, Zn, Mn, Fe, Cu, and particle size. One core was kept on wet ice and used to measure pH; nitrate and ammonium (via KCl extraction) and gravimetric soil moisture. At approximately 11:00 a.m., one core per replicate was sectioned, rolled flat, and flash-frozen in liquid nitrogen within two minutes for nucleic acid extractions. At approximately 12:00 p.m., rhizosphere soils were collected for nucleic acid extraction by gently uprooting a plant, shaking off excess soil, trimming a section of interior root with sterile scissors, and submerging it in 8 mL Powerbead Solution, and shaking vigorously with sterile forceps. The root was then discarded, and the tube was flash-frozen in liquid nitrogen within 2 minutes. All nucleic acid samples were transported on dry ice and stored at -80°C until extraction.

### Trace gas sampling

For May and June timepoints, a Picarro G2508 gas analyzer was used in tandem with an Eosense eosMX multiplexer to measure CH_4_, N_2_O, CO_2_, and NH_3_ concentrations. With the exception of the pre-seeding May timepoint, flux measurements were collected using a transparent chamber to allow for the photosynthesis of rice plants. The transparent chamber was removed following each measurement cycle to allow for the air exchange and prevent overheating of the plants. All gas measurements were done in a closed-loop system where headspace air was continuously recirculated. The multiplexer returned flux measurements calculated at ∼10-minute intervals in nmol/m^2^/s, which was converted to flux in grams of C or N per hectare per day.

For September and October timepoints, no multiplexer was used, so gas concentrations (ppm) over time were reported by the G2508 Picarro. These values were converted to grams of C or N emitted/volume using the ideal gas law and ratio of molar mass of C or N to the total weight of the molecule measured (e.g. 12.01/16.04 for CH_4_). Flux was then calculated by fitting a linear regression to these values over time. The slope of the best fit line was then multiplied by the volume of the chamber head. For field data, this value was normalized by the surface area of the chamber and converted to a per-hectare flux. Data points to be included in each regression were manually curated to use a stretch of time (typically 5 to 10-minute time windows) that started once gas from previous measurements were cleared from the Picarro intake line and excluded artificial spikes in methane from ebullition caused by physical disturbance.

### Gravimetric soil moisture

Within 24h - 48 h of sampling, ∼10g of soil was weighed to 3 decimal places on a pre-weighed aluminum tin. Samples were baked at ∼105-110 °C for at least 2 days until additional drying did not alter weight, at which point weight was recorded to 3 decimal places. Gravimetric moisture content was calculated as (weight of fresh soil – weight of dry soil)/ weight of dry soil.

### KCl extractable nitrogen

In advance of sampling, one funnel corresponding to each sample was acid washed and DI rinsed. One WhatmanTM grade 1, 125 mm filter circle corresponding to each sample was folded into quarters and placed in the DI-rinsed funnel of a vacuum flask, and rinsed first with 50 mL DI water, then with 50 mL autoclaved 2M KCl. Filters were then stored at 4°C until needed. A specimen “S cup” (to be used for shaking) was weighed to three decimal places, filled with 50 mL of autoclaved 2M KCl, weighed again, and stored at 4°C. On the day of collection, ∼ 5g of soil that had been kept on wet ice was added to each sample’s S cup and shaken at 200 rpm for 1 hour. S cups were then allowed to settle on the benchtop for 20 minutes, before being poured through prepared Whatman filters into a fresh specimen cup and stored at -20°C. Samples were then shipped on dry ice to the UC Davis analytical lab, where NO2-N, NO3-N and NH4-N were measured by flow injection analyzer method^66–68^ and reported in mg N/L. These values were then blank-corrected by subtracting the average value of blanks collected for each time point, and converted to ug N/g dry soil, considering the volume of 2M KCl used and gravimetric moisture content of soils.

### Soil pH

Within 24h - 48 h of sampling, 5 g from each sample (stored on wet ice) was partitioned into a 50 mL Falcon tube (Thermo Fisher Scientific) along with 10 mL DI water for pH measurement. In order to dissolve the clay, samples were vortexed for 5 minutes, shaken at 200 rpm for 30 minutes, then uncapped and allowed to equilibrate with the atmosphere for 30 minutes. pH was measured with an Oakton instruments 110 series pH meter calibrated with a pH 7 buffer solution (Fisher Chemical), and pH 4.006 and 10.012 InLab solution sachets (Mettler Toledo).

### Nucleic acid extraction and sequencing

Frozen soil cores were crushed into small fragments on dry ice using an ELIMINase-treated (DeconLabs) aluminum-foil wrapped hammer in order to homogenize sections. RNA and DNA were co-extracted from 0-2 cm sections, 2-10 cm sections, and rhizosphere soil using the RNeasy PowerSoil Total RNA Kit (Qiagen) in tandem with the RNeasy PowerSoil DNA Elution Kit (Qiagen). For rhizosphere extractions, collection tubes with powerbead solution were weighed before and after sample collection in order to estimate mass of soil sampled. Immediately upon thawing, the rhizosphere slurry was homogenized and 3.6 mL of solution was aliquoted into a powerbead tube, and extraction proceeded with the addition of solutions SR1 and IRS.

For deeper 10-20 cm and deeper samples, DNA was extracted from ∼ 6 grams of soil the DNeasy PowerMax Soil Kit (Qiagen) with the modification that the final C6 elution buffer was passed twice through the DNA column and concentrated via ethanol precipitation.

Metagenomic libraries were prepared with ∼600–800 bp inserts using the KAPA HyperPrep Kit with PCR amplification (Roche) and sequenced across seven lanes on an Illumina NovaSeq 6000 SP flow cell with 250 bp paired-end reads at Maryland Genomics.

Stranded metatranscriptomic libraries were also prepared with KAPA RNA HyperPrep Kits (Roche). First, RNA was first DNase-treated with the Zymo Quick-RNA kit. Next, 0.2 ng of ArrayControl RNA standards (ThermoFisher) were spiked into 300 ng sample RNA. The standard mixture consisted of ArrayControl spike #1 (750 nt) and spike #8 (2000 nt) combined at a ratio of 2.66:1 in order to have roughly equal concentrations after fragmentation. Finally, ribosomal RNA depletion was performed with RiboCop META (LEXOGEN). RNA Libraries were pooled and sequenced on one NovaSeq S4 150PE lane at QB3 Genomics (UC Berkeley).

### Genomic data curation and assembly

Illumina PE250 reads were curated using the ggkbase metagenomic data preparation pipeline (https://ggkbase-help.berkeley.edu/overview/data-preparation-metagenome/), including adaptor and illumina trace contaminant removal with BBTools, and quality trimming with Sickle. Assembly was performed with metaspades, followed by gene prediction with prodigal in metagenome mode and annotation with usearch against KEGG, UniRef100, and UniProt databases for scaffolds > 1 kb.

### rpS3 marker gene analysis

Bacterial and archaeal rpS3 sequences were identified from predicted metagenomic proteins using HMMER hmmsearch with the Pfam ribosomal protein S3 C-terminal domain model PF00189.22 and trusted cutoffs. Matching genes were retrieved as nucleotide sequences from the corresponding predicted gene fasta file, because nucleotide-level clustering preserves higher strain- and species-level resolution. rpS3 nucleotide sequences were clustered at 99% identity with vsearch --cluster_fast, and cluster centroids were used as representative sequences. For each rpS3 cluster, the longest associated scaffold was selected and used for read recruitment. Metagenomic reads were mapped to these scaffolds, and scaffold coverage was calculated with CoverM using a minimum read identity of 99%; length-normalized mean coverage was retained only for scaffolds with at least 50% covered fraction. Taxonomic annotations of rpS3 centroids were assigned by BLASTX against NCBI NR, retaining the top hit, and taxonomic lineages were recovered from NCBI taxon identifiers using TaxonKit. To assess bin representation, rpS3-containing scaffolds were linked to DAStool bins using scaffold-to-bin tables generated from the metagenomic binning workflow.

Beta diversity was calculated from the unbinned rpS3 cluster abundance table using Bray–Curtis dissimilarities. NMDS ordination was performed with vegan::metaMDS using trymax = 200, and site scores from this solution were plotted in Fig. S5. Effects of compartment/depth, time and replicate identity on community composition were tested by PERMANOVA using adonis2 with 9,999 permutations and the model bray_dist ∼ depth_code * days_since_start + replicate. The Bray–Curtis NMDS stress was 0.11055.

### Genome-resolved metagenomics

Scaffolds were binned by sample cross-mapping with BBMap ^69^, followed by binning with concoct, maxbin2, metabat2, and vamb, then choosing the best overall bin set with dastool. The dastool bins were then dereplicated with dRep using default parameters (minimum completeness 75%, maximum contamination 25%), and the resulting 473 bins were manually curated to remove obvious taxonomic, GC, and coverage anomalies. All bins were at least 50% complete and less than 10% contaminated after manual curation. This bin set was missing some important players (e.g. methanogens), so in order to recover bins that may be excluded by the dRep’s standard checkM1 filter and stringent completeness threshold, the original dastool bins were dereplicated again with the ‘--ignoreGenomeQuality’ flag. The resulting 1031 genomes were screened with checkM2 and bins with a minimum of 50% completeness and a maximum of 10% contamination. These 719 bins were reconciled with the original manually curated bin set using a final round of dRep with no completeness or contamination threshold, ultimately producing 204 high-quality and 525 medium quality MAGs.

Metagenomic reads were mapped with a 95% ID threshold to the final dereplicated set of 729 consensus rice-field genomes with BBMap^69^. Mean genome coverage and the fraction of the genome covered were calculated from the BAM files using CoverM ‘-m mean covered_fracion’. A covered fraction of 30% was required for a genome to be considered present in a sample.

Metatranscriptomic reads were also mapped to the dereplicated genome set at 95% identity. Genome-level transcript coverage and covered fraction were calculated from sorted BAM files using CoverM ‘-m mean covered_fraction’ with ‘--min-covered-fraction 0’, because transcript coverage was expected to be sparse across genome scaffolds. To quantify gene-level expression, ORFs were predicted on the concatenated consensus-bin contigs with prodigal in metagenomic mode, and strand-specific counts for each CDS were calculated from RNA BAM files using featureCounts^70^ with paired-end reads. Because the RNA libraries were reverse-stranded, ‘-s 2’ was used for sense expression. Sense expression was normalized as transcripts per million (TPM) for downstream analyses.

### Cohort determination and functional enrichment

We clustered the dereplicated genome set based on co-occurrence patterns using Weighted Group Correlation Network Analysis (WGCNA) with a soft-power threshold of 7, a module eigengene dissimilarity threshold of 0.35, and a minimum module size of 5 genomes^71^. We used a signed adjacency matrix and required that genomes in each module had coverage values that positively correlated with the overall cohort eigenvalues (average abundance profile). We tested pairwise Spearman correlations between each cohort’s eigenvalues and soil metadata variables using the corr.test() function of the R psych^72^ package. P-values were FDR adjusted using p.adjust().

KEGG ortholog (KO) enrichment within genome cohorts was tested using DRAM-annotated KOs. For each KO detected in at least five genomes, genomes were scored by KO presence/absence and assigned to WGCNA cohorts. Fisher’s exact tests were then used to compare the frequency of each KO in a given cohort against all other genomes. P values were corrected across tests using the Benjamini–Hochberg procedure, and KOs with FDR-adjusted P < 0.05 and odds ratio > 1 were considered enriched.

### Correlation analysis of within-cohort genome abundances

Library-normalized metagenomic coverage values were averaged across replicate samples for each genome × depth × date combination, assigned to previously defined WGCNA cohorts, and expressed as each genome’s percentage contribution to total cohort abundance. To quantify conservation of within-cohort abundance structure across soil compartments, genomes were ranked within each cohort, date and depth by percentage contribution. Samples with fewer than five ranked genomes were excluded. For each cohort and pair of compartments, matched genome × date observations were pooled and compared using Spearman rank correlation. Null distributions were generated by permuting genome identities within each cohort, date and compartment pair, preserving cohort size and marginal rank distributions while disrupting the correspondence of individual genomes between compartments.

To assess whether rank stability declined with distance from each cohort’s dominant bulk-soil habitat, an anchor depth was defined as the bulk-soil compartment with the highest mean cohort abundance across sampling dates. Rhizosphere samples were excluded from anchor assignment. Analyses were restricted to comparisons between the anchor and other bulk-soil depths, with distance defined as the number of bulk-soil compartments separating the two depths. For each matched genome × date observation, rank displacement was calculated as the absolute rank difference between the anchor and comparison depth, normalized by the mean number of ranked genomes in the two samples: |rankcomparison − rankanchor| / ((nanchor + ncomparison) / 2). For each cohort × distance combination, the median normalized rank displacement was reported. Comparisons were retained only when both cohort-date-depth samples contained at least five ranked genomes, the summarized comparison contained at least five matched genome × date observations, and the non-anchor depth had mean cohort abundance ≥0.015.

### Identification of hydrogenases and terminal electron accepting processes

[NiFe]-, [FeFe]- and [Fe]-hydrogenases were identified using HydDB HMMs^48^ with a trusted-cut threshold. Alignments were built for each hydrogenase type using MAFFT (‘--autò). Alignments were trimmed with TrimAL in automatic trimming mode (‘-gappyout’) before being manually curated in Geneious. For [NiFe]-hydrogenases, sequences without the characteristic N- and C-terminal CxxC motifs^73^ were removed, and known HydDB references^48^ were used to distinguish hydrogen-evolving, hydrogen-consuming, and bidirectional classes as previously described^74^. Maximum-likelihood phylogenies were built using IQtree and visualized using iTOL (https://itol.embl.de/). Branch support was determined with ultrafast bootstrap approximation using 1,000 replicates.

Methanogens and anaerobic methanotrophs were identified using *mcrA* as a marker gene, while aerobic methanotrophs were identified using *pmoB*. Dissimilatory nitrate reduction potential was inferred from the presence of either periplasmic nitrate reductase (*napAB*; TIGR01706, PF03892) or membrane-bound nitrate reductase (*narGH*; TIGR01580, TIGR01660). Dissimilatory sulfate reduction was identified based on the presence of reductive *dsrA* and *dsrB* genes annotated using DiSCo.

Multiheme c-type cytochromes (MHCs) were identified in our 95% ANI dereplicated genome set based on the presence of repeated Cys-X₂-Cys-His (CXXCH) heme-binding motifs in protein sequences, which are required for covalent heme c attachment and are diagnostic of c-type cytochromes. To identify *Geobacter*- and *Shewanella*-like EET, we clustered all MHCs with either 6 or 10 MHCs (n= 1230) using mmseqs easy-linclust with the parameters ‘--min-seq-id 0.3 --cov-mode 1 -c 0.2’, then folded each clusters representative sequence (repseq; n= 448) using colabfold with template search enabled and 3 recycles. The best models for each repseq were annotated against alphafold swissprot and PDB using foldseek, and the best annotation with an e-value of at least 1e-10 was retained. MHCs with a representative sequence annotated as *omcS* and a final psortb prediction of extracellular were considered to be markers of *Geobacter*-like EET (n=26). Additionally, we included all genes annotated via metabolic’s built in *omcS* HMM and a final psortb prediction of extracellular, which added 8 genes, all of which had fewer than 6 heme-binding motifs but were retained after individual evaluation. Seven were partial *omcS* sequences and one was a complete sequence that appears to be a true *omcS* homolog that only binds 4 hemes (foldseek e-value of 2.96e-36 to OmcS structure 6EF8). MHCs with a representative sequence annotated as *mtrA* with a periplasmic psortb final prediction or *mtrC* with an extracellular psortb final prediction were considered to be markers of *Shewanella*-like EET (n= 157).

### Genome-independent marker gene analysis

We constructed a non-redundant protein catalogue from all predicted open reading frames (ORFs) in the paired metagenomic (minimum length: 1 kb) and metatranscriptomic assemblies. For the metatranscriptome-derived ORF set, translated sequences shorter than 100 amino acids were excluded before clustering. The final clustering input comprised 96,251,231 metagenomic proteins and 3,538,340 metatranscriptomic proteins.

Translated ORFs from the metagenomic and metatranscriptomic assemblies were clustered jointly with MMseqs2 easy-linclust using a minimum amino-acid identity of 95% and a coverage threshold of 85% (‘--min-seq-id 0.95 -c 0.85 --cov-mode 0 --cluster-mode 2’). This bidirectional coverage criterion required the aligned region to cover at least 85% of both sequences. The longest sequence in each greedy incremental cluster was retained as the representative sequence. This procedure produced 64,950,587 representative protein clusters from the 99,789,571 input proteins. Source contigs corresponding to representative proteins were then recovered from the original assemblies by parsing representative ORF identifiers to their parent contig identifiers. In total, 36,750,623 unique source contigs were retained.

Sample-level DNA abundance and RNA expression were quantified by mapping reads back to the representative-cluster source contigs. Metagenomic reads were mapped to the representative contig set with Bowtie2, and coverage was summarized with CoverM v0.7.0 using a minimum read identity of 95%. Mean coverage, covered fraction, contig length and raw read counts were calculated per representative ORF or contig; coverage summaries used a minimum covered fraction of 30%. Metatranscriptomic reads were mapped with Bowtie2 to the same representative-contig set. Because of the size of the reference, the contig set was split into four approximately equal chunks for indexing and mapping. Mapped RNA alignments were filtered to retain only reads with at least 95% nucleotide identity to the reference. Strand-specific RNA fragment counts were then assigned to CDS features with featureCounts using paired-end mode. Because the RNA libraries were reverse-stranded, ‘-s 2’ was used for sense expression. Sense expression was normalized as transcripts per million (TPM) for downstream marker-gene analyses.

Marker genes were identified directly in the 95% representative protein set. McrA candidates were detected with the Pfam MCR alpha C-terminal and N-terminal HMMs PF02249 and PF02745 using HMMER hmmsearch with curated gathering thresholds (‘--cut_nc’). The union of the two HMM result sets contained 1,553 candidate McrA representative proteins; sequences shorter than 200 amino acids were excluded, leaving 760 McrA sequences for annotation and phylogenetic curation. PmoB/AmoB candidates were detected with TIGR03079.1 using ‘--cut_nc’, yielding 1,066 candidates and 849 sequences after the 200 amino-acid length filter. Candidate marker proteins were compared against NCBI NR with BLASTP, retaining up to 5 to 10 hits per query depending on marker set. Taxonomic identifiers were converted to lineages with taxonkit after removing secondary taxon identifiers following semicolons; hits without resolved taxonomy were discarded. The best annotated BLASTP hit per query was selected by e-value, with percent identity used as an additional tie-breaker for PsbA.

For markers requiring family-level discrimination, HMM and BLASTP annotations were refined phylogenetically. Representative marker sequences and their best nr reference proteins were aligned with MAFFT, trimmed with trimAl using the ‘gappyout’ heuristic, and placed in maximum-likelihood trees. McrA and PmoB/AmoB trees were inferred with IQ-TREE using 1,000 ultrafast bootstraps; exploratory FastTree trees were also generated during curation. Tree topology, BLASTP annotations and reference lineages were used to distinguish methanogenic McrA from methanotrophic McrA, and methane monooxygenase PmoB from ammonia monooxygenase AmoB-like sequences. Curated marker identifiers were then used to query the RNA-expression SQLite database.

Genome-independent coordination among TEAP pathways was tested on the sample-by-pathway TPM matrix. Pairwise pathway correlations were calculated across the 47 samples using Spearman correlation on log1p-transformed TPM values; Pearson correlations were also recorded. P values were adjusted for multiple testing with the Benjamini-Hochberg procedure. To separate shared temporal and depth structure from residual co-expression, the log1p-transformed TPM of each pathway was additionally residualized with a linear model including sampling date and depth as covariates, and pairwise correlations were recalculated on the residuals. Marker presence was broad across samples: DsrAB, McrA, MtrAC, NapAB, NarGH and PmoB were detected in all 47 samples, and OmcS was detected in 46 samples.

To test whether different representative sequences assigned to the same marker showed coordinated expression across samples, TPM values were aggregated for each representative sequence within each date-depth-replicate sample. For each marker, log1p(TPM) vectors across the 47 samples were correlated for all pairs of representative sequences using Pearson correlation. The observed mean pairwise correlation was compared with a permutation null generated by independently permuting sample labels within each representative sequence 200 times. Empirical one-sided P values were calculated as ‘(number of permuted mean correlations >= observed mean + 1)/(number of permutations + 1)’ and adjusted across marker sets with the Benjamini-Hochberg procedure. This analysis was performed separately for individual marker proteins and for combined multi-subunit groups.

### Identification of putative nitric oxide dismutase (NOD) proteins

Putative NOD homologs were identified in predicted proteins from our dereplicated genome set using the NOD HMM from Murali et al^75^ (hmmsearch E-value 2 × 10⁻⁴⁰). Matching protein sequences were extracted with pullseq and combined with heme-copper oxidoreductase reference sequences, including published NOD references. Protein sequences were aligned with MAFFT, trimmed with trimAl using -gappyout, and phylogenetic trees were inferred with IQ-TREE using 1,000 ultrafast bootstrap replicates and 1,000 SH-aLRT replicates. Putative NODs were assigned based on clustering with reference NOD sequences rather than HMM detection alone.

### PacBio HiFi genome recovery

High molecular weight DNA was extracted in quadruplicate from sample RES_21_09_03_R3_M2 (20-30 cm bulk soil core from replicate 3 sampled on 9/3/21) using DNeasy PowerMax Soil Kit (Qiagen), replacing bead-beating (step 4) with 20 minutes in a 65°C water bath and intermittent manual mixing. The resulting DNA was sequenced across one PacBio Revio SMRT cell (HiFi/CCS mode, 24 hour movie) after PacBioHiFi library preparation with size selection at Maryland Genomics. Reads were trimmed with BBDuk ^69^ (minavgquality = 20, qtrim=rl, trimq=20), then assembled with hifiasm_meta^76^.

## Acknowledgments

Library preparation and sequencing was conducted at Maryland Genomics for DNA and QB3 genomics (UC Berkeley) for RNA. We thank the Rice Experiment Station, operated by the California Cooperative Rice Research Foundation (CCRRF), for access to the rice paddy. We thank J. R. and L. Stogsdill, E. L. Brodie, W. L. Silver, K. Y. Estera-Molina, J. Hoff, K. Creamer, S. Diamond, Z. Zhang, E. Tang, P. Penev, A. Kust, L. Law, S. Lei, and L. Miloslavich for technical assistance and sampling support. We thank K. DiMarco for high molecular weight DNA extraction; L. E. Valentin-Alvarado, A. Guitor, D. Gittins and J. West-Roberts for bioinformatic consultation; and J. A. Giammona and M. Ghotbi for helpful discussions.

## Author contributions

Conceptualization: BCK, JJK, RS, BAL, JFB

Methodology: BCK, JJK, RS, BAL, JFB

Investigation: BCK, JJK, RS, LS, BAL, JFB

Visualization: BCK

Funding acquisition: JFB

Writing: The manuscript was primarily written by BCK, with substantial input from JFB. All authors contributed to the final manuscript.

## Data and materials availability

All genomes are available on ggkbase. Illumina genomes will be available at https://ggkbase.berkeley.edu/RES-PILOT_consensus_3_22_24_DNA_drep_95ANI/organisms and pacbio genomes will be available at https://ggkbase.berkeley.edu/RES-PILOT_PB_N-cyclers/organisms upon publication. Shotgun metagenomic and metatranscriptomic reads will be available at the National Center for Biotechnology Information (NCBI) upon publication. All other data are available in the main or supplementary text.

## Supplementary Materials

Figs. S1 to S47

Table S1 to S2

Data S1 to S10

---

**Extended Data Fig. 1:**
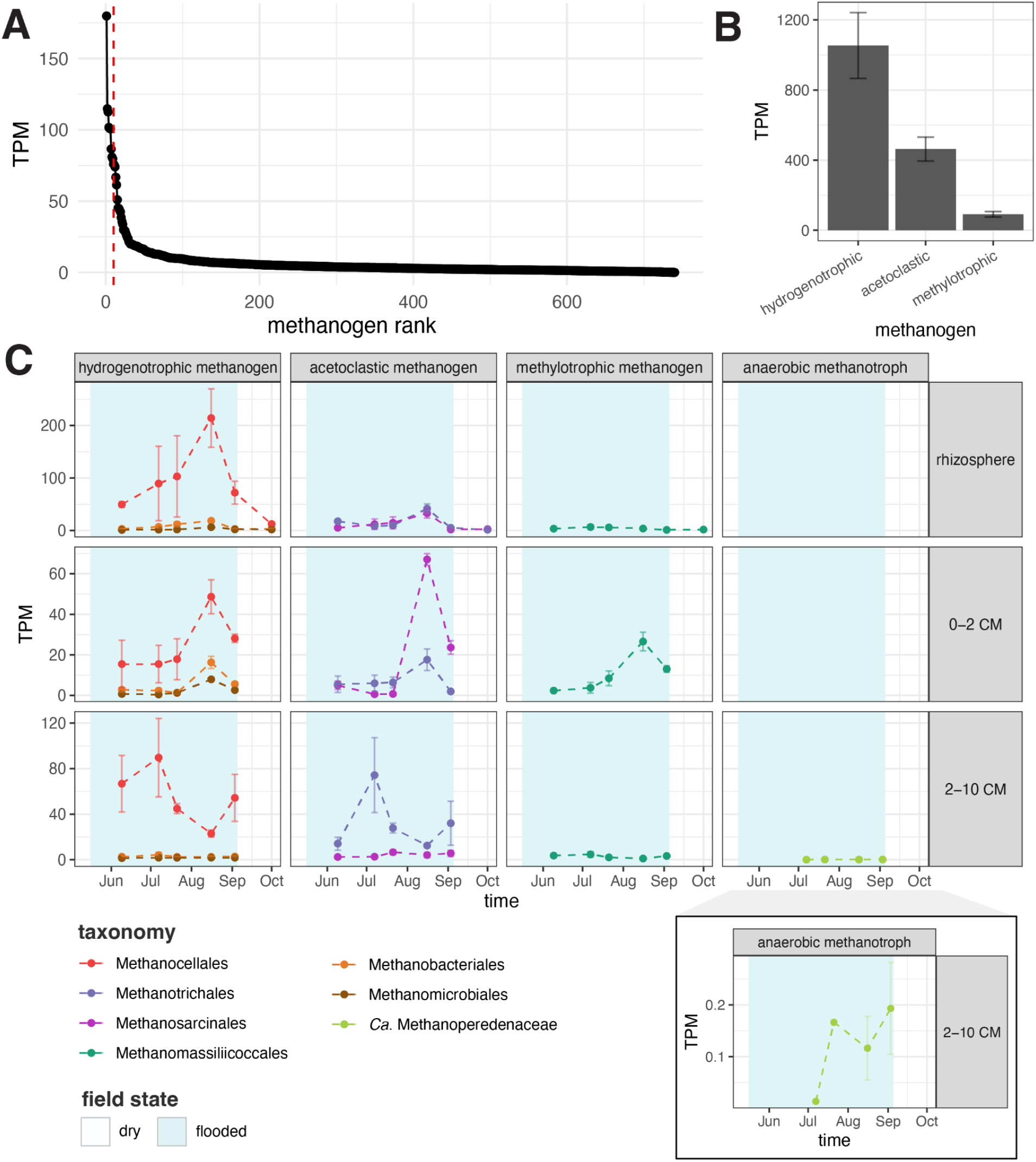
Taxonomic and metabolic diversity of *mcrA* expression. **(A)** Rank abundance of expressed species-level (95% identity) McrA protein clusters pertaining to methanogens. Expression was normalized as transcripts per million (TPM) across the genome-independent marker gene set. The dashed red line indicates top 10 clusters, which account for 21.4% of methanogenic mcrA gene expression. **(B)** Overall contributions of hydrogenotrophic, acetoclastic and methylotrophic methanogens to *mcrA* expression. **(C)** Seasonal *mcrA* expression pertaining to hydrogenotrophic, acetoclastic, and methylotrophic methanogens and anaerobic methanotrophs colored by taxonomy across soil compartments (facet rows). Note that y-axis scales differ by soil compartment. Inset plot shows seasonal anaerobic methanotroph *mcrA* expression on its own scale.

**Extended Data Fig. 2.**
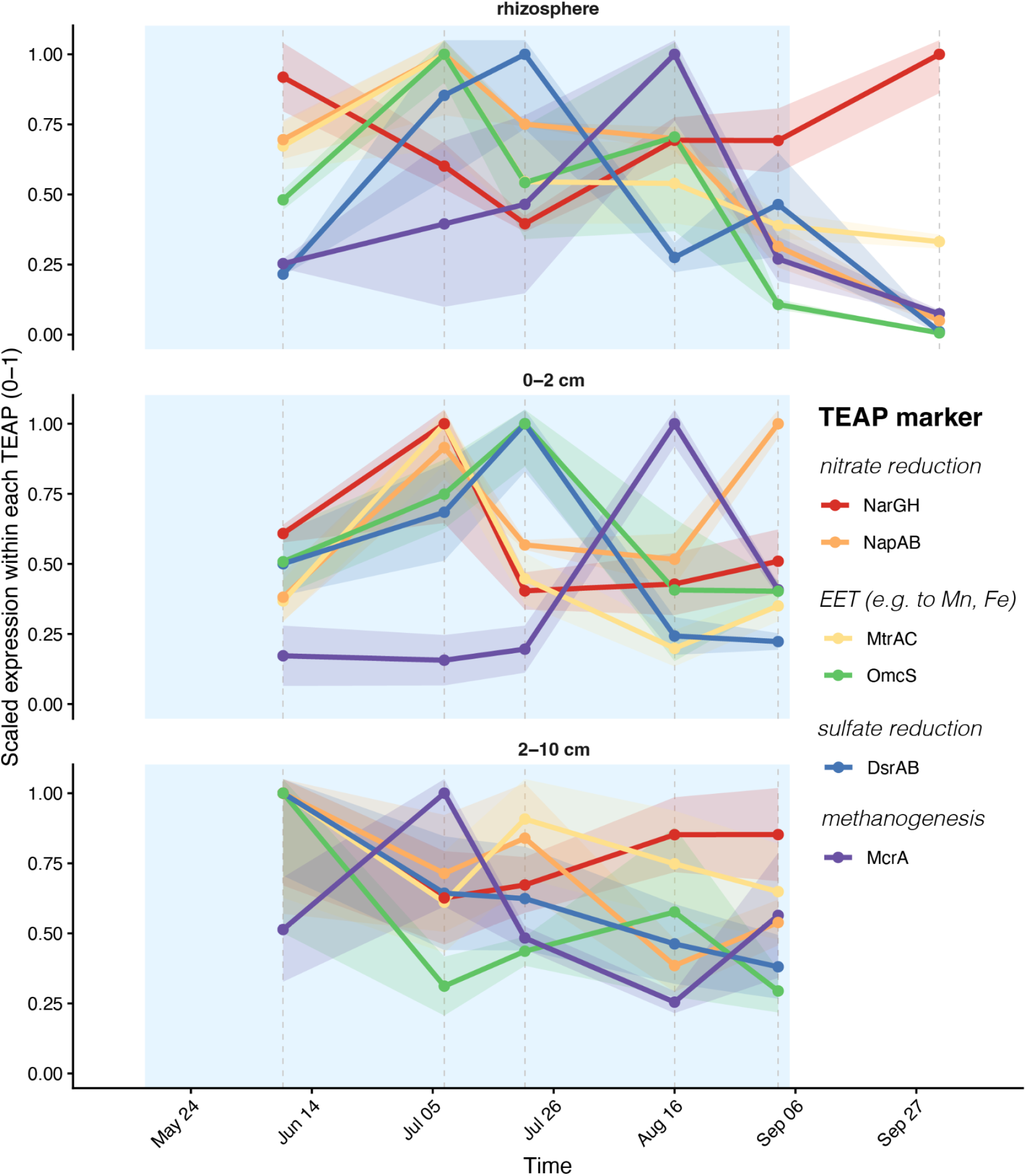
Seasonal patterns of TEAP expression across soil compartments. Genome-independent comparison of seasonal contributions of major TEAPs in the rhizosphere and bulk soil (0 -2 cm, 2-10 cm). Expression is scaled to 0-1 within each TEAP. Points represent means across triplicate samples and shaded regions represent standard error.

**Extended Data Fig. 3:**
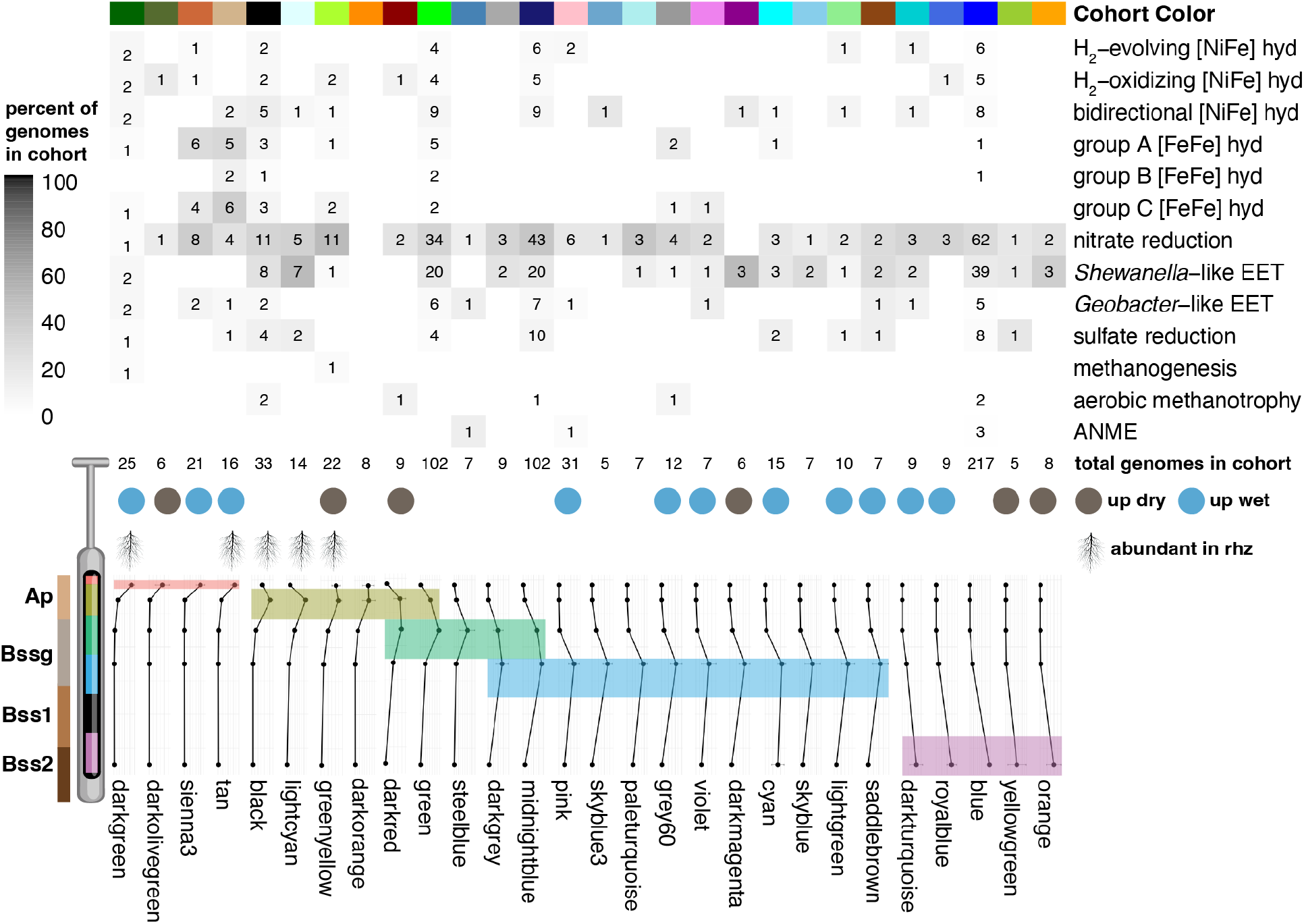
Terminal electron accepting processes (TEAPs) are broadly distributed across genome cohorts. Distribution of hydrogenases and TEAP-associated metabolic functions across microbial genome cohorts. Tile shading indicates the percentage of genomes within each cohort encoding a given function, with numbers indicating the total number of genomes encoding that function. Rows represent metabolic functions and columns represent genome cohorts. Below the heatmap, rows indicate the total number of genomes per cohort, cohorts enriched during dry (brown) versus flooded (blue) time points, rhizosphere-associated cohorts (root icon), and the average bulk soil abundance profile across depth. Cohorts are ordered by increasing depth of maximum abundance, with colors indicating the depth interval at which each cohort is most abundant. Finally, the average bulk soil coverage profile across depth is given for each module. Modules are ordered by increasing depth, and the depth at which they are most abundant is highlighted (red= 0 - 2 cm, yellow = 2 - 10 cm, green = 10 - 20 cm, blue = 20 - 30 cm, purple = 50 - 60 cm). A schematic of Esquon clay soil horizons is shown for reference: A schematic of Esquon clay soil horizons is shown for reference (Ap, plowed A horizon; Bssg, gleyed B horizon with shrink–swell slickensides; Bss1 and Bss2, Bss horizons derived from distinct parent materials).

**Extended Data Fig. 4:**
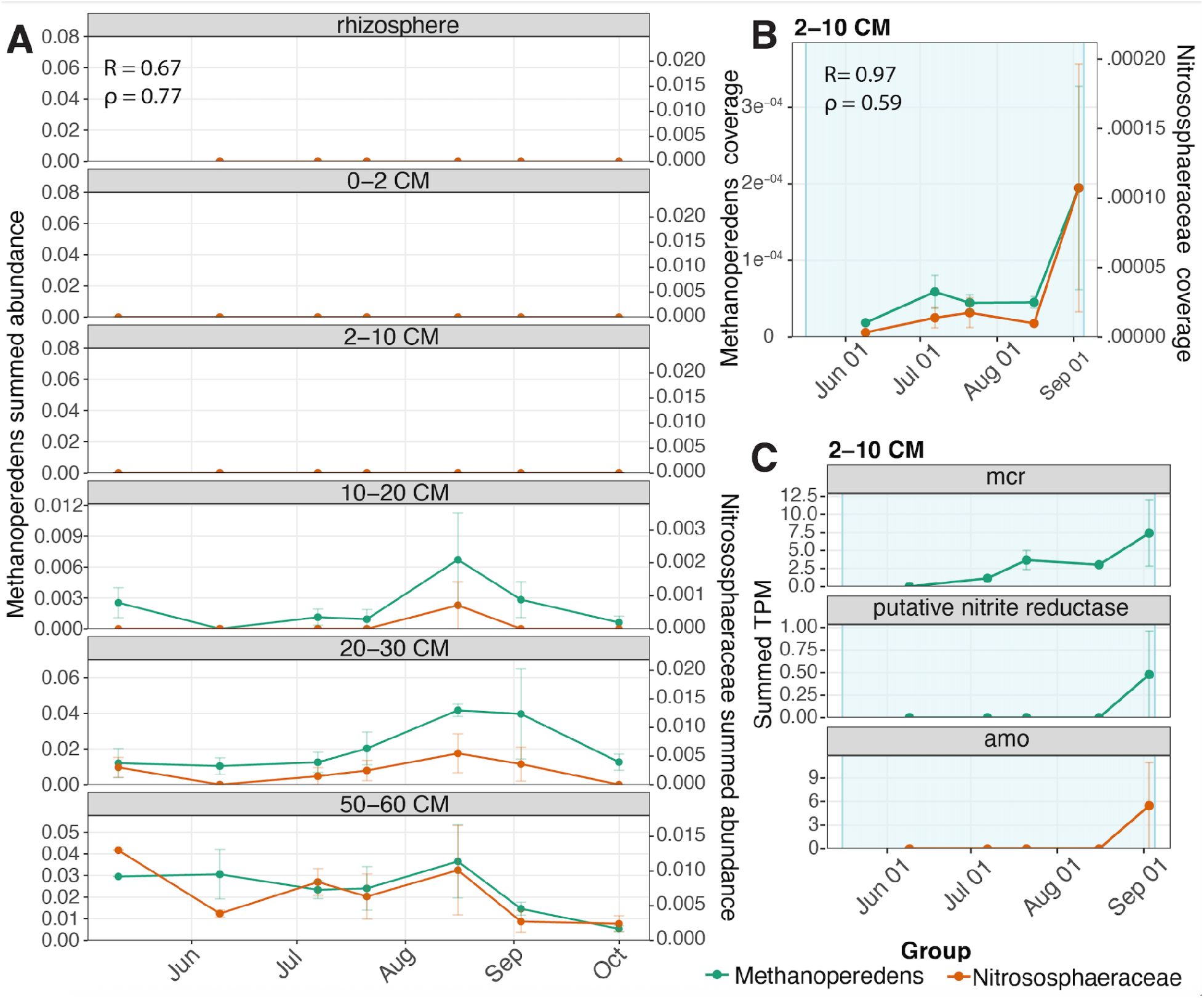
Co-occurrence and coordinated expression of Methanoperedens and Nitrososphaeraceae across rice soil compartments. **(A)** Summed genome abundance profiles of Methanoperedens (green, left y-axis) and select Nitrososphaeraceae (orange, right y-axis) across depth intervals over time, based on mean library-normalized metagenomic coverage across replicate samples. Depths are shown as separate facets. **(B)** Whole-genome RNA coverage from the deepest compartment from which we obtained RNA (2–10 cm). Points and error bars show mean ± SE across replicate samples. **(C)** Summed metatranscriptomic expression in the 2–10 cm interval for Methanoperedens mcr, a putative formate-dependent nitrite reductase, and Nitrososphaeraceae amo, shown as mean ± SE across replicate samples. Blue shading indicates the flooded period. Pearson’s R and Spearman’s ρ are given as correlation metrics. The figure includes Methanoperedensgenomes 167, 446, 447, 593, and 666 and Nitrososphaeraceae genomes 352, 402, and 696. The putative formate-dependent nitrite reductase signal represents expression of gene IDs RES_21_05_11_R0_33-63CM_MG_783_22 and RES_21_09_03_R3_M2_MG_4091_1.

**Extended Data Fig. 5:**
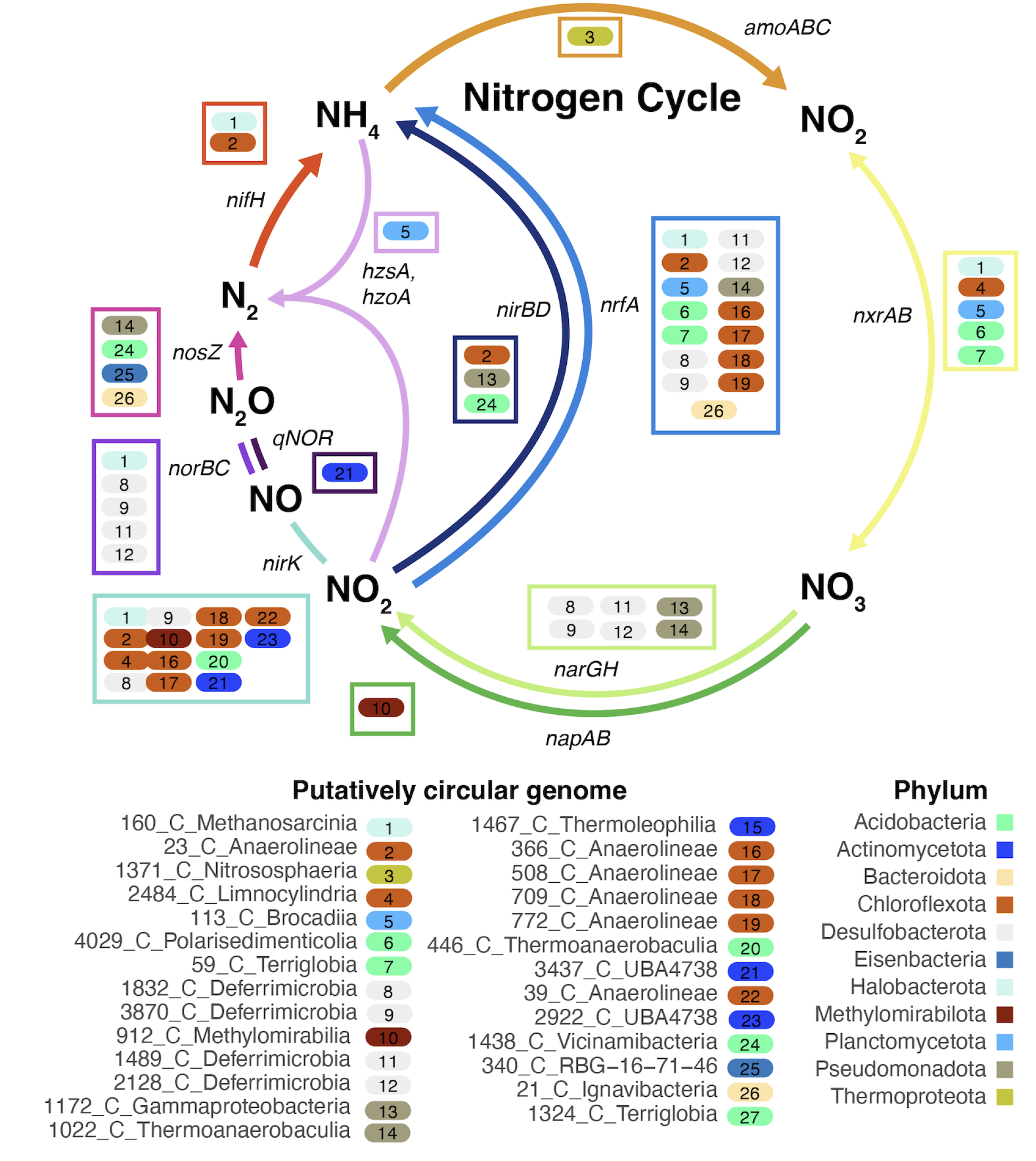
Nitrogen cycling genes encoded by putatively circular genomes. A schematic of the nitrogen cycle overlaid with genomes that encode markers for each component. Genomes are shown in numbered colored ovals that correspond to genome name and classification key.

