## Supplementary Materials for "Rice soil methane emissions are modulated by microbial cohorts that shape redox succession over the growing season"

#### Supplementary Figures

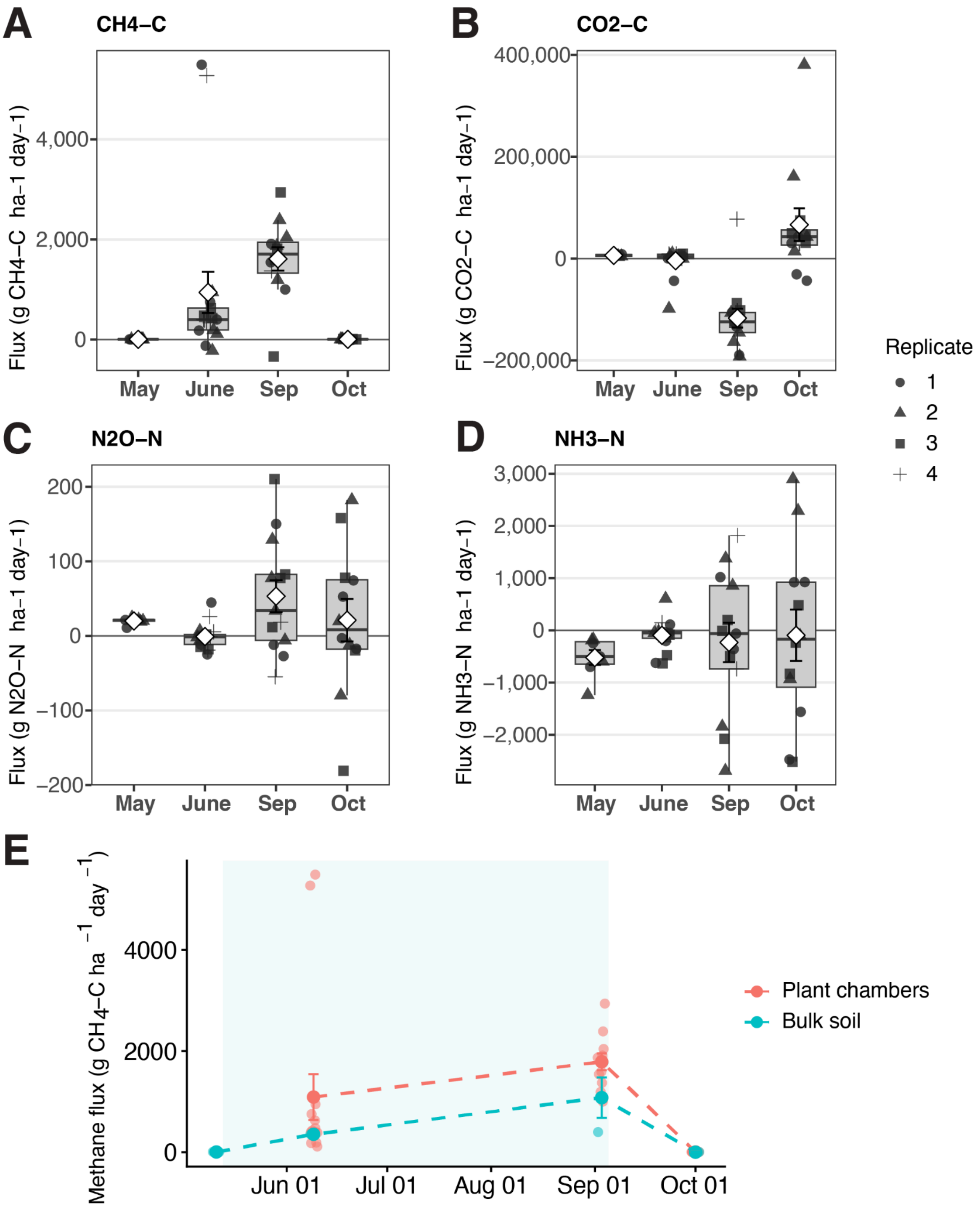


**Fig. S1. Seasonal trace gas measurements.** Seasonal fluxes of $CH_4$, $CO_2$, $N_2O$, and $NH_3$ (**A-D**) measured with a Picarro gas analyzer using a chamber placed over rice plants, when present. (**E**) Comparison of $CH_4$ fluxes measured over plants vs bulk soil. Blue shading indicates the flooded period.

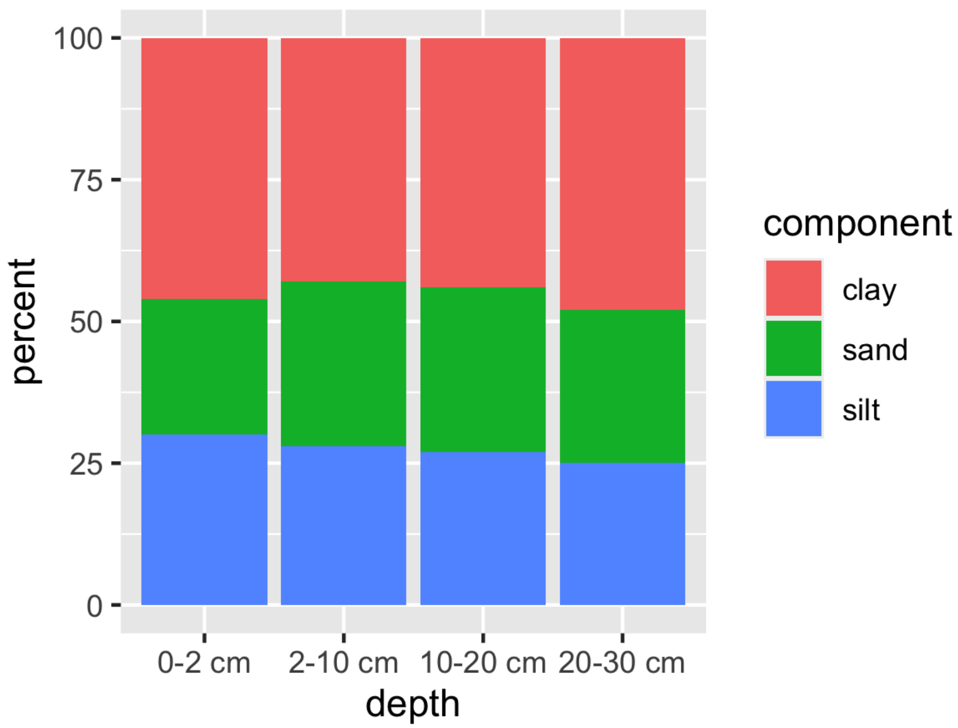


**Fig. S2. Relative contribution of clay, silt, and sand to soils from the Rice Experiment Station field site by depth, determined by hydrometer particle-size analysis.** Clay (< 2 µm), silt (2–50 µm), and sand (50–2000 µm) fractions are shown.

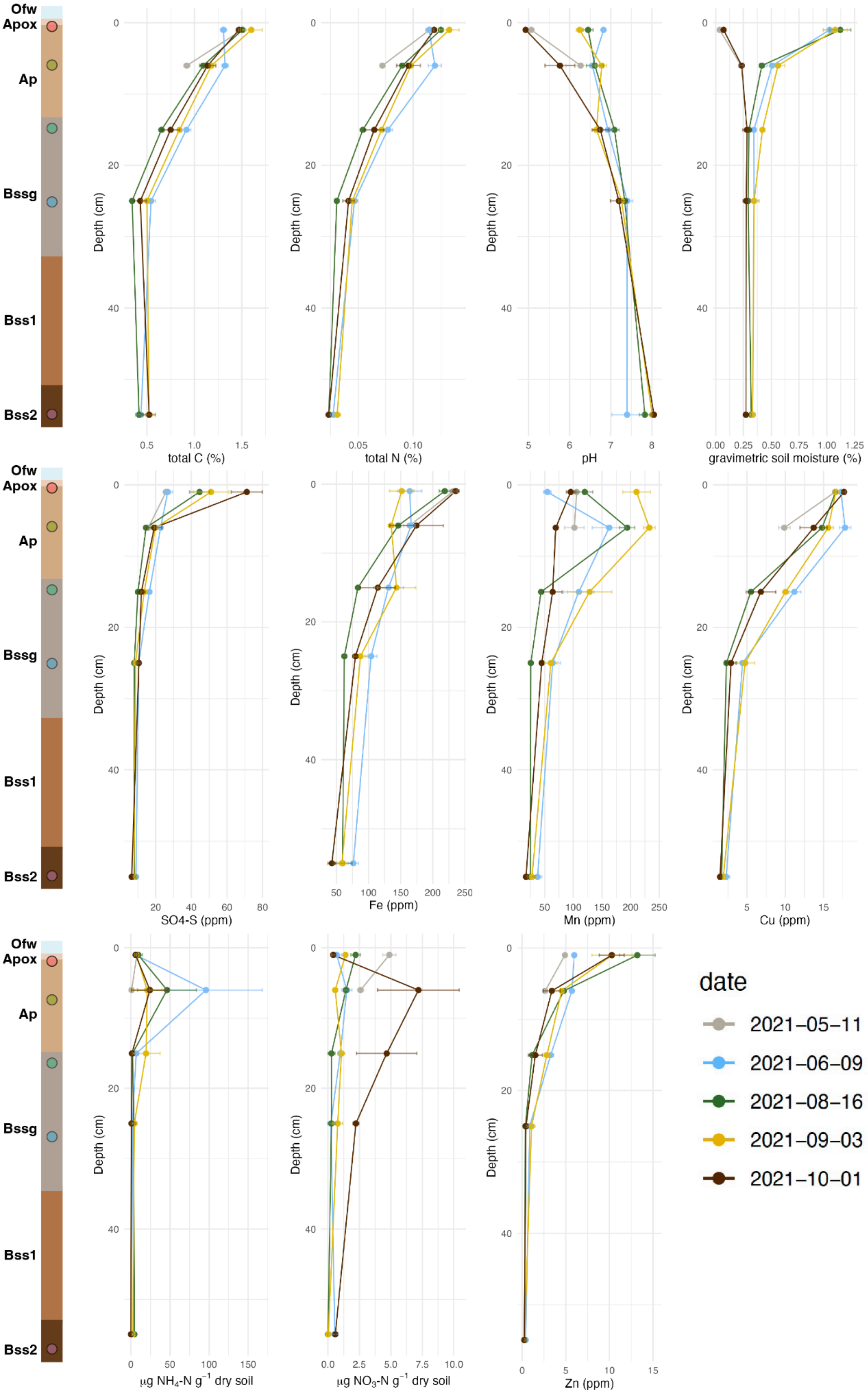
Ofw
Apox
Ap
Bssg
Bss1
Bss2
Depth (cm)
total C (%)
total N (%)
pH
gravimetric soil moisture (%)
SO4-S (ppm)
Fe (ppm)
Mn (ppm)
Cu (ppm)
µg NH4-N g-1 dry soil
µg NO3-N g-1 dry soil
Zn (ppm)
date
2021–05–11
2021–06–09
2021–08–16
2021–09–03
2021–10–01

**Fig. S3. Soil chemistry profiles across the growing season.**
Left: Cartoon of soil horizons. **Ofw** = surface organic layer influenced by floodwater; **Apox** = plowed A horizon, oxic conditions; **Ap** = plowed A horizon; **Bssg** = B horizon with slickensides and gleying; **Bss1** = upper B horizon with slickensides; **Bss2** = lower B horizon with slickensides. Right: Depth profiles for total carbon, total nitrogen, pH, gravimetric soil moisture, sulfate, iron, manganese, copper, ammonium, nitrate, and zinc, colored by date. Points represent averages across triplicate measurements and error bars give standard error.

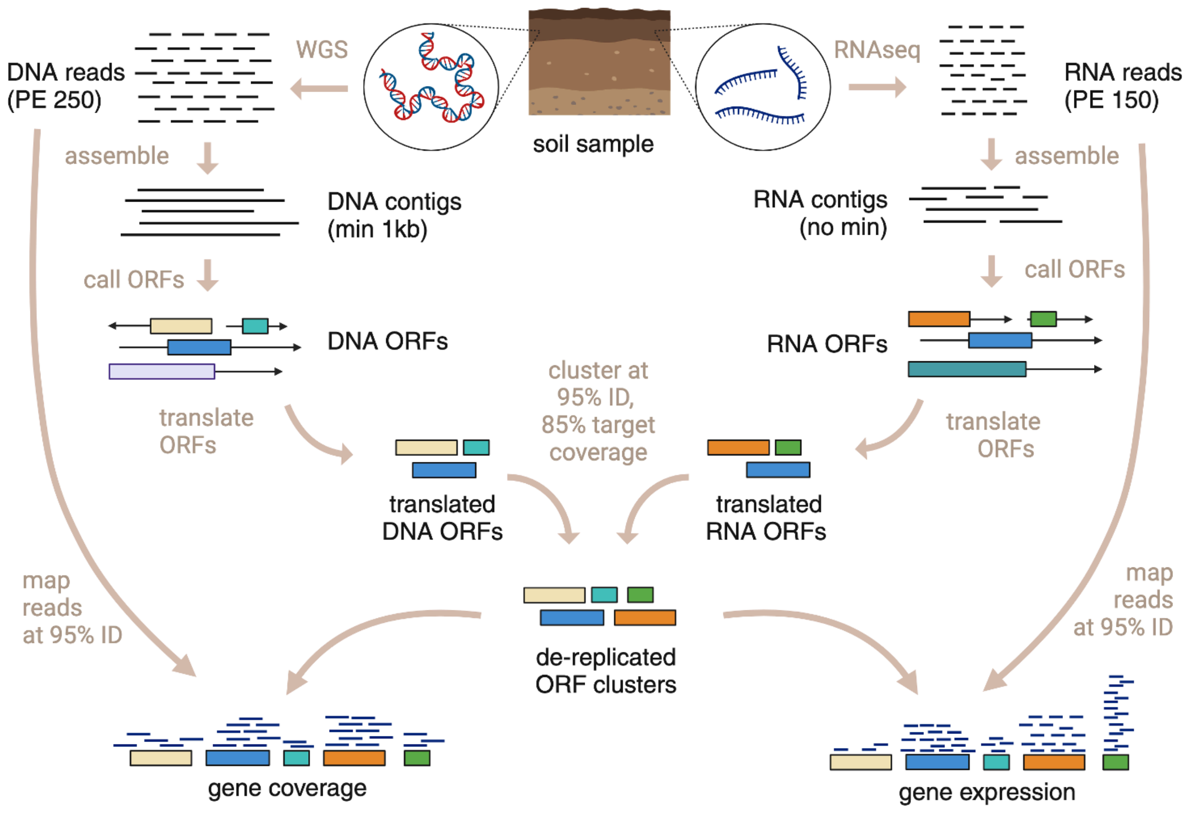


**Fig. S4. Genome-independent molecular analysis pipeline.** DNA and RNA were co-extracted from the same soil samples. DNA was sequenced using longer (PE250) reads than RNA (PE150), both on the Illumina platform. DNA and RNA contigs were assembled, and DNA contigs shorter than 1kb were discarded. Open reading frames (ORFs) were called on DNA and DNA contigs, and translated. The resulting proteins were clustered together with MMseqs2 easy-linclust at 95% identity and 85% target coverage, with one representative sequence chosen for each cluster. DNA and RNA reads were mapped back the contigs containing repseqs at 95% identity. For RNA, featureCounts was used to tally reads mapping specifically to repseqs.

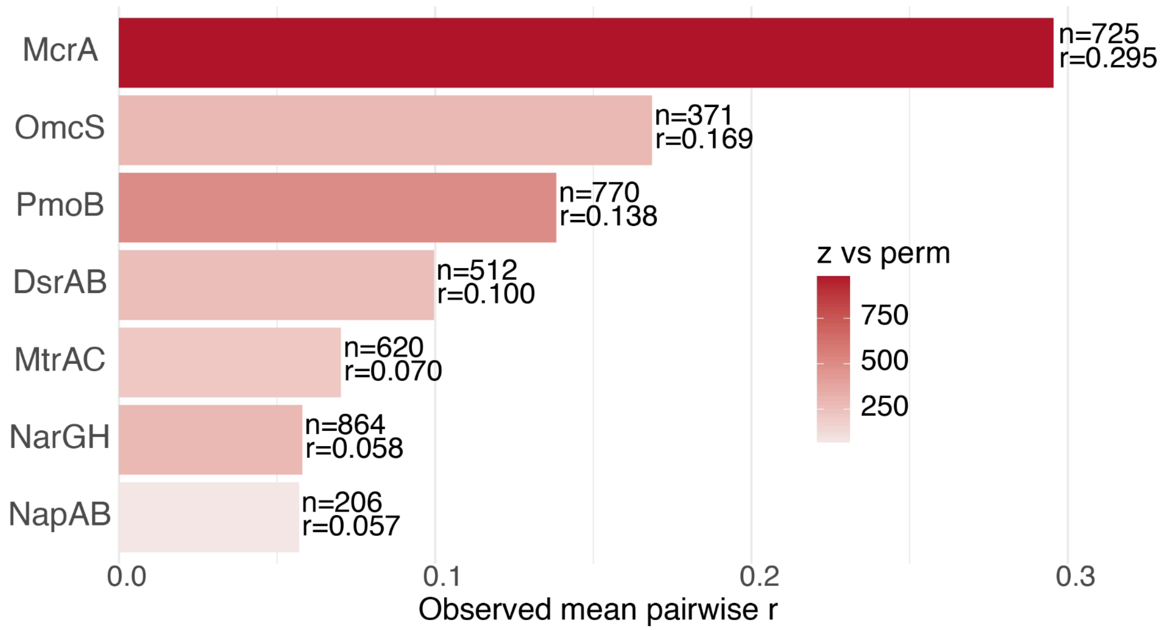


**Fig. S5. Coordination of expression of each TEAP marker across contributing species.** This analysis used 95% amino acid identity protein clusters from all genomic and transcriptomic scaffolds. For each marker family (DsrAB, NapAB, NarGH, MtrAC, McrA, OmcS, PmoB), clusters detected in at least 5 samples were retained. For each retained cluster, a 47-sample expression profile was built from date × depth × replicate transcript per million (TPM) values, log-transformed as log1p(TPM). The bar height shows the observed mean pairwise Pearson correlation among all cluster profiles within that marker family, and marker families are ordered from highest to lowest mean correlation. Bar color shows z vs perm, the number of null standard deviations by which the observed mean correlation exceeds a permutation-based null expectation generated by shuffling sample labels within clusters. Higher bars indicate stronger coordinated expression among repseqs assigned to the same marker family.

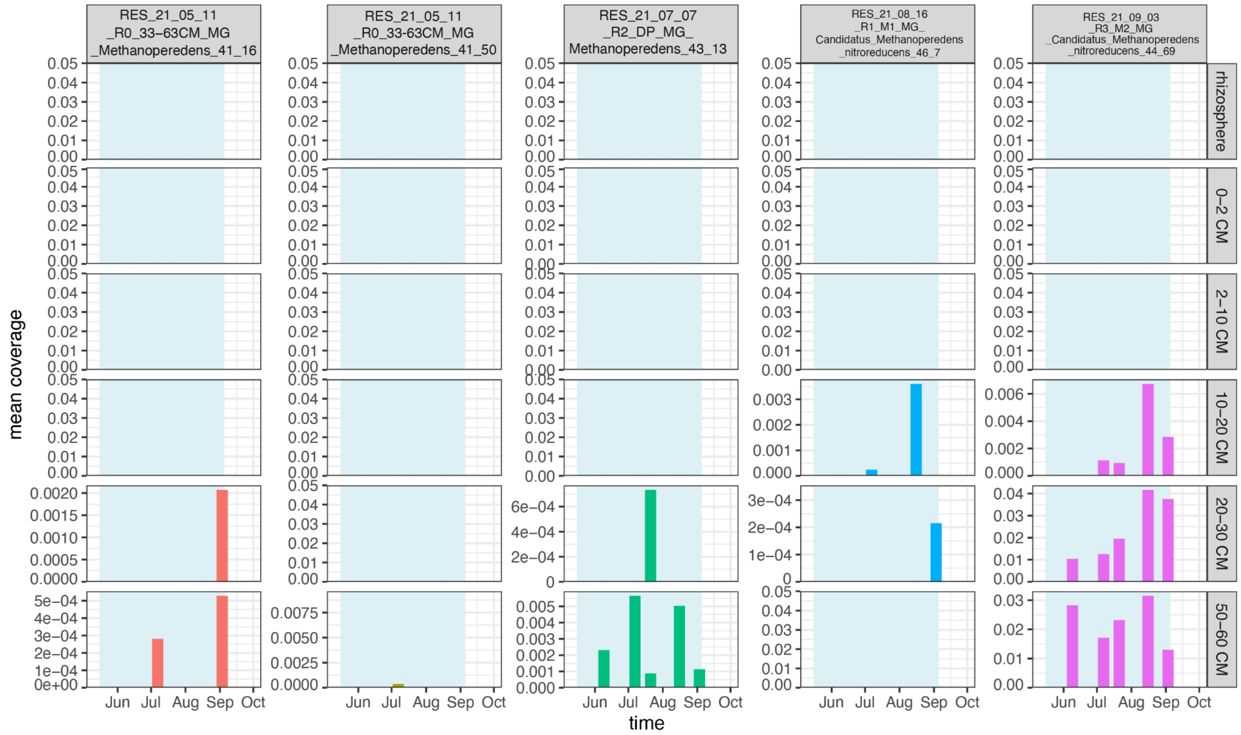


**Fig. S6. Abundance of *Methanoperedens* genomes across depth and time.** Each column represents a *Methanoperedens* genome. Each soil compartment (horizontal facets) is plotted on its own y-axis scale. Flooded timepoints are shaded in blue.

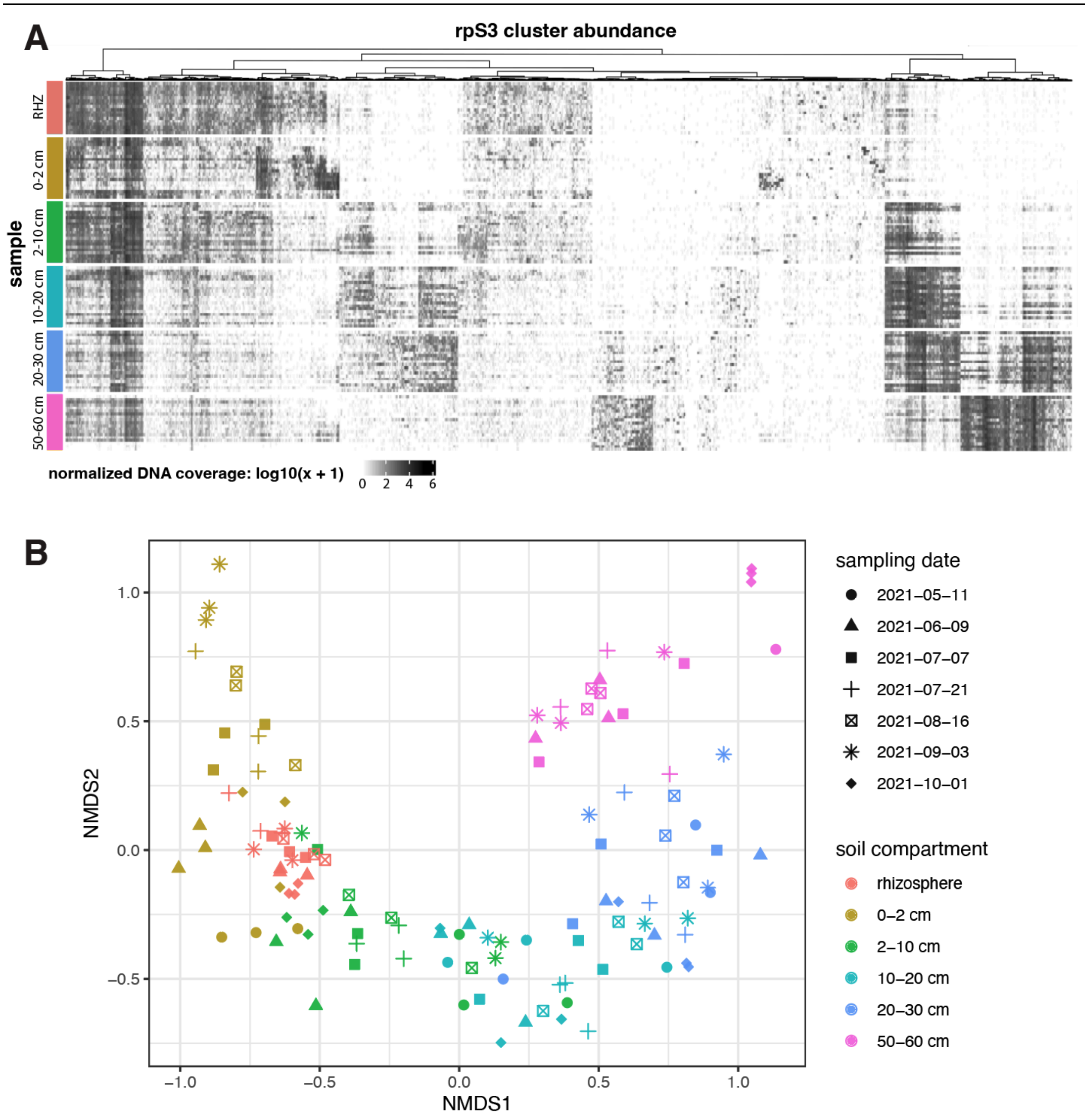


**Fig. S7. rpS3 community structure is stratified by depth. (A)** Heatmap of 99% ID rpS3 clusters abundance across samples, organized by soil depth (RHZ = rhizosphere, 0-2 cm, 2-10 cm, 10-20 cm, 20-30 cm, and 50-60 cm). Normalized coverage values were summed within each RPS3 cluster per sample and plotted as log10(x + 1), where x is length-normalized reads per million mapped reads (DNA coverage metric analogous to TPM). Rows represent samples grouped by soil depth, and columns represent RPS3 clusters hierarchically clustered by their abundance profiles. **(B)** NMDS ordination of rpS3 community composition based on Bray-Curtis dissimilarity. Each point represents one metagenomic sample from the rice paddy dataset.

Community composition was calculated from sample-wise, length-normalized relative abundance profiles of unbinned rpS3 clusters, and pairwise Bray-Curtis dissimilarities were ordinated with non-metric multidimensional scaling (NMDS). Points are colored by soil compartment and shaped by sampling date. In the associated PERMANOVA, compartment/depth explained the largest fraction of community variation ($R^2 = 0.4468$, $p = 1e\text{-}04$), whereas time was weaker ($R^2 = 0.0088$, $p = 0.0531$) and the depth-by-time interaction was not significant ($R^2 = 0.0302$, $p = 0.0913$).

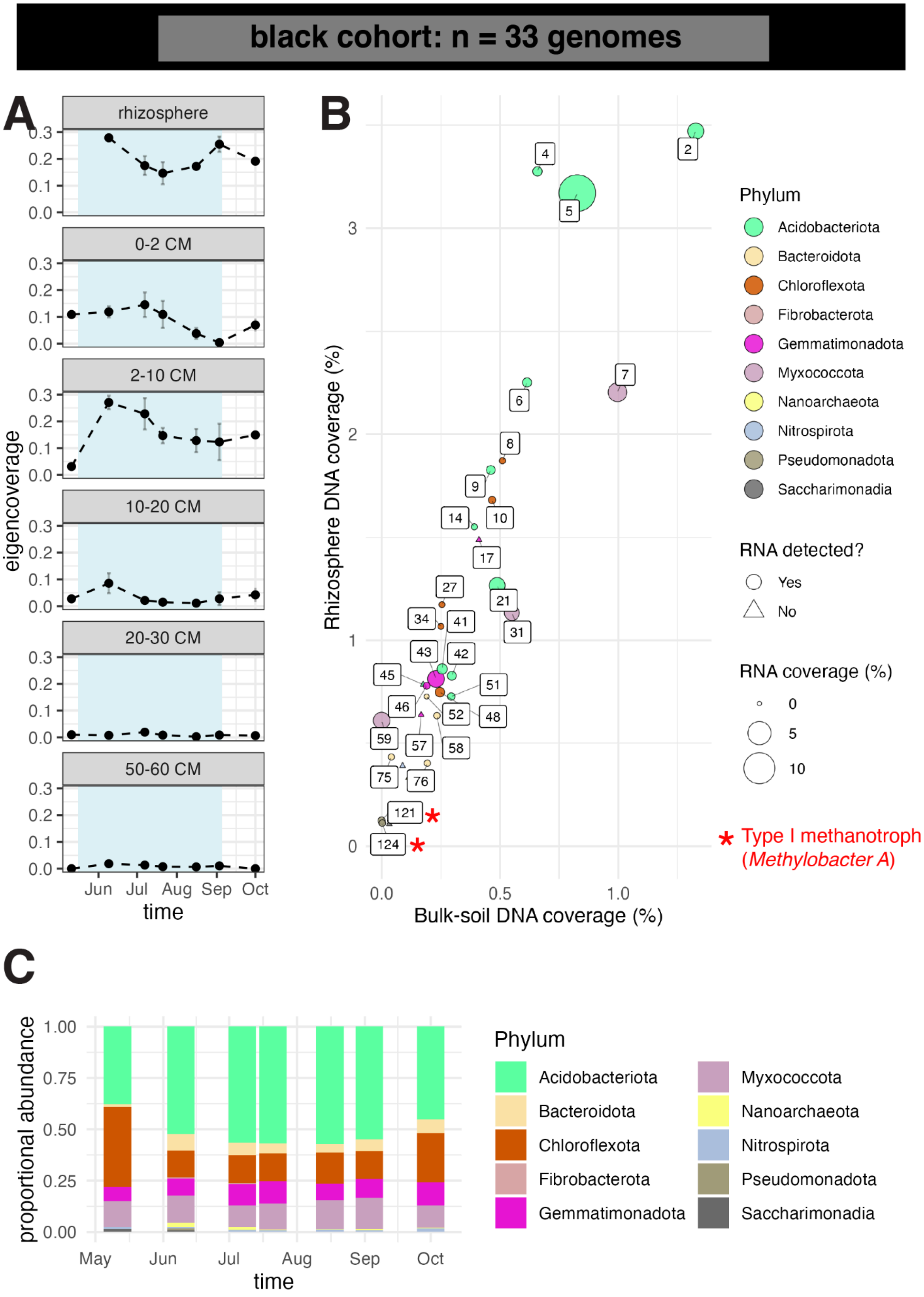
black cohort: n = 33 genomes
A
rhizosphere
0-2 CM
2-10 CM
10-20 CM
20-30 CM
50-60 CM
eigencoverage
Jun Jul Aug Sep Oct
time
B
Rhizosphere DNA coverage (%)
Bulk-soil DNA coverage (%)
Phylum
Acidobacteriota
Bacteroidota
Chloroflexota
Fibrobacterota
Gemmatimonadota
Myxococcota
Nanoarchaeota
Nitrospirota
Pseudomonadota
Saccharimonadia
RNA detected?
Yes
No
RNA coverage (%)
0
5
10
* Type I methanotroph (*Methylobacter A*)
C
proportional abundance
May Jun Jul Aug Sep Oct
time
Phylum
Acidobacteriota
Bacteroidota
Chloroflexota
Fibrobacterota
Gemmatimonadota
Myxococcota
Nanoarchaeota
Nitrospirota
Pseudomonadota
Saccharimonadia

**Fig. S8. Black cohort abundance, activity, and taxonomic composition. (A)** Mean abundance profiles of cohort genomes across time and soil compartments. Error bars show standard errors across biological replicates. Blue shading indicates flooded time points. **(B)** Genome abundance in the rhizosphere compared with bulk soil, colored by phylum. Genomes with detectable RNA are shown as circles and scaled by whole-genome RNA coverage estimated with CoverM. Genomes without detectable RNA are shown as triangles. Genomes are numbered by decreasing abundance across all samples; full genome names are provided in Data S1. **(C)** Phylum-level cohort composition over time across soil compartments.

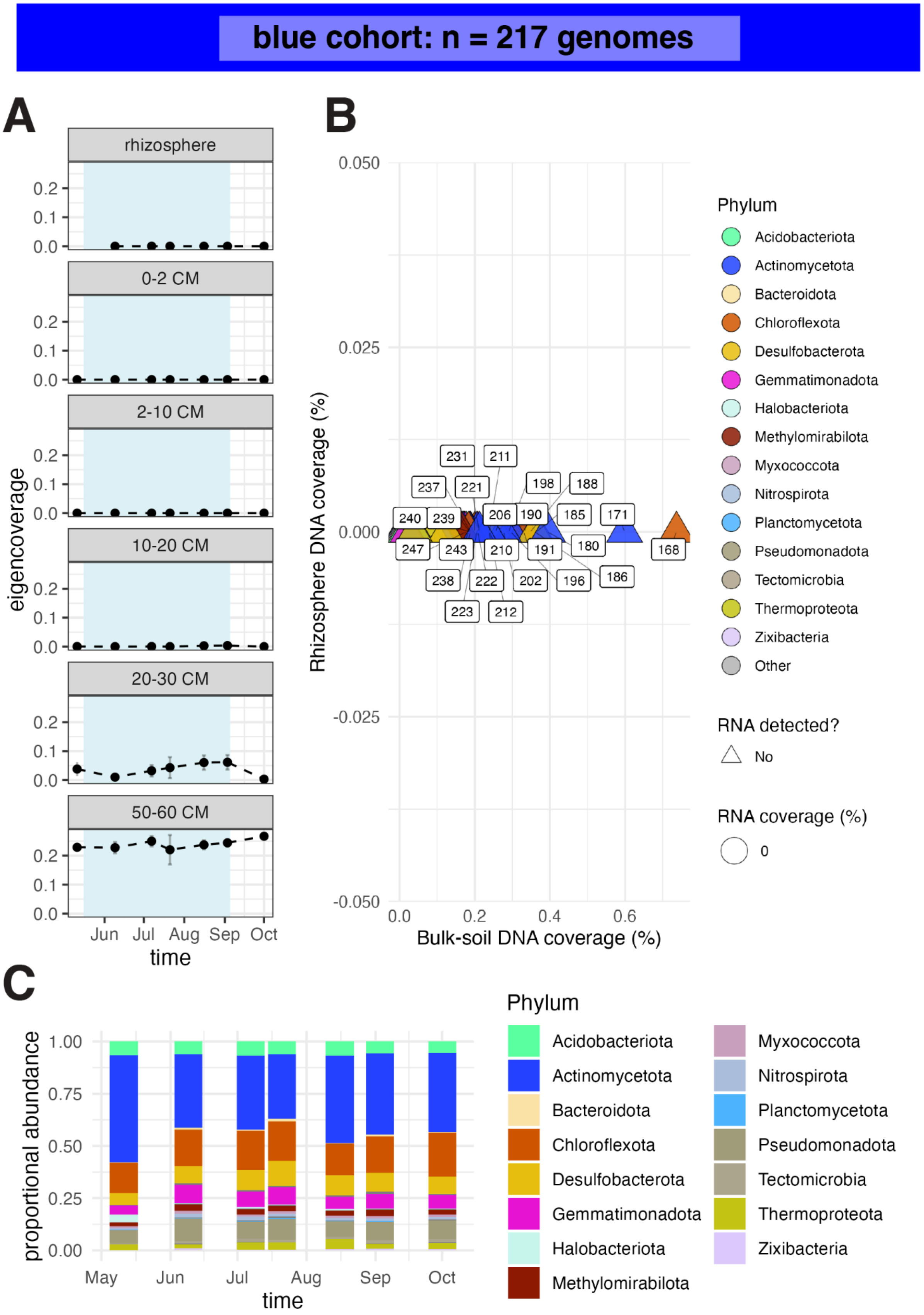

blue cohort: n = 217 genomes
A
rhizosphere
0-2 CM
2-10 CM
10-20 CM
20-30 CM
50-60 CM
eigencoverage
0.0
0.1
0.2
Jun
Jul
Aug
Sep
Oct
time
B
Rhizosphere DNA coverage (%)
0.050
0.025
0.000
-0.025
-0.050
0.0
0.2
0.4
0.6
Bulk-soil DNA coverage (%)
231
211
237
221
198
188
240
239
206
190
185
171
247
243
210
191
180
168
238
222
202
196
186
223
212
Phylum
Acidobacteriota
Actinomycetota
Bacteroidota
Chloroflexota
Desulfobacterota
Gemmatimonadota
Halobacteriota
Methylomirabilota
Myxococcota
Nitrospirota
Planctomycetota
Pseudomonadota
Tectomicrobia
Thermoproteota
Zixibacteria
Other
RNA detected?
No
RNA coverage (%)
0
C
proportional abundance
1.00
0.75
0.50
0.25
0.00
May
Jun
Jul
Aug
Sep
Oct
time
Phylum
Acidobacteriota
Actinomycetota
Bacteroidota
Chloroflexota
Desulfobacterota
Gemmatimonadota
Halobacteriota
Methylomirabilota
Myxococcota
Nitrospirota
Planctomycetota
Pseudomonadota
Tectomicrobia
Thermoproteota
Zixibacteria

**Fig. S9. Blue cohort abundance, activity, and taxonomic composition. (A)** Mean abundance profiles of cohort genomes across time and soil compartments. Error bars show standard errors across biological replicates. Blue shading indicates flooded time points. **(B)** Genome abundance in the rhizosphere compared with bulk soil, colored by phylum. Genomes with detectable RNA are shown as circles and scaled by whole-genome RNA coverage estimated with CoverM. Genomes without detectable RNA are shown as triangles. Genomes are numbered by decreasing abundance; full genome names are provided in Data S1. **(C)** Phylum-level cohort composition over time across soil compartments.

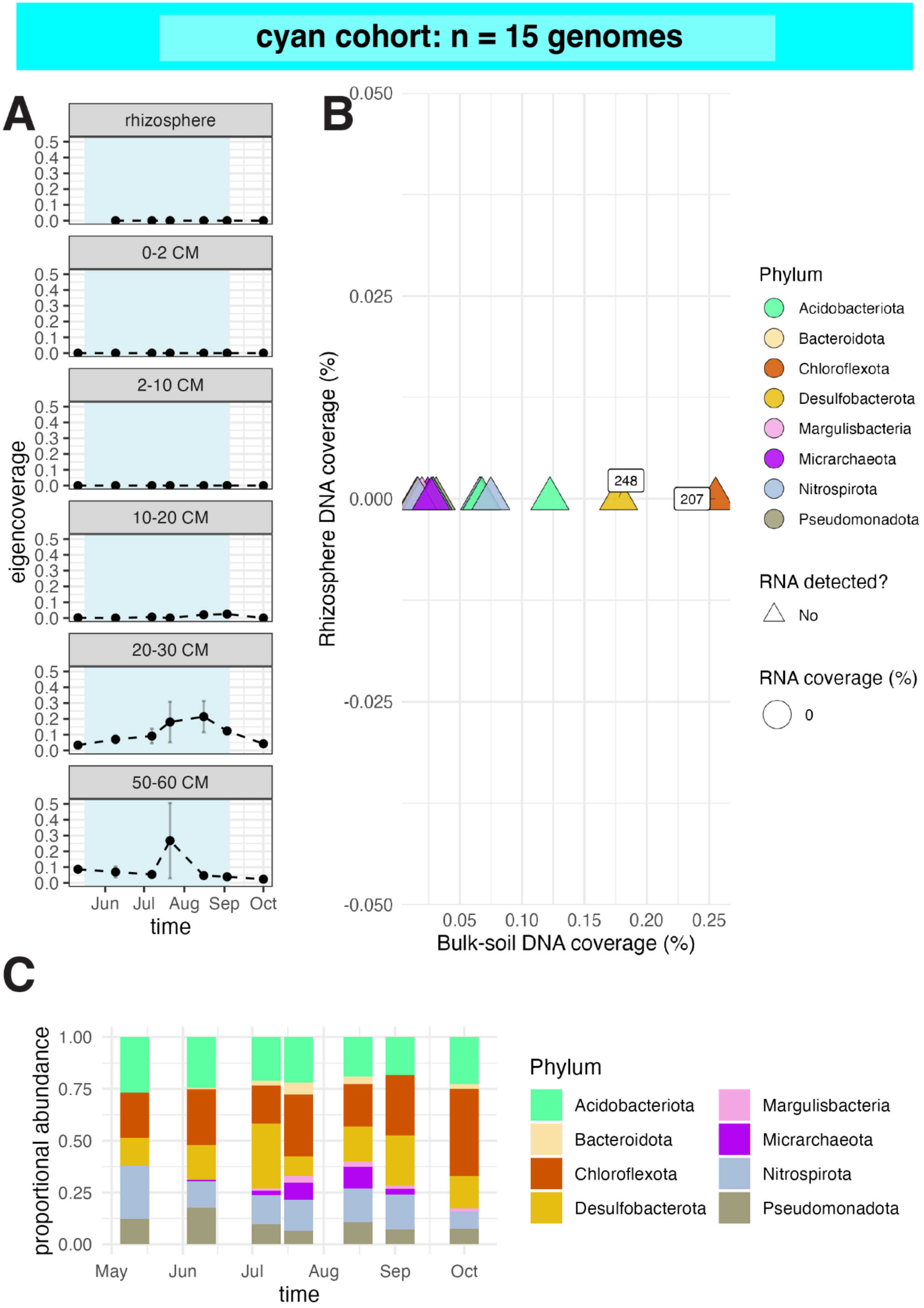

cyan cohort: n = 15 genomes
A
rhizosphere
0-2 CM
2-10 CM
10-20 CM
20-30 CM
50-60 CM
eigencoverage
0.0
0.1
0.2
0.3
0.4
0.5
Jun
Jul
Aug
Sep
Oct
time
B
Rhizosphere DNA coverage (%)
0.050
0.025
0.000
-0.025
-0.050
248
207
0.05
0.10
0.15
0.20
0.25
Bulk-soil DNA coverage (%)
Phylum
Acidobacteriota
Bacteroidota
Chloroflexota
Desulfobacterota
Margulisbacteria
Micrarchaeota
Nitrospirota
Pseudomonadota
RNA detected?
No
RNA coverage (%)
0
C
proportional abundance
0.00
0.25
0.50
0.75
1.00
May
Jun
Jul
Aug
Sep
Oct
time
Phylum
Acidobacteriota
Bacteroidota
Chloroflexota
Desulfobacterota
Margulisbacteria
Micrarchaeota
Nitrospirota
Pseudomonadota

**Fig. S10. Cyan cohort abundance, activity, and taxonomic composition. (A)** Mean abundance profiles of cohort genomes across time and soil compartments. Error bars show standard errors across biological replicates. Blue shading indicates flooded time points. **(B)** Genome abundance in the rhizosphere compared with bulk soil, colored by phylum. Genomes with detectable RNA are shown as circles and scaled by whole-genome RNA coverage estimated with CoverM. Genomes without detectable RNA are shown as triangles. Genomes are numbered by decreasing abundance; full genome names are provided in Data S1. **(C)** Phylum-level cohort composition over time across soil compartments.

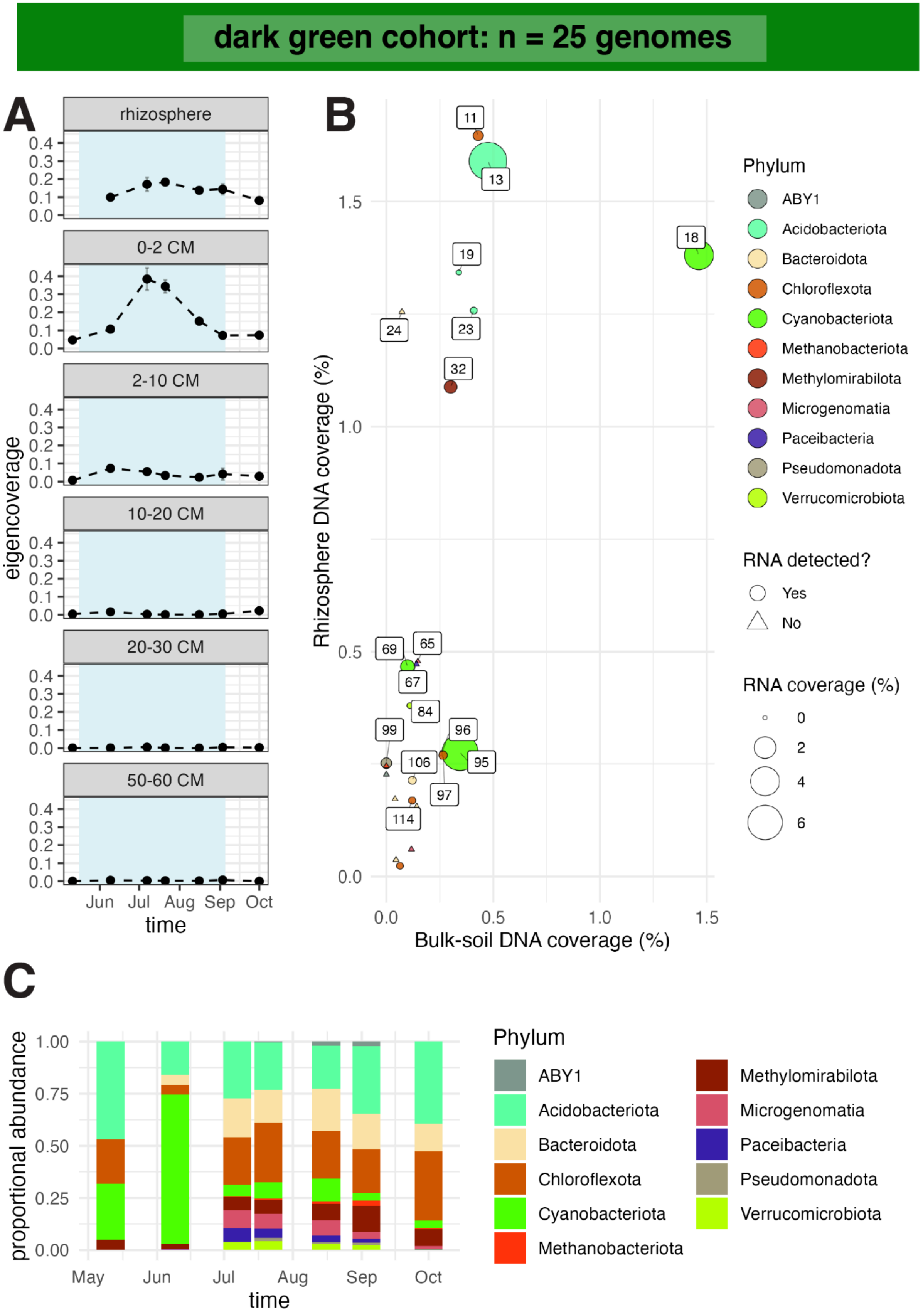
dark green cohort: n = 25 genomes
A
rhizosphere
0-2 CM
2-10 CM
10-20 CM
20-30 CM
50-60 CM
eigencoverage
0.0
0.1
0.2
0.3
0.4
Jun
Jul
Aug
Sep
Oct
time
B
11
13
18
19
24
23
32
69
65
67
84
99
96
106
95
97
114
Rhizosphere DNA coverage (%)
0.0
0.5
1.0
1.5
Bulk-soil DNA coverage (%)
Phylum
ABY1
Acidobacteriota
Bacteroidota
Chloroflexota
Cyanobacteriota
Methanobacteriota
Methylomirabilota
Microgenomatia
Paceibacteria
Pseudomonadota
Verrucomicrobiota
RNA detected?
Yes
No
RNA coverage (%)
0
2
4
6
C
proportional abundance
0.00
0.25
0.50
0.75
1.00
May
Jun
Jul
Aug
Sep
Oct
time
Phylum
ABY1
Acidobacteriota
Bacteroidota
Chloroflexota
Cyanobacteriota
Methanobacteriota
Methylomirabilota
Microgenomatia
Paceibacteria
Pseudomonadota
Verrucomicrobiota

**Fig. S11. Darkgreen cohort abundance, activity, and taxonomic composition. (A)** Mean abundance profiles of cohort genomes across time and soil compartments. Error bars show standard errors across biological replicates. Blue shading indicates flooded time points. **(B)** Genome abundance in the rhizosphere compared with bulk soil, colored by phylum. Genomes with detectable RNA are shown as circles and scaled by whole-genome RNA coverage estimated with CoverM. Genomes without detectable RNA are shown as triangles. Genomes are numbered by decreasing abundance; full genome names are provided in Data S1. **(C)** Phylum-level cohort

composition over time across soil compartments.

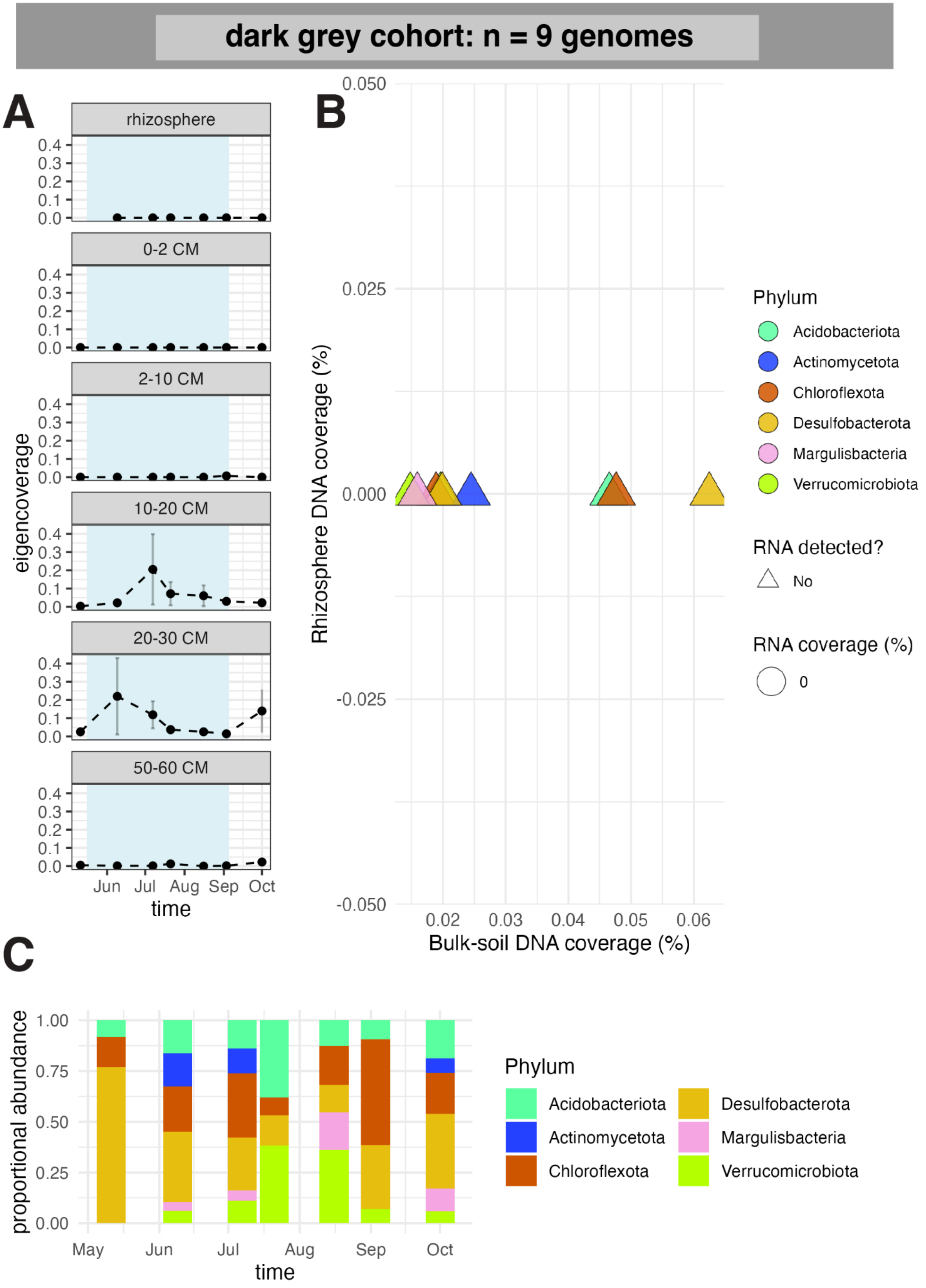
dark grey cohort: n = 9 genomes
A
rhizosphere
0-2 CM
2-10 CM
10-20 CM
20-30 CM
50-60 CM
eigencoverage
0.4
0.3
0.2
0.1
0.0
Jun
Jul
Aug
Sep
Oct
time
B
0.050
0.025
0.000
-0.025
-0.050
Rhizosphere DNA coverage (%)
0.02
0.03
0.04
0.05
0.06
Bulk-soil DNA coverage (%)
Phylum
Acidobacteriota
Actinomycetota
Chloroflexota
Desulfobacterota
Margulisbacteria
Verrucomicrobiota
RNA detected?
No
RNA coverage (%)
0
C
proportional abundance
1.00
0.75
0.50
0.25
0.00
May
Jun
Jul
Aug
Sep
Oct
time
Phylum
Acidobacteriota
Actinomycetota
Chloroflexota
Desulfobacterota
Margulisbacteria
Verrucomicrobiota

**Fig. S12. Darkgrey cohort abundance, activity, and taxonomic composition. (A)** Mean abundance profiles of cohort genomes across time and soil compartments. Error bars show standard errors across biological replicates. Blue shading indicates flooded time points. **(B)** Genome abundance in the rhizosphere compared with bulk soil, colored by phylum. Genomes with detectable RNA are shown as circles and scaled by whole-genome RNA coverage estimated with CoverM. Genomes without detectable RNA are shown as triangles. Genomes are numbered by decreasing abundance; full genome names are provided in Data S1. **(C)** Phylum-level cohort composition over time across soil compartments.

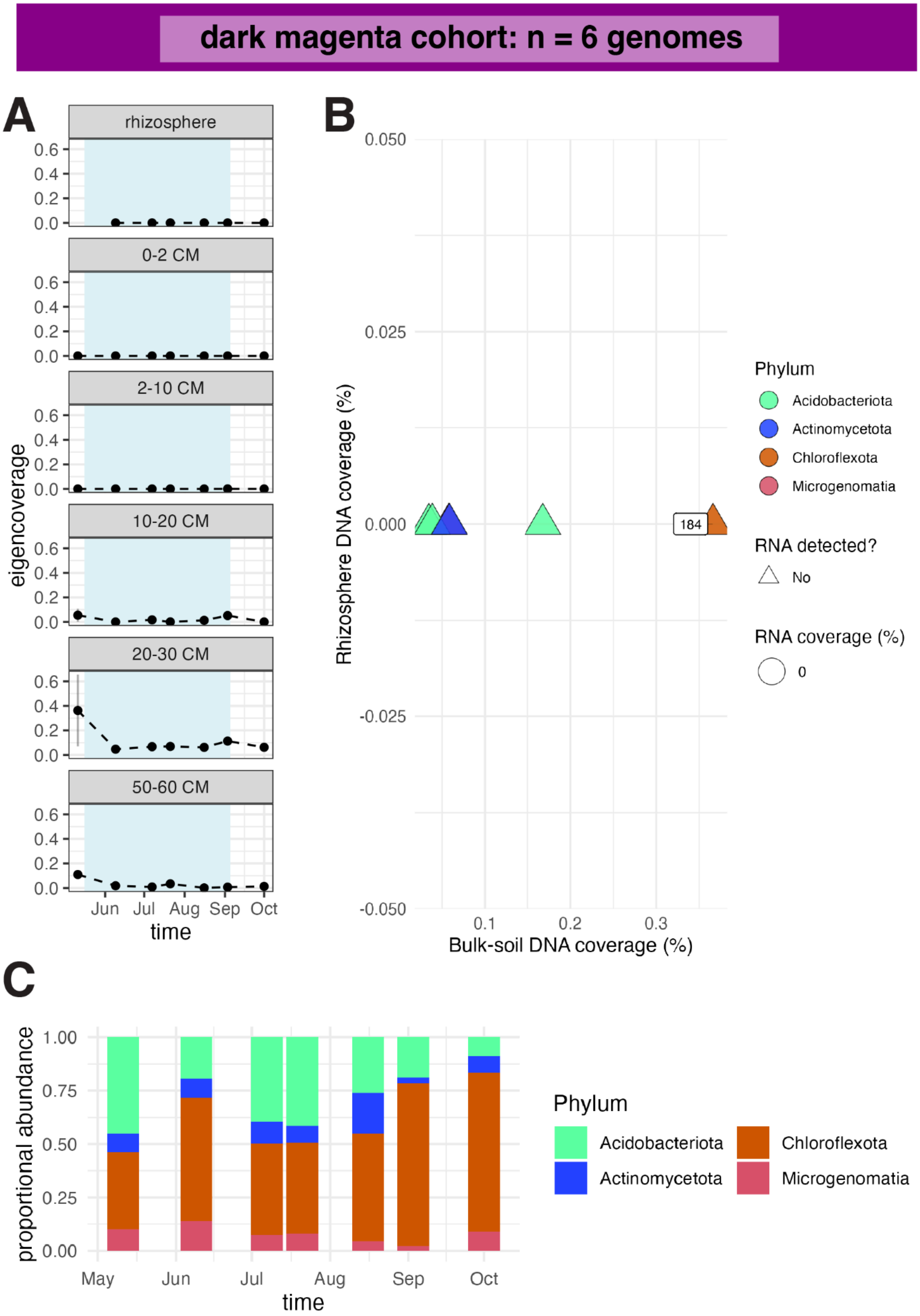
dark magenta cohort: n = 6 genomes
A
rhizosphere
0-2 CM
2-10 CM
10-20 CM
20-30 CM
50-60 CM
eigencoverage
0.6
0.4
0.2
0.0
Jun Jul Aug Sep Oct
time
B
Rhizosphere DNA coverage (%)
0.050
0.025
0.000
-0.025
-0.050
184
0.1 0.2 0.3
Bulk-soil DNA coverage (%)
Phylum
Acidobacteriota
Actinomycetota
Chloroflexota
Microgenomatia
RNA detected?
No
RNA coverage (%)
0
C
proportional abundance
1.00
0.75
0.50
0.25
0.00
May Jun Jul Aug Sep Oct
time
Phylum
Acidobacteriota
Actinomycetota
Chloroflexota
Microgenomatia

**Fig. S13. Darkmagenta cohort abundance, activity, and taxonomic composition. (A)** Mean abundance profiles of cohort genomes across time and soil compartments. Error bars show standard errors across biological replicates. Blue shading indicates flooded time points. **(B)** Genome abundance in the rhizosphere compared with bulk soil, colored by phylum. Genomes with detectable RNA are shown as circles and scaled by whole-genome RNA coverage estimated with CoverM. Genomes without detectable RNA are shown as triangles. Genomes are numbered by decreasing abundance; full genome names are provided in Data S1. **(C)** Phylum-level cohort composition over time across soil compartments.

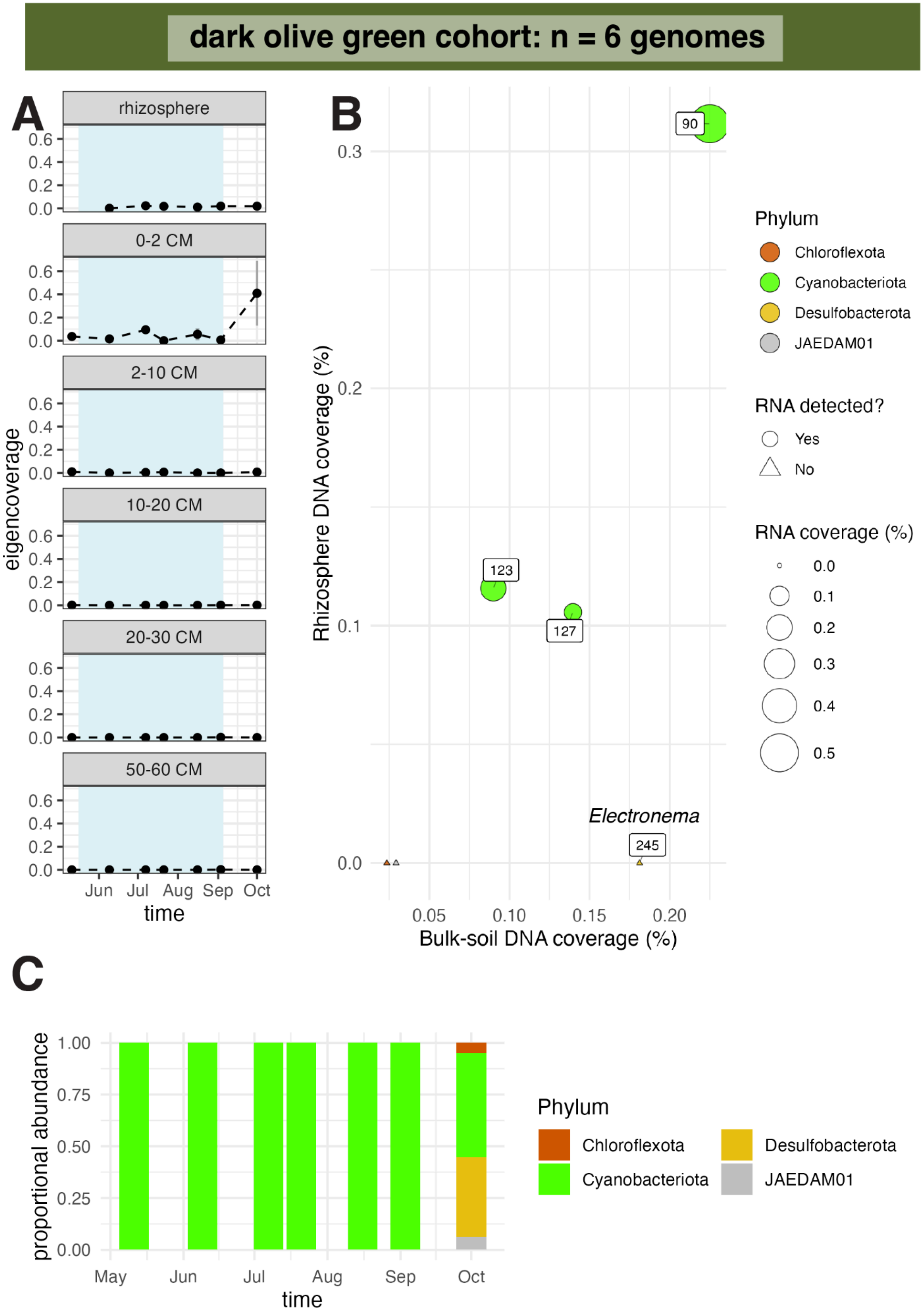

dark olive green cohort: n = 6 genomes
A
rhizosphere
0-2 CM
2-10 CM
10-20 CM
20-30 CM
50-60 CM
eigencoverage
Jun Jul Aug Sep Oct
time
B
Rhizosphere DNA coverage (%)
Bulk-soil DNA coverage (%)
90
123
127
245
Electronema
Phylum
Chloroflexota
Cyanobacteriota
Desulfobacterota
JAEDAM01
RNA detected?
Yes
No
RNA coverage (%)
C
proportional abundance
May Jun Jul Aug Sep Oct
time
Phylum
Chloroflexota
Desulfobacterota
Cyanobacteriota
JAEDAM01

**Fig. S14. Darkolivegreen cohort abundance, activity, and taxonomic composition. (A)** Mean abundance profiles of cohort genomes across time and soil compartments. Error bars show standard errors across biological replicates. Blue shading indicates flooded time points. **(B)** Genome abundance in the rhizosphere compared with bulk soil, colored by phylum. Genomes with detectable RNA are shown as circles and scaled by whole-genome RNA coverage estimated with CoverM. Genomes without detectable RNA are shown as triangles. Genomes are numbered by decreasing abundance; full genome names are provided in Data S1. **(C)** Phylum-level cohort

composition over time across soil compartments.

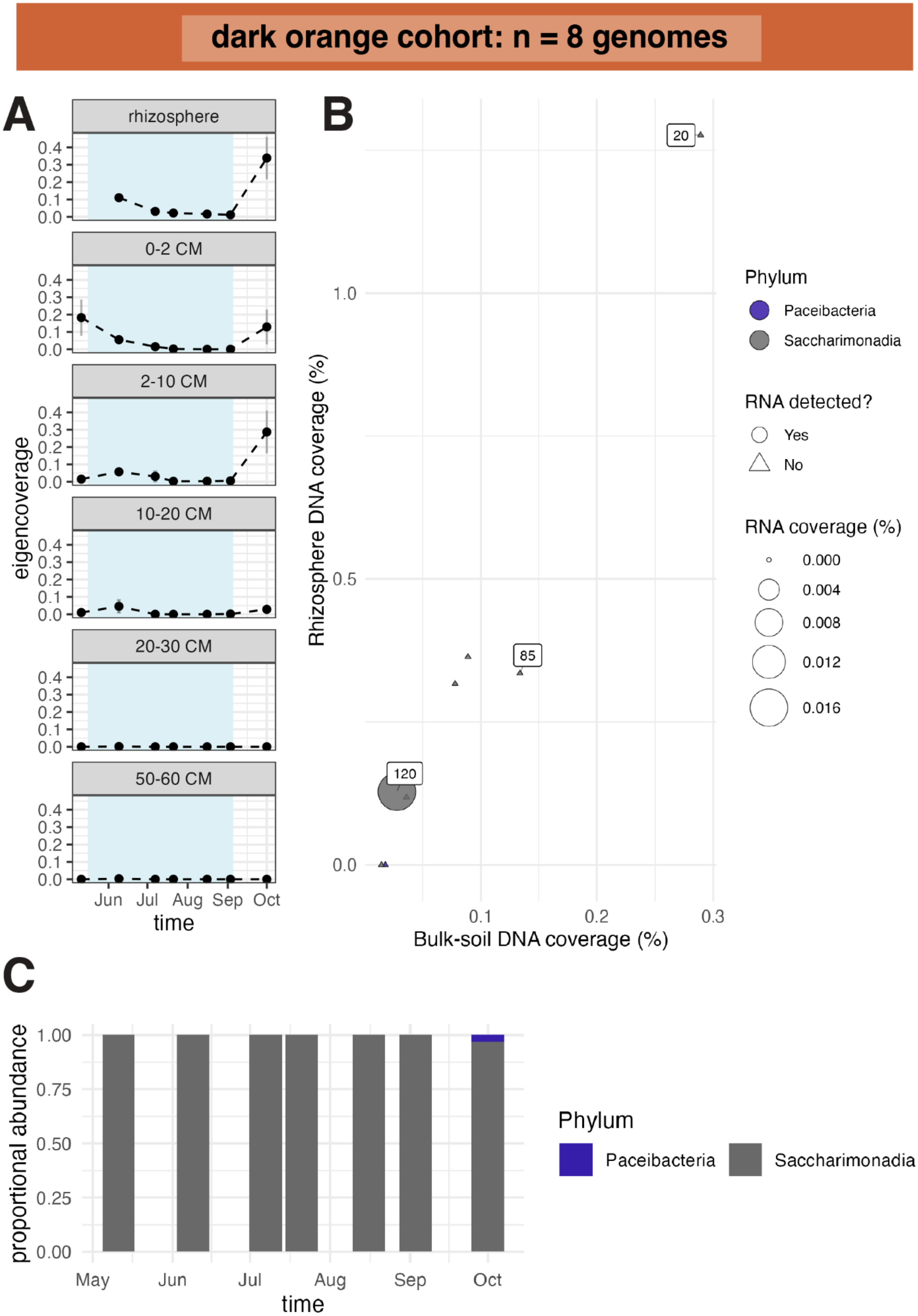
dark orange cohort: n = 8 genomes
A
rhizosphere
0-2 CM
2-10 CM
10-20 CM
20-30 CM
50-60 CM
eigencoverage
0.0
0.1
0.2
0.3
0.4
Jun
Jul
Aug
Sep
Oct
time
B
20
85
120
Rhizosphere DNA coverage (%)
0.0
0.5
1.0
0.1
0.2
0.3
Bulk-soil DNA coverage (%)
Phylum
Paceibacteria
Saccharimonadia
RNA detected?
Yes
No
RNA coverage (%)
0.000
0.004
0.008
0.012
0.016
C
proportional abundance
0.00
0.25
0.50
0.75
1.00
May
Jun
Jul
Aug
Sep
Oct
time
Phylum
Paceibacteria
Saccharimonadia

**Fig. S15. Darkorange cohort abundance, activity, and taxonomic composition. (A)** Mean abundance profiles of cohort genomes across time and soil compartments. Error bars show standard errors across biological replicates. Blue shading indicates flooded time points. **(B)** Genome abundance in the rhizosphere compared with bulk soil, colored by phylum. Genomes with detectable RNA are shown as circles and scaled by whole-genome RNA coverage estimated with CoverM. Genomes without detectable RNA are shown as triangles. Genomes are numbered by decreasing abundance; full genome names are provided in Data S1. **(C)** Phylum-level cohort composition over time across soil compartments.

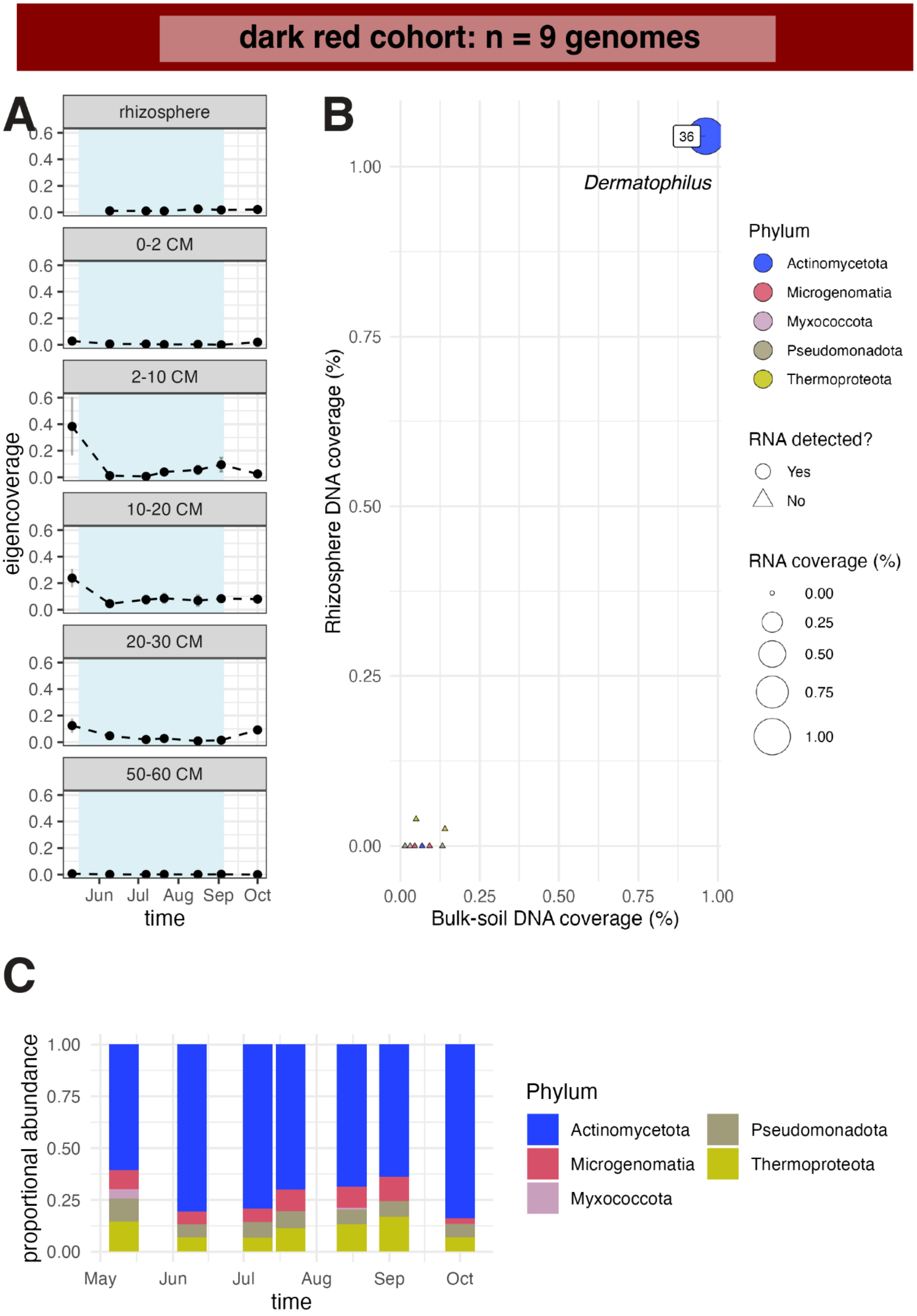

dark red cohort: n = 9 genomes
A
rhizosphere
0-2 CM
2-10 CM
10-20 CM
20-30 CM
50-60 CM
eigencoverage
0.0
0.2
0.4
0.6
Jun
Jul
Aug
Sep
Oct
time
B
36
Dermatophilus
Rhizosphere DNA coverage (%)
Bulk-soil DNA coverage (%)
0.00
0.25
0.50
0.75
1.00
Phylum
Actinomycetota
Microgenomatia
Myxococcota
Pseudomonadota
Thermoproteota
RNA detected?
Yes
No
RNA coverage (%)
0.00
0.25
0.50
0.75
1.00
C
proportional abundance
0.00
0.25
0.50
0.75
1.00
May
Jun
Jul
Aug
Sep
Oct
time
Phylum
Actinomycetota
Microgenomatia
Myxococcota
Pseudomonadota
Thermoproteota

**Fig. S16. Dark red cohort abundance, activity, and taxonomic composition. (A)** Mean abundance profiles of cohort genomes across time and soil compartments. Error bars show standard errors across biological replicates. Blue shading indicates flooded time points. **(B)** Genome abundance in the rhizosphere compared with bulk soil, colored by phylum. Genomes with detectable RNA are shown as circles and scaled by whole-genome RNA coverage estimated with CoverM. Genomes without detectable RNA are shown as triangles. Genomes are numbered by decreasing abundance; full genome names are provided in Data S1. **(C)** Phylum-level cohort composition over time across soil compartments.

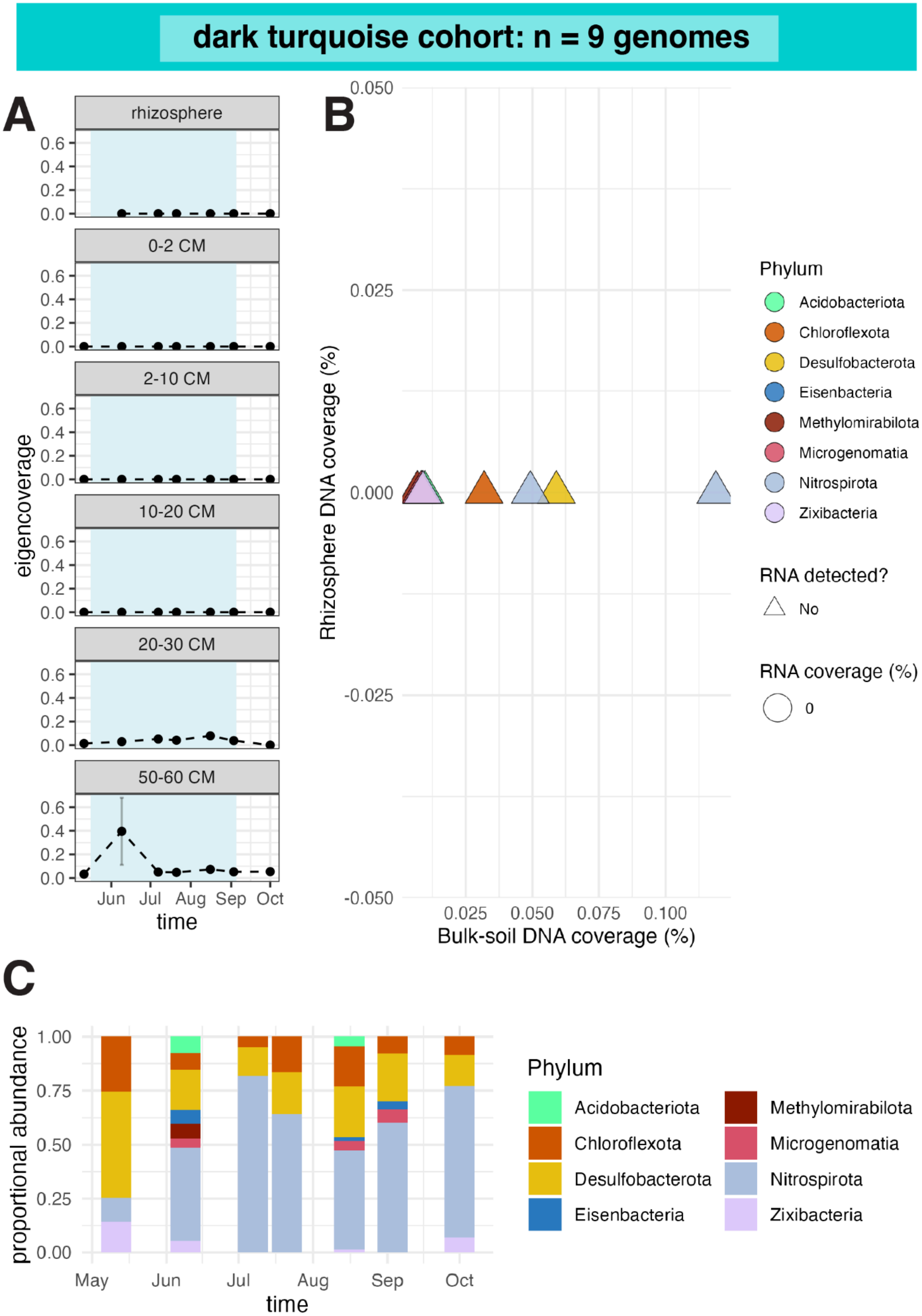
dark turquoise cohort: n = 9 genomes
A
rhizosphere
0-2 CM
2-10 CM
10-20 CM
20-30 CM
50-60 CM
0.6
0.4
0.2
0.0
eigencoverage
Jun Jul Aug Sep Oct
time
B
0.050
0.025
0.000
-0.025
-0.050
Rhizosphere DNA coverage (%)
0.025 0.050 0.075 0.100
Bulk-soil DNA coverage (%)
Phylum
Acidobacteriota
Chloroflexota
Desulfobacterota
Eisenbacteria
Methylomirabilota
Microgenomatia
Nitrospirota
Zixibacteria
RNA detected?
No
RNA coverage (%)
0
C
1.00
0.75
0.50
0.25
0.00
proportional abundance
May Jun Jul Aug Sep Oct
time
Phylum
Acidobacteriota
Chloroflexota
Desulfobacterota
Eisenbacteria
Methylomirabilota
Microgenomatia
Nitrospirota
Zixibacteria

**Fig. S17. Dark turquoise cohort abundance, activity, and taxonomic composition. (A)** Mean abundance profiles of cohort genomes across time and soil compartments. Error bars show standard errors across biological replicates. Blue shading indicates flooded time points. **(B)** Genome abundance in the rhizosphere compared with bulk soil, colored by phylum. Genomes with detectable RNA are shown as circles and scaled by whole-genome RNA coverage estimated with CoverM. Genomes without detectable RNA are shown as triangles. Genomes are numbered by decreasing abundance; full genome names are provided in Data S1. **(C)** Phylum-level cohort

composition over time across soil compartments.

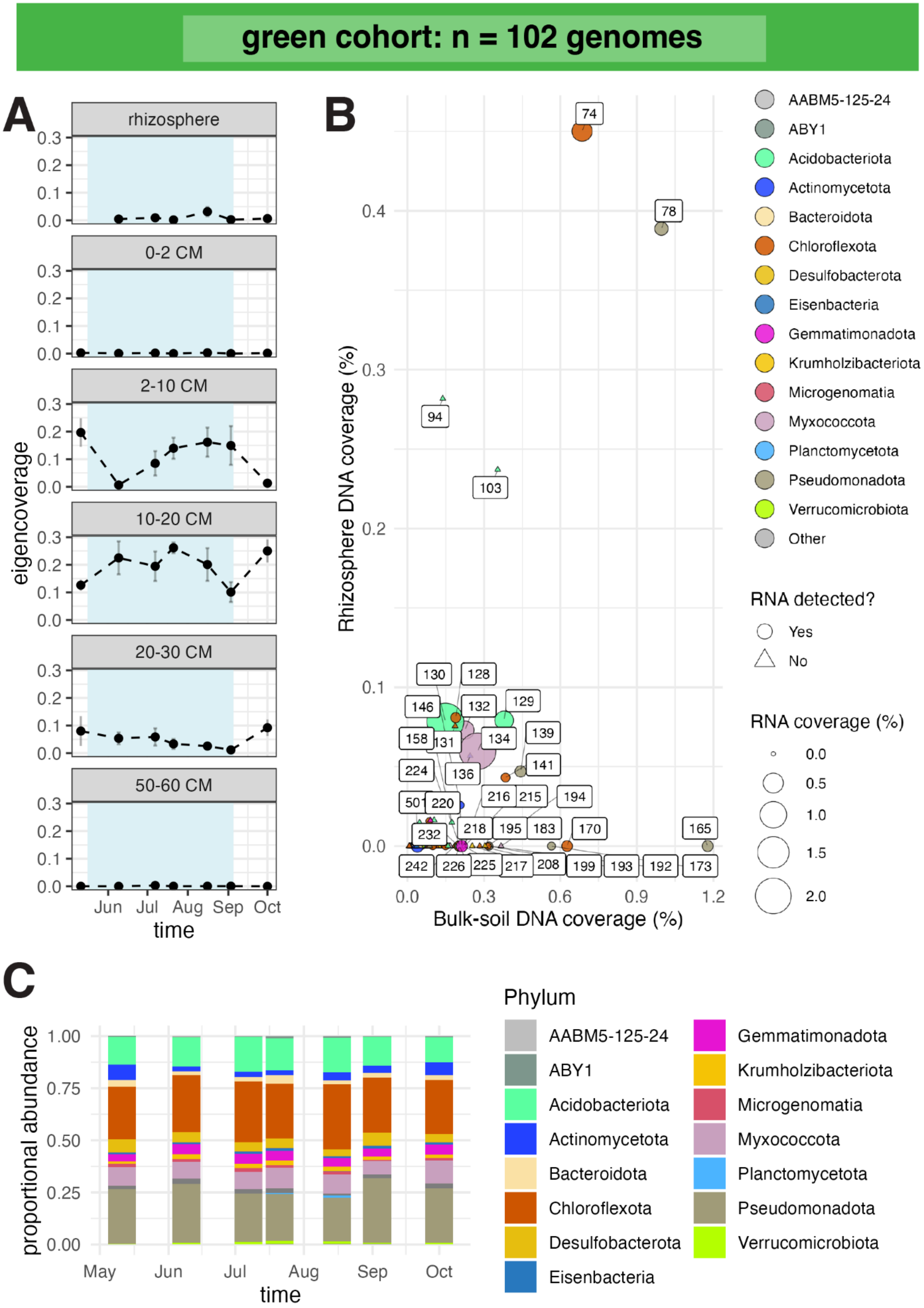

green cohort: n = 102 genomes
A
rhizosphere
0-2 CM
2-10 CM
10-20 CM
20-30 CM
50-60 CM
eigencoverage
Jun Jul Aug Sep Oct
time
B
Rhizosphere DNA coverage (%)
Bulk-soil DNA coverage (%)
AABM5-125-24
ABY1
Acidobacteriota
Actinomycetota
Bacteroidota
Chloroflexota
Desulfobacterota
Eisenbacteria
Gemmatimonadota
Krumholzibacteriota
Microgenomatia
Myxococcota
Planctomycetota
Pseudomonadota
Verrucomicrobiota
Other
RNA detected?
Yes
No
RNA coverage (%)
C
proportional abundance
May Jun Jul Aug Sep Oct
time
Phylum
AABM5-125-24
ABY1
Acidobacteriota
Actinomycetota
Bacteroidota
Chloroflexota
Desulfobacterota
Eisenbacteria
Gemmatimonadota
Krumholzibacteriota
Microgenomatia
Myxococcota
Planctomycetota
Pseudomonadota
Verrucomicrobiota

**Fig. S18. Green cohort abundance, activity, and taxonomic composition. (A)** Mean abundance profiles of cohort genomes across time and soil compartments. Error bars show standard errors across biological replicates. Blue shading indicates flooded time points. **(B)** Genome abundance in the rhizosphere compared with bulk soil, colored by phylum. Genomes with detectable RNA are shown as circles and scaled by whole-genome RNA coverage estimated with CoverM. Genomes without detectable RNA are shown as triangles. Genomes are numbered by decreasing abundance; full genome names are provided in Data S1. **(C)** Phylum-level cohort

composition over time across soil compartments.

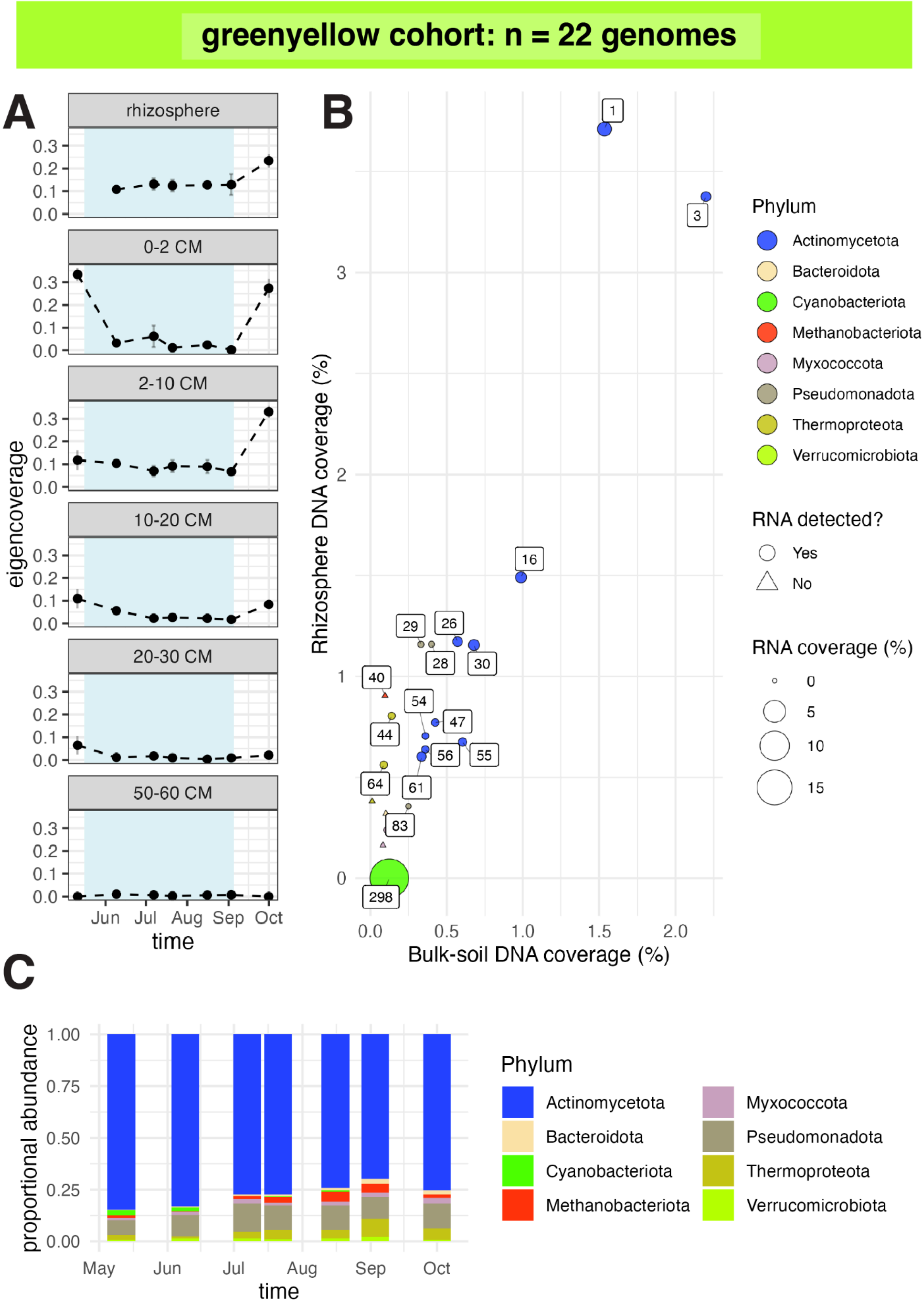
greenyellow cohort: n = 22 genomes
A
rhizosphere
0-2 CM
2-10 CM
10-20 CM
20-30 CM
50-60 CM
eigencoverage
Jun Jul Aug Sep Oct
time
B
Rhizosphere DNA coverage (%)
Bulk-soil DNA coverage (%)
1
3
16
29
26
28
30
40
54
47
44
56
55
64
61
83
298
Phylum
Actinomycetota
Bacteroidota
Cyanobacteriota
Methanobacteriota
Myxococcota
Pseudomonadota
Thermoproteota
Verrucomicrobiota
RNA detected?
Yes
No
RNA coverage (%)
0
5
10
15
C
proportional abundance
May Jun Jul Aug Sep Oct
time
Phylum
Actinomycetota
Bacteroidota
Cyanobacteriota
Methanobacteriota
Myxococcota
Pseudomonadota
Thermoproteota
Verrucomicrobiota

**Fig. S19. Greenyellow cohort abundance, activity, and taxonomic composition. (A)** Mean abundance profiles of cohort genomes across time and soil compartments. Error bars show standard errors across biological replicates. Blue shading indicates flooded time points. **(B)** Genome abundance in the rhizosphere compared with bulk soil, colored by phylum. Genomes with detectable RNA are shown as circles and scaled by whole-genome RNA coverage estimated with CoverM. Genomes without detectable RNA are shown as triangles. Genomes are numbered by decreasing abundance; full genome names are provided in Data S1. **(C)** Phylum-level cohort composition over time across soil compartments.

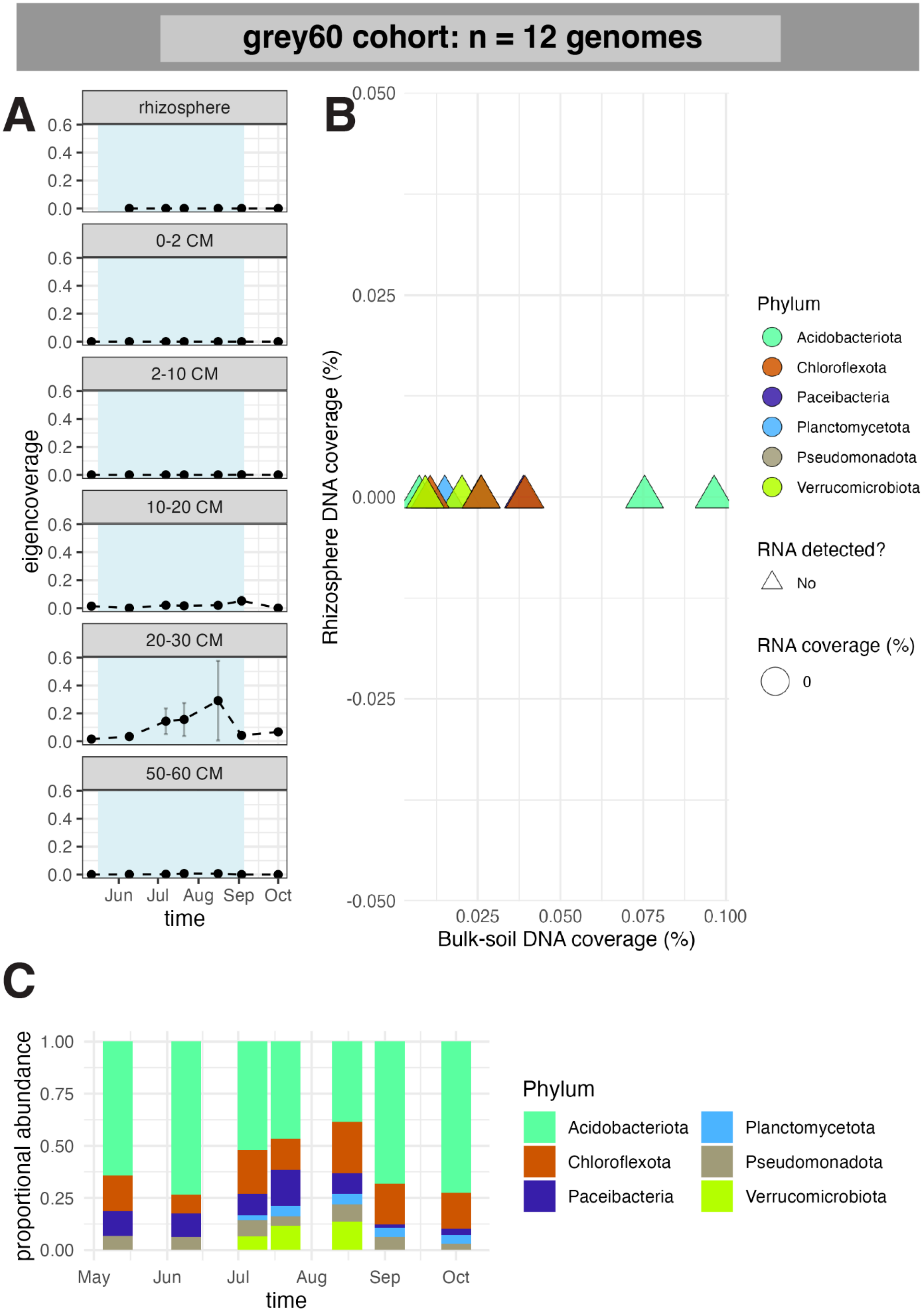

grey60 cohort: n = 12 genomes
A
rhizosphere
0-2 CM
2-10 CM
10-20 CM
20-30 CM
50-60 CM
eigencoverage
0.0
0.2
0.4
0.6
Jun
Jul
Aug
Sep
Oct
time
B
Rhizosphere DNA coverage (%)
0.050
0.025
0.000
-0.025
-0.050
Bulk-soil DNA coverage (%)
0.025
0.050
0.075
0.100
Phylum
Acidobacteriota
Chloroflexota
Paceibacteria
Planctomycetota
Pseudomonadota
Verrucomicrobiota
RNA detected?
No
RNA coverage (%)
0
C
proportional abundance
1.00
0.75
0.50
0.25
0.00
May
Jun
Jul
Aug
Sep
Oct
time
Phylum
Acidobacteriota
Chloroflexota
Paceibacteria
Planctomycetota
Pseudomonadota
Verrucomicrobiota

**Fig. S20. Grey60 cohort abundance, activity, and taxonomic composition. (A)** Mean abundance profiles of cohort genomes across time and soil compartments. Error bars show standard errors across biological replicates. Blue shading indicates flooded time points. **(B)** Genome abundance in the rhizosphere compared with bulk soil, colored by phylum. Genomes with detectable RNA are shown as circles and scaled by whole-genome RNA coverage estimated with CoverM. Genomes without detectable RNA are shown as triangles. Genomes are numbered by decreasing abundance; full genome names are provided in Data S1. **(C)** Phylum-level cohort composition over time across soil compartments.

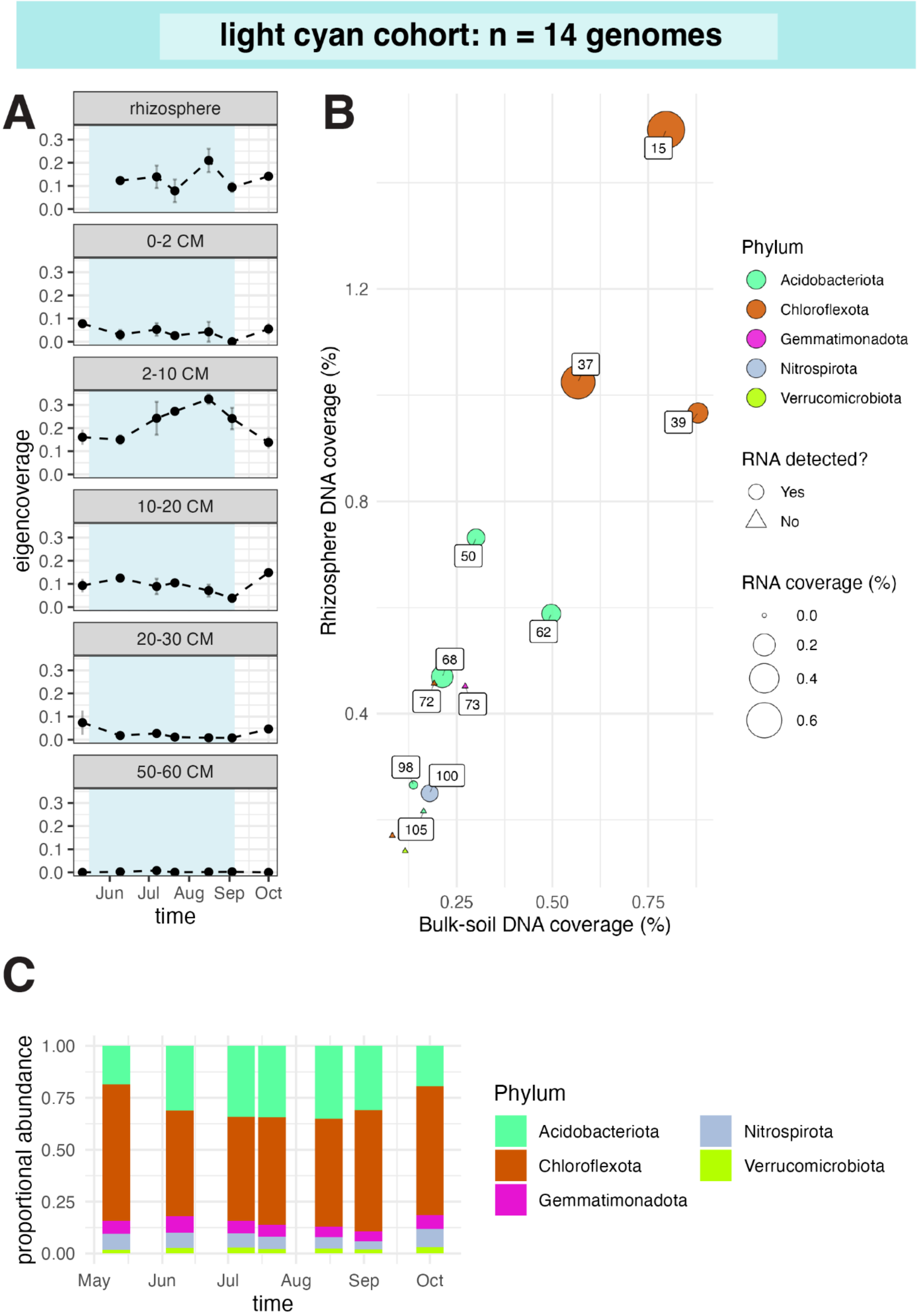

light cyan cohort: n = 14 genomes
A
rhizosphere
0-2 CM
2-10 CM
10-20 CM
20-30 CM
50-60 CM
0.0
0.1
0.2
0.3
eigencoverage
Jun
Jul
Aug
Sep
Oct
time
B
Rhizosphere DNA coverage (%)
0.4
0.8
1.2
Bulk-soil DNA coverage (%)
0.25
0.50
0.75
15
37
39
50
62
68
72
73
98
100
105
Phylum
Acidobacteriota
Chloroflexota
Gemmatimonadota
Nitrospirota
Verrucomicrobiota
RNA detected?
Yes
No
RNA coverage (%)
0.0
0.2
0.4
0.6
C
proportional abundance
0.00
0.25
0.50
0.75
1.00
May
Jun
Jul
Aug
Sep
Oct
time
Phylum
Acidobacteriota
Chloroflexota
Gemmatimonadota
Nitrospirota
Verrucomicrobiota

**Fig. S21. Light cyan cohort abundance, activity, and taxonomic composition. (A)** Mean abundance profiles of cohort genomes across time and soil compartments. Error bars show standard errors across biological replicates. Blue shading indicates flooded time points. **(B)** Genome abundance in the rhizosphere compared with bulk soil, colored by phylum. Genomes with detectable RNA are shown as circles and scaled by whole-genome RNA coverage estimated with CoverM. Genomes without detectable RNA are shown as triangles. Genomes are numbered by decreasing abundance; full genome names are provided in Data S1. **(C)** Phylum-level cohort

composition over time across soil compartments.

lightgreen cohort: n = 10 genomes

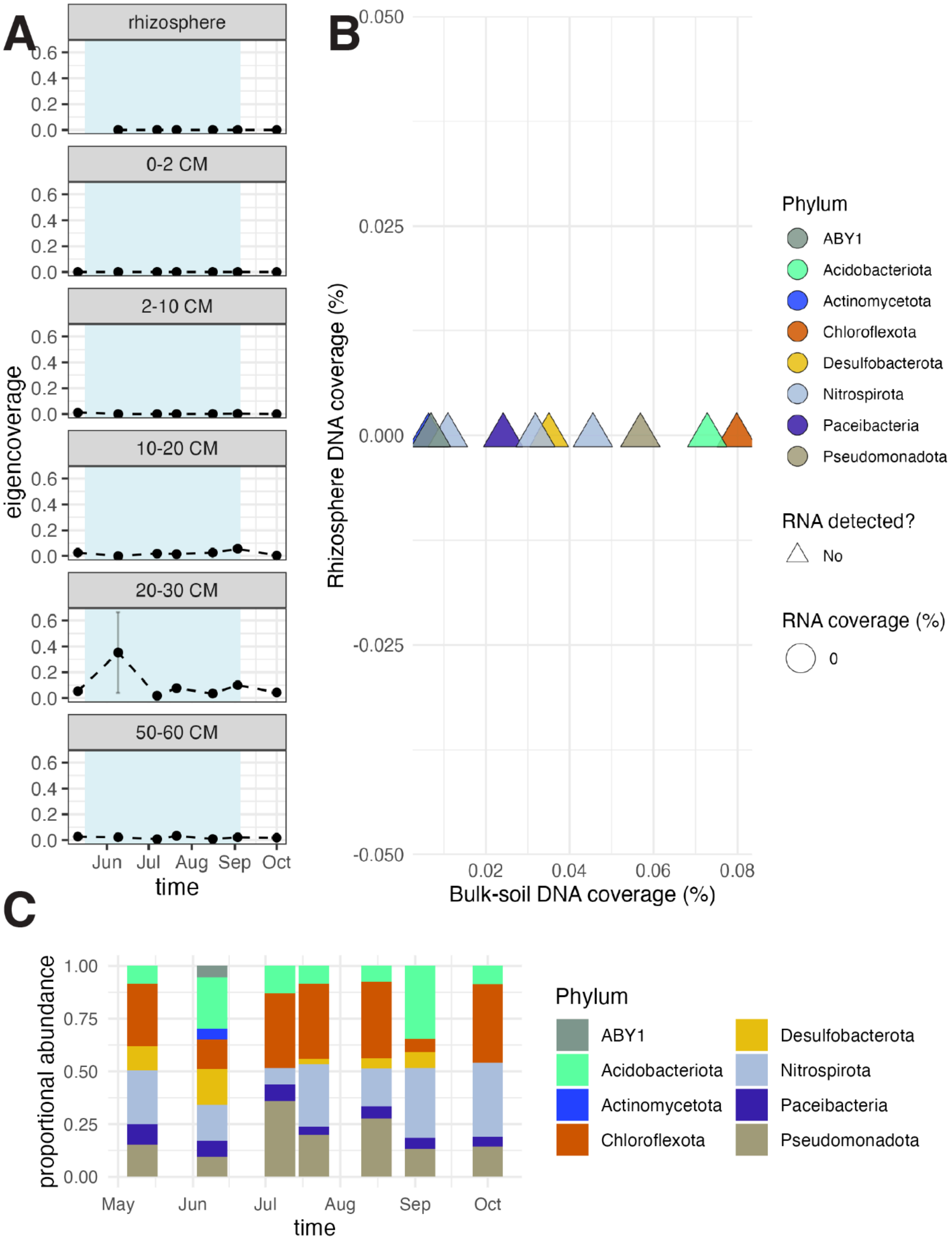

**Fig. S22. Lightgreen cohort abundance, activity, and taxonomic composition. (A)** Mean abundance profiles of cohort genomes across time and soil compartments. Error bars show standard errors across biological replicates. Blue shading indicates flooded time points. **(B)** Genome abundance in the rhizosphere compared with bulk soil, colored by phylum. Genomes with detectable RNA are shown as circles and scaled by whole-genome RNA coverage estimated with CoverM. Genomes without detectable RNA are shown as triangles. Genomes are numbered by decreasing abundance; full genome names are provided in Data S1. **(C)** Phylum-level cohort composition over time across soil compartments.

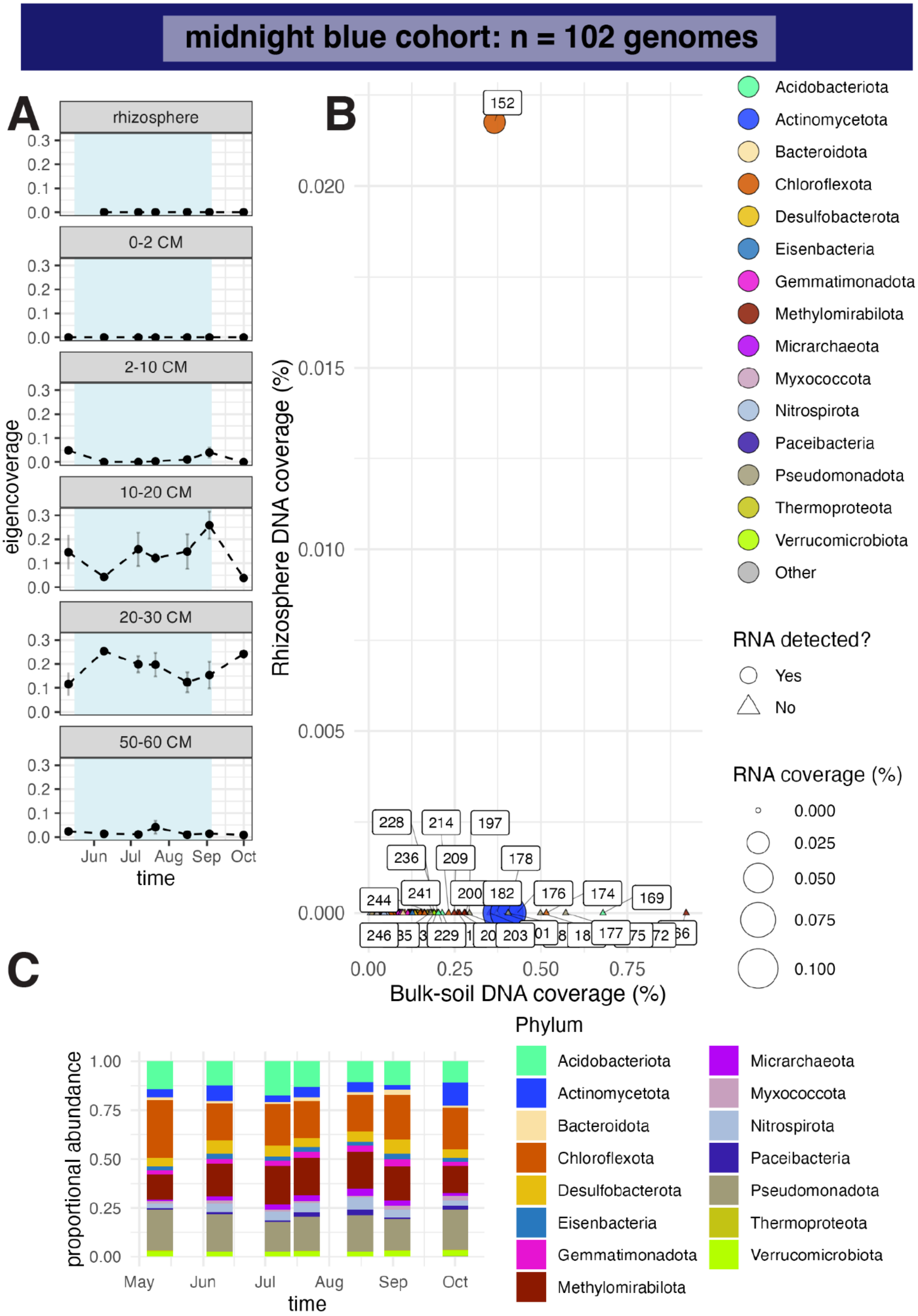

midnight blue cohort: n = 102 genomes
A
rhizosphere
0-2 CM
2-10 CM
10-20 CM
20-30 CM
50-60 CM
eigencoverage
Jun Jul Aug Sep Oct
time
B
Rhizosphere DNA coverage (%)
Bulk-soil DNA coverage (%)
Acidobacteriota
Actinomycetota
Bacteroidota
Chloroflexota
Desulfobacterota
Eisenbacteria
Gemmatimonadota
Methylomirabilota
Micrarchaeota
Myxococcota
Nitrospirota
Paceibacteria
Pseudomonadota
Thermoproteota
Verrucomicrobiota
Other
RNA detected?
Yes
No
RNA coverage (%)
C
proportional abundance
May Jun Jul Aug Sep Oct
time
Phylum

**Fig. S23. Midnight blue cohort abundance, activity, and taxonomic composition. (A)** Mean abundance profiles of cohort genomes across time and soil compartments. Error bars show standard errors across biological replicates. Blue shading indicates flooded time points. **(B)** Genome abundance in the rhizosphere compared with bulk soil, colored by phylum. Genomes with detectable RNA are shown as circles and scaled by whole-genome RNA coverage estimated with CoverM. Genomes without detectable RNA are shown as triangles. Genomes are numbered by decreasing abundance; full genome names are provided in Data S1. **(C)** Phylum-level cohort composition over time across soil compartments.

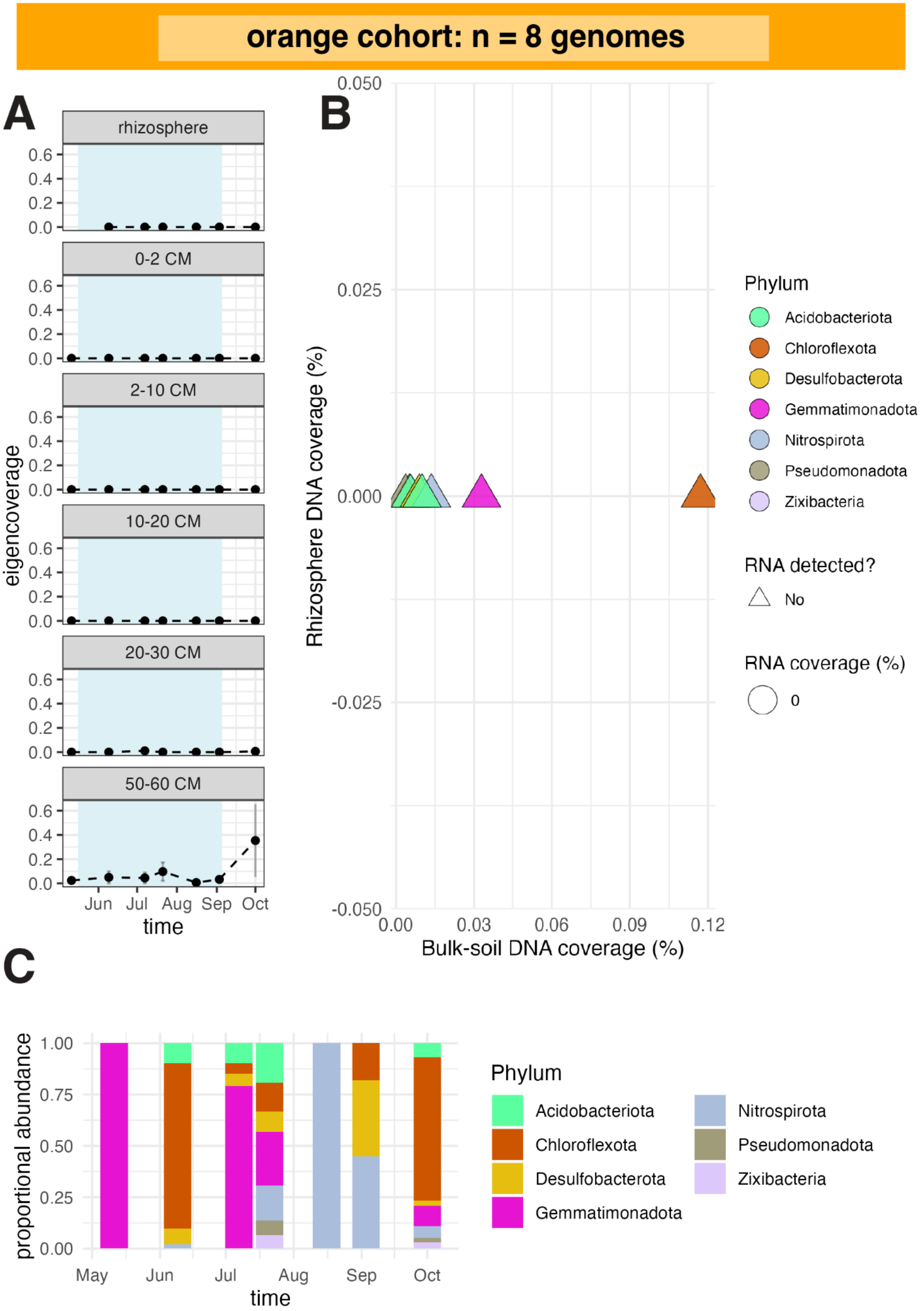
orange cohort: n = 8 genomes
A
rhizosphere
0-2 CM
2-10 CM
10-20 CM
20-30 CM
50-60 CM
eigencoverage
0.0
0.2
0.4
0.6
Jun
Jul
Aug
Sep
Oct
time
B
Rhizosphere DNA coverage (%)
0.050
0.025
0.000
-0.025
-0.050
0.00
0.03
0.06
0.09
0.12
Bulk-soil DNA coverage (%)
Phylum
Acidobacteriota
Chloroflexota
Desulfobacterota
Gemmatimonadota
Nitrospirota
Pseudomonadota
Zixibacteria
RNA detected?
No
RNA coverage (%)
0
C
proportional abundance
0.00
0.25
0.50
0.75
1.00
May
Jun
Jul
Aug
Sep
Oct
time
Phylum
Acidobacteriota
Chloroflexota
Desulfobacterota
Gemmatimonadota
Nitrospirota
Pseudomonadota
Zixibacteria

**Fig. S24. Orange cohort abundance, activity, and taxonomic composition. (A)** Mean abundance profiles of cohort genomes across time and soil compartments. Error bars show standard errors across biological replicates. Blue shading indicates flooded time points. **(B)** Genome abundance in the rhizosphere compared with bulk soil, colored by phylum. Genomes with detectable RNA are shown as circles and scaled by whole-genome RNA coverage estimated with CoverM. Genomes without detectable RNA are shown as triangles. Genomes are numbered by decreasing abundance; full genome names are provided in Data S1. **(C)** Phylum-level cohort composition over time across soil compartments.

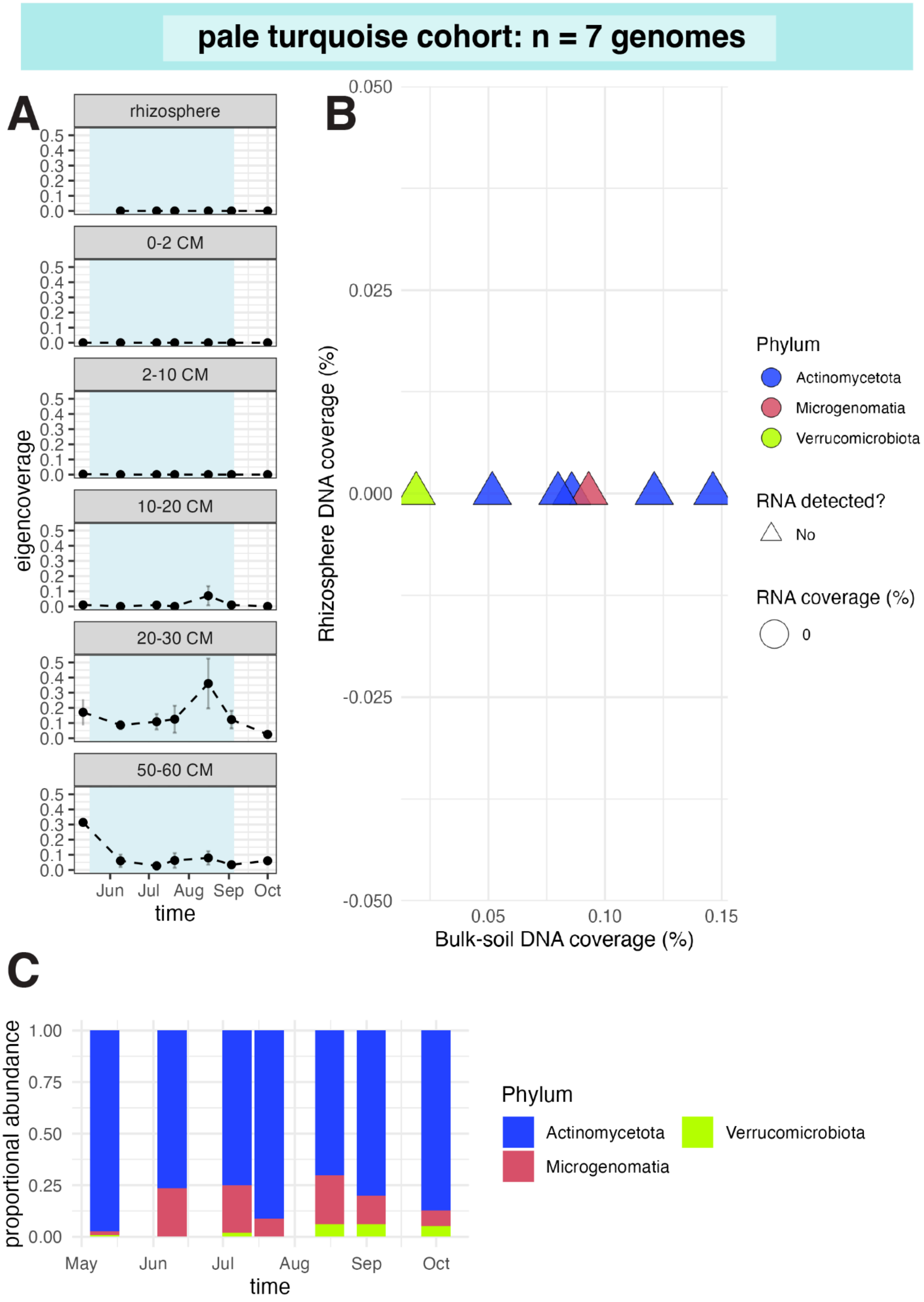
pale turquoise cohort: n = 7 genomes
A
rhizosphere
0-2 CM
2-10 CM
10-20 CM
20-30 CM
50-60 CM
eigencoverage
0.0 0.1 0.2 0.3 0.4 0.5
Jun Jul Aug Sep Oct
time
B
Rhizosphere DNA coverage (%)
0.050 0.025 0.000 -0.025 -0.050
0.05 0.10 0.15
Bulk-soil DNA coverage (%)
Phylum
Actinomycetota
Microgenomatia
Verrucomicrobiota
RNA detected?
No
RNA coverage (%)
0
C
proportional abundance
1.00 0.75 0.50 0.25 0.00
May Jun Jul Aug Sep Oct
time
Phylum
Actinomycetota
Microgenomatia
Verrucomicrobiota

**Fig. S25. Pale turquoise cohort abundance, activity, and taxonomic composition. (A)** Mean abundance profiles of cohort genomes across time and soil compartments. Error bars show standard errors across biological replicates. Blue shading indicates flooded time points. **(B)** Genome abundance in the rhizosphere compared with bulk soil, colored by phylum. Genomes with detectable RNA are shown as circles and scaled by whole-genome RNA coverage estimated with CoverM. Genomes without detectable RNA are shown as triangles. Genomes are numbered by decreasing abundance; full genome names are provided in Data S1. **(C)** Phylum-level cohort composition over time across soil compartments.

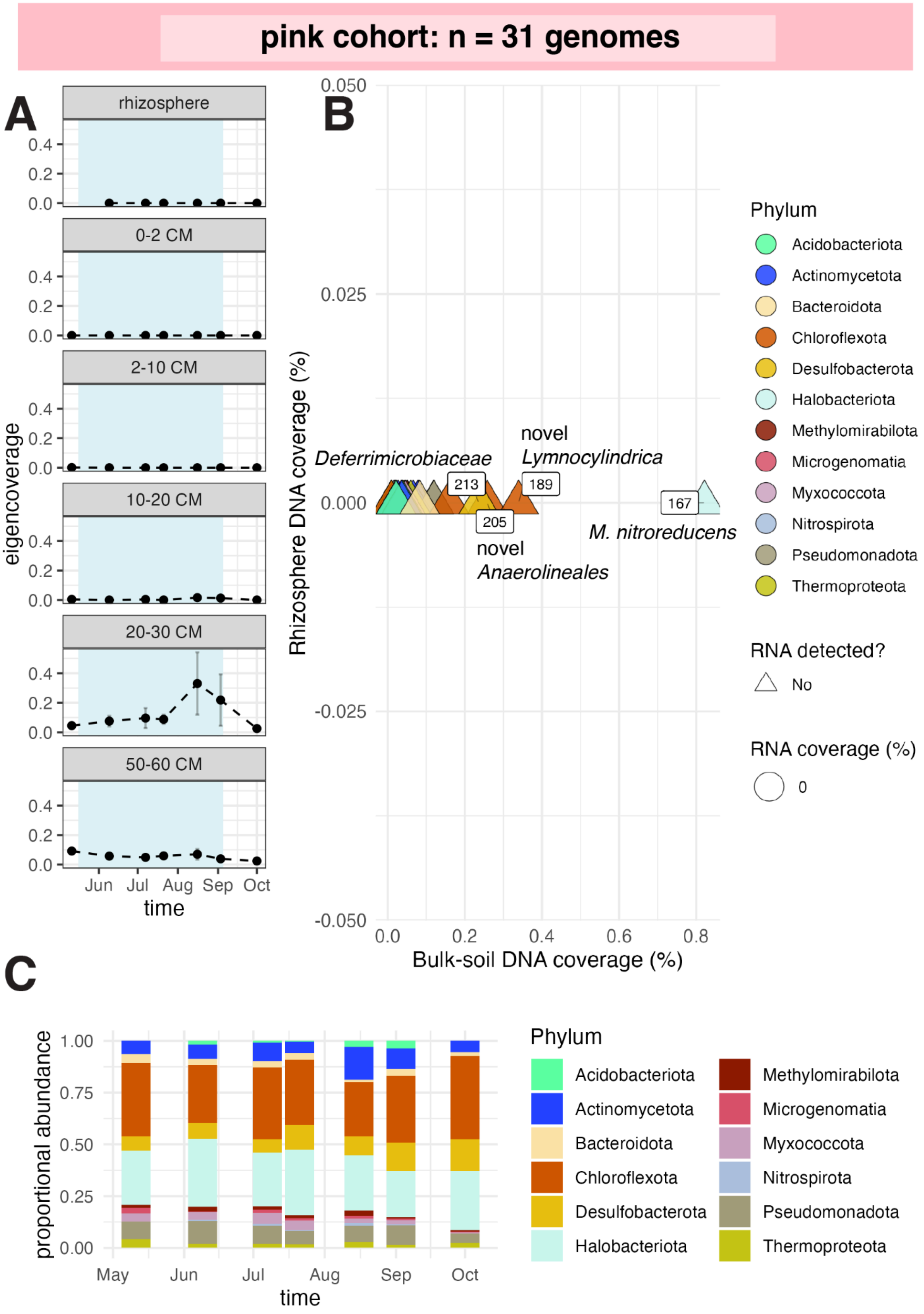
pink cohort: n = 31 genomes
A
rhizosphere
0-2 CM
2-10 CM
10-20 CM
20-30 CM
50-60 CM
eigencoverage
0.0
0.2
0.4
Jun
Jul
Aug
Sep
Oct
time
B
Rhizosphere DNA coverage (%)
0.050
0.025
0.000
-0.025
-0.050
Bulk-soil DNA coverage (%)
0.0
0.2
0.4
0.6
0.8
Deferrimicrobiaceae
novel
Lymnocylindrica
novel
Anaerolineales
M. nitroreducens
213
189
205
167
Phylum
Acidobacteriota
Actinomycetota
Bacteroidota
Chloroflexota
Desulfobacterota
Halobacteriota
Methylomirabilota
Microgenomatia
Myxococcota
Nitrospirota
Pseudomonadota
Thermoproteota
RNA detected?
No
RNA coverage (%)
0
C
proportional abundance
0.00
0.25
0.50
0.75
1.00
May
Jun
Jul
Aug
Sep
Oct
time
Phylum
Acidobacteriota
Actinomycetota
Bacteroidota
Chloroflexota
Desulfobacterota
Halobacteriota
Methylomirabilota
Microgenomatia
Myxococcota
Nitrospirota
Pseudomonadota
Thermoproteota

**Fig. S26. Pink cohort abundance, activity, and taxonomic composition. (A)** Mean abundance profiles of cohort genomes across time and soil compartments. Error bars show standard errors across biological replicates. Blue shading indicates flooded time points. **(B)** Genome abundance in the rhizosphere compared with bulk soil, colored by phylum. Genomes with detectable RNA are shown as circles and scaled by whole-genome RNA coverage estimated with CoverM. Genomes without detectable RNA are shown as triangles. Genomes are numbered by decreasing abundance; full genome names are provided in Data S1. **(C)** Phylum-level cohort composition over time across soil compartments.

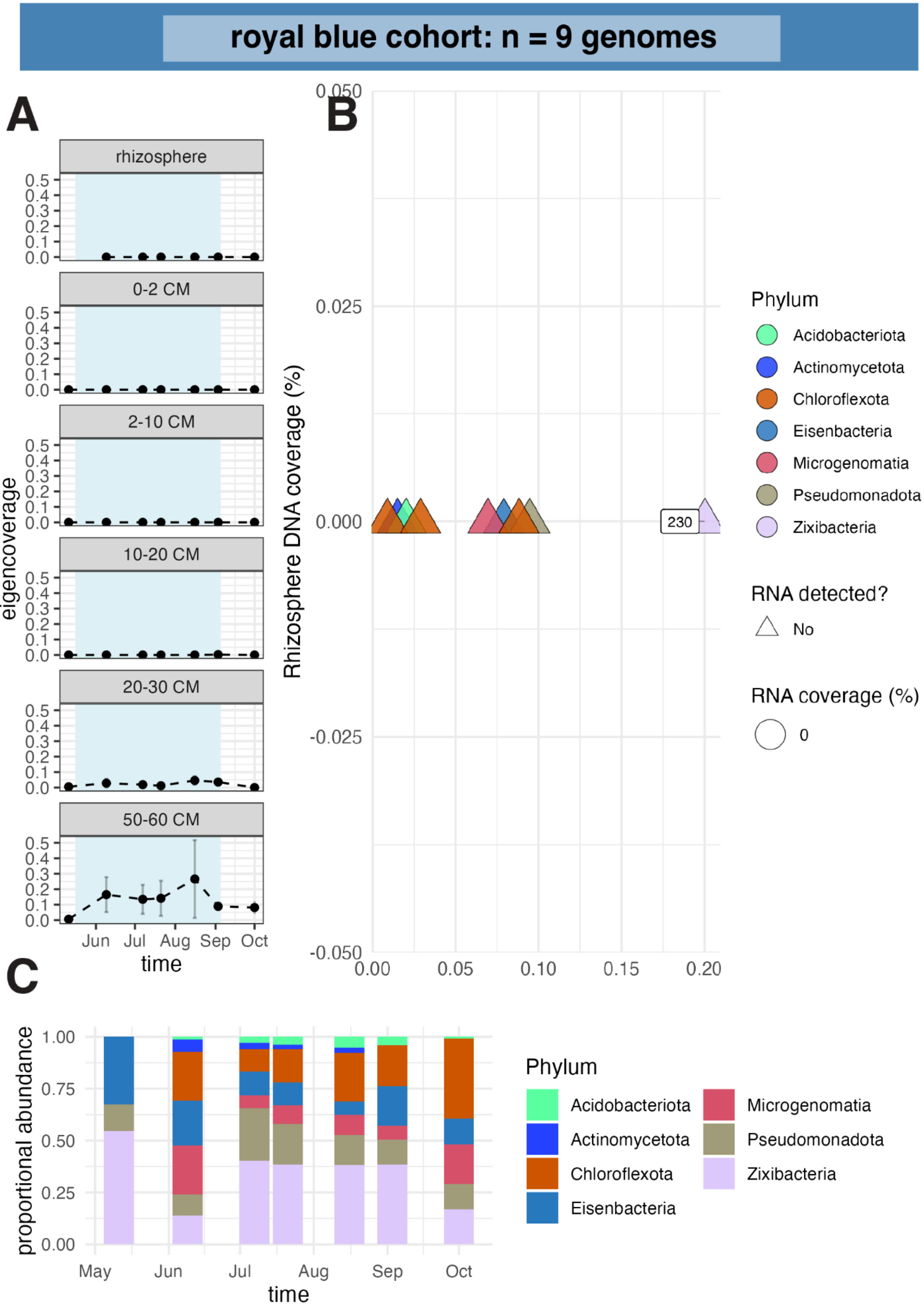
royal blue cohort: n = 9 genomes
A
B
C
rhizosphere
0-2 CM
2-10 CM
10-20 CM
20-30 CM
50-60 CM
eigencoverage
time
Jun Jul Aug Sep Oct
Rhizosphere DNA coverage (%)
230
Phylum
Acidobacteriota
Actinomycetota
Chloroflexota
Eisenbacteria
Microgenomatia
Pseudomonadota
Zixibacteria
RNA detected?
No
RNA coverage (%)
0
proportional abundance
May Jun Jul Aug Sep Oct
time

**Fig. S27. Royal blue cohort abundance, activity, and taxonomic composition. (A)** Mean abundance profiles of cohort genomes across time and soil compartments. Error bars show standard errors across biological replicates. Blue shading indicates flooded time points. **(B)** Genome abundance in the rhizosphere compared with bulk soil, colored by phylum. Genomes with detectable RNA are shown as circles and scaled by whole-genome RNA coverage estimated with CoverM. Genomes without detectable RNA are shown as triangles. Genomes are numbered by decreasing abundance; full genome names are provided in Data S1. **(C)** Phylum-level cohort composition over time across soil compartments.

### saddle brown cohort: n = 7 genomes

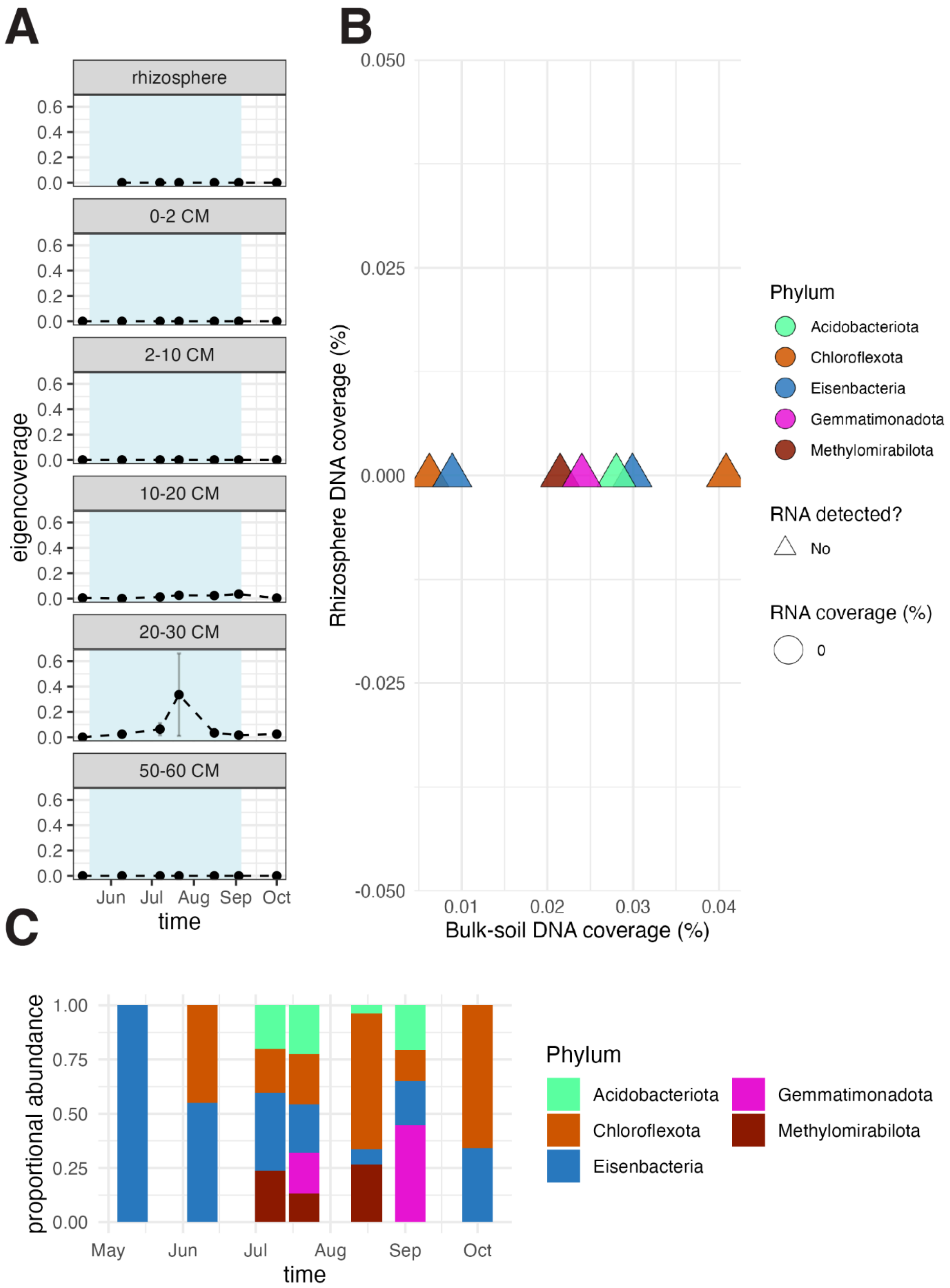

**Fig. S28. Saddle brown cohort abundance, activity, and taxonomic composition. (A)** Mean abundance profiles of cohort genomes across time and soil compartments. Error bars show standard errors across biological replicates. Blue shading indicates flooded time points. **(B)** Genome abundance in the rhizosphere compared with bulk soil, colored by phylum. Genomes with detectable RNA are shown as circles and scaled by whole-genome RNA coverage estimated with CoverM. Genomes without detectable RNA are shown as triangles. Genomes are numbered by decreasing abundance; full genome names are provided in Data S1. **(C)** Phylum-level cohort composition over time across soil compartments.

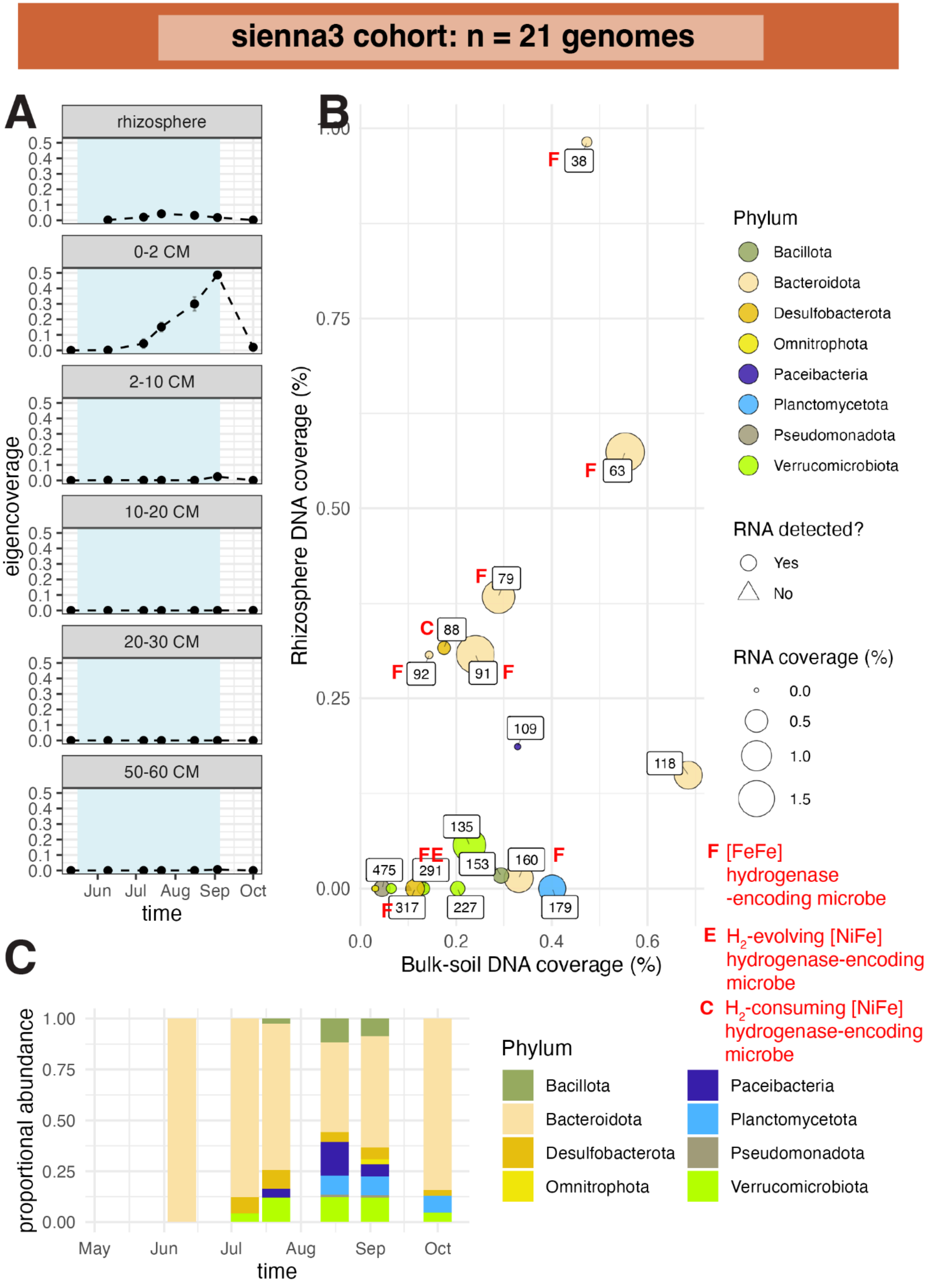

sienna3 cohort: n = 21 genomes
A
rhizosphere
0-2 CM
2-10 CM
10-20 CM
20-30 CM
50-60 CM
eigencoverage
Jun Jul Aug Sep Oct
time
B
Rhizosphere DNA coverage (%)
Bulk-soil DNA coverage (%)
38 63 79 88 92 91 109 118 135 475 291 153 160 317 227 179
Phylum
Bacillota
Bacteroidota
Desulfobacterota
Omnitrophota
Paceibacteria
Planctomycetota
Pseudomonadota
Verrucomicrobiota
RNA detected?
Yes
No
RNA coverage (%)
F [FeFe] hydrogenase -encoding microbe
E $H_2$-evolving [NiFe] hydrogenase-encoding microbe
C $H_2$-consuming [NiFe] hydrogenase-encoding microbe
C
proportional abundance
May Jun Jul Aug Sep Oct
time

**Fig. S29. Sienna3 cohort abundance, activity, and taxonomic composition. (A)** Mean abundance profiles of cohort genomes across time and soil compartments. Error bars show standard errors across biological replicates. Blue shading indicates flooded time points. **(B)** Genome abundance in the rhizosphere compared with bulk soil, colored by phylum. Genomes with detectable RNA are shown as circles and scaled by whole-genome RNA coverage estimated with CoverM. Genomes without detectable RNA are shown as triangles. Genomes are numbered by decreasing abundance; full genome names are provided in Data S1. **(C)** Phylum-level cohort composition over time across soil compartments.

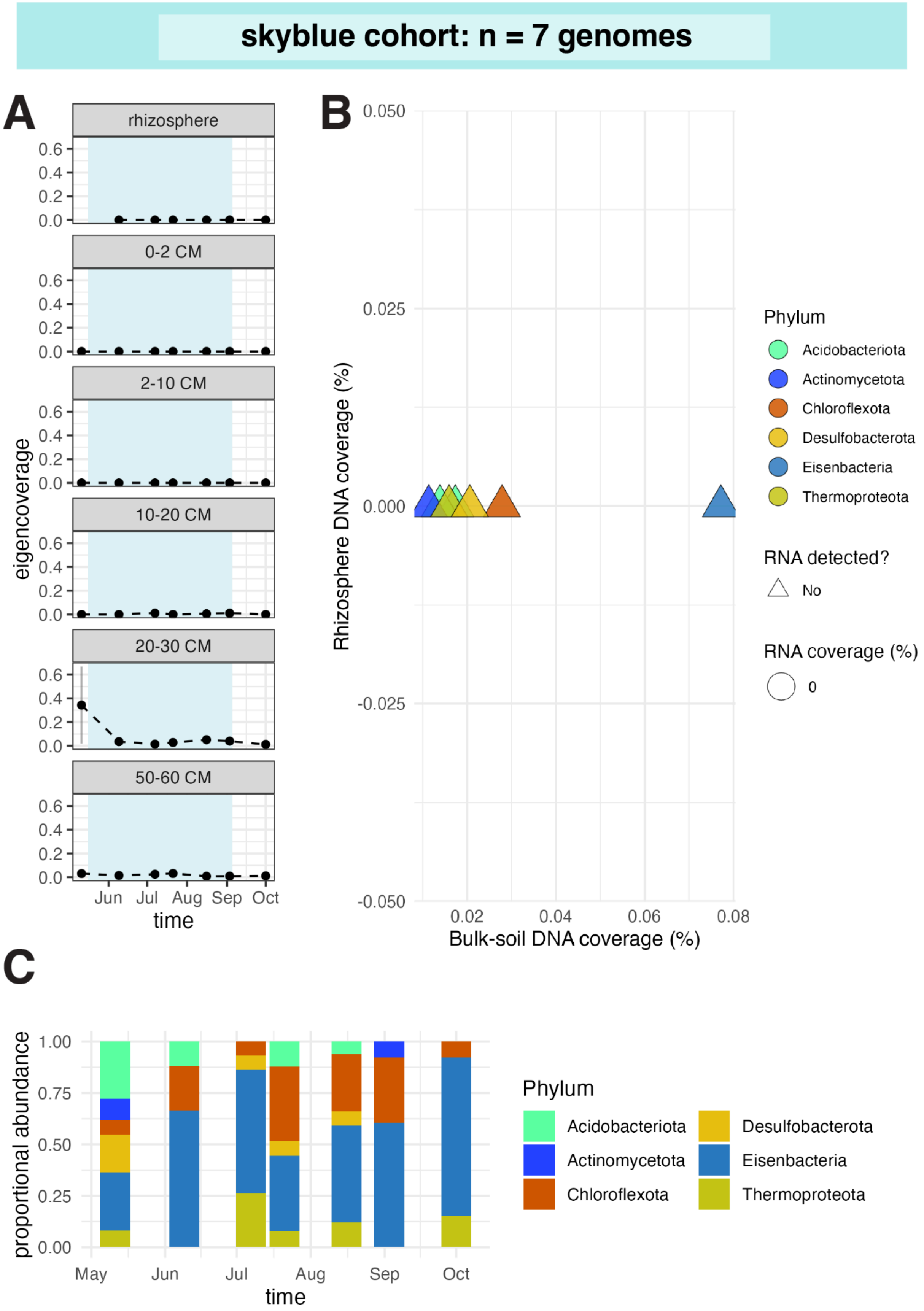
skyblue cohort: n = 7 genomes
A
rhizosphere
0-2 CM
2-10 CM
10-20 CM
20-30 CM
50-60 CM
eigencoverage
0.6
0.4
0.2
0.0
Jun Jul Aug Sep Oct
time
B
Rhizosphere DNA coverage (%)
0.050
0.025
0.000
-0.025
-0.050
0.02 0.04 0.06 0.08
Bulk-soil DNA coverage (%)
Phylum
Acidobacteriota
Actinomycetota
Chloroflexota
Desulfobacterota
Eisenbacteria
Thermoproteota
RNA detected?
No
RNA coverage (%)
0
C
proportional abundance
1.00
0.75
0.50
0.25
0.00
May Jun Jul Aug Sep Oct
time
Phylum
Acidobacteriota
Actinomycetota
Chloroflexota
Desulfobacterota
Eisenbacteria
Thermoproteota

**Fig. S30. Skyblue cohort abundance, activity, and taxonomic composition. (A)** Mean abundance profiles of cohort genomes across time and soil compartments. Error bars show standard errors across biological replicates. Blue shading indicates flooded time points. **(B)** Genome abundance in the rhizosphere compared with bulk soil, colored by phylum. Genomes with detectable RNA are shown as circles and scaled by whole-genome RNA coverage estimated with CoverM. Genomes without detectable RNA are shown as triangles. Genomes are numbered by decreasing abundance; full genome names are provided in Data S1. **(C)** Phylum-level cohort composition over time across soil compartments.

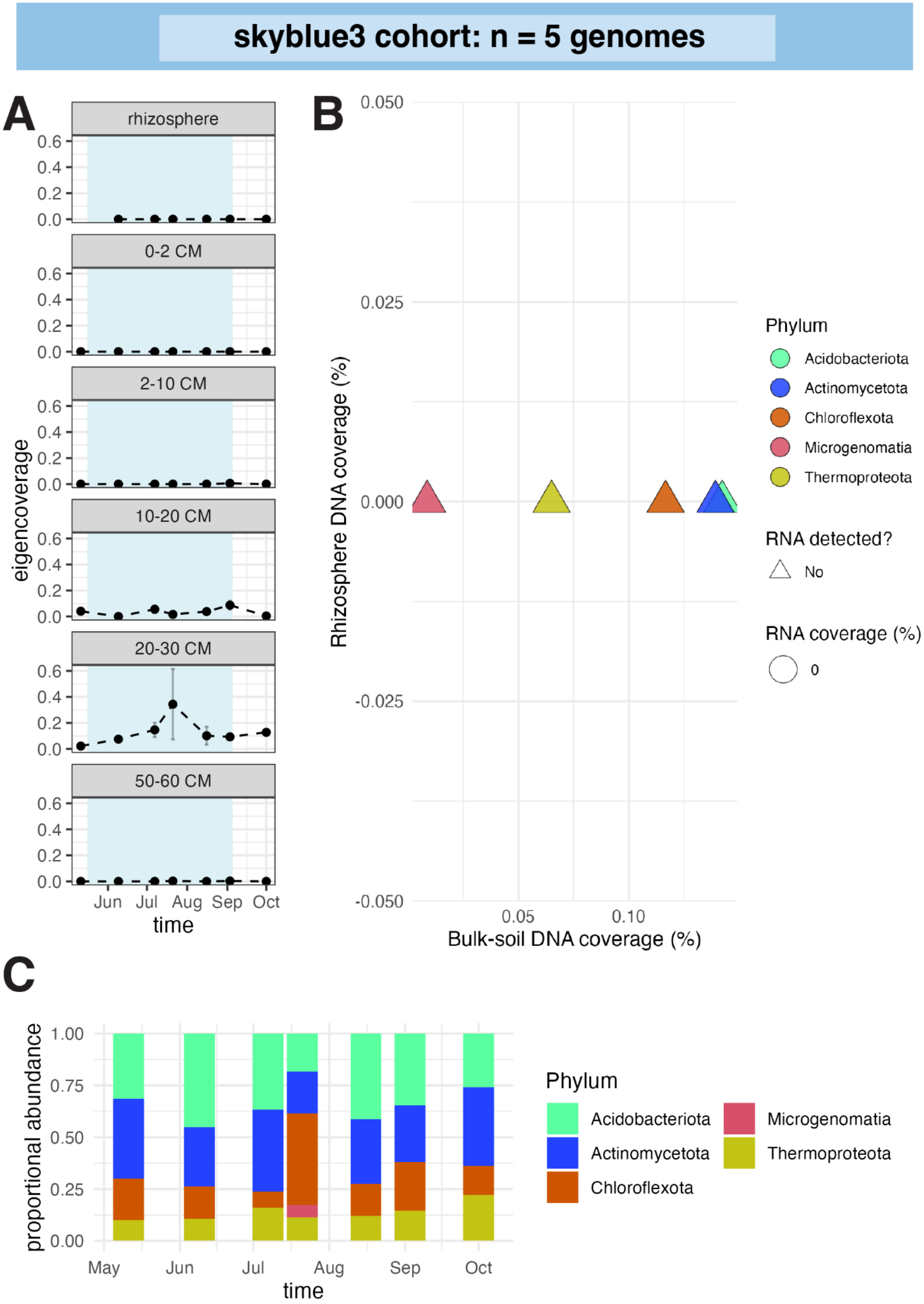

skyblue3 cohort: n = 5 genomes
A
rhizosphere
0-2 CM
2-10 CM
10-20 CM
20-30 CM
50-60 CM
eigencoverage
0.0
0.2
0.4
0.6
Jun
Jul
Aug
Sep
Oct
time
B
Rhizosphere DNA coverage (%)
0.050
0.025
0.000
-0.025
-0.050
0.05
0.10
Bulk-soil DNA coverage (%)
Phylum
Acidobacteriota
Actinomycetota
Chloroflexota
Microgenomatia
Thermoproteota
RNA detected?
No
RNA coverage (%)
0
C
proportional abundance
0.00
0.25
0.50
0.75
1.00
May
Jun
Jul
Aug
Sep
Oct
time
Phylum
Acidobacteriota
Actinomycetota
Chloroflexota
Microgenomatia
Thermoproteota

**Fig. S31. Skyblue3 cohort abundance, activity, and taxonomic composition. (A)** Mean abundance profiles of cohort genomes across time and soil compartments. Error bars show standard errors across biological replicates. Blue shading indicates flooded time points. **(B)** Genome abundance in the rhizosphere compared with bulk soil, colored by phylum. Genomes with detectable RNA are shown as circles and scaled by whole-genome RNA coverage estimated with CoverM. Genomes without detectable RNA are shown as triangles. Genomes are numbered by decreasing abundance; full genome names are provided in Data S1. **(C)** Phylum-level cohort composition over time across soil compartments.

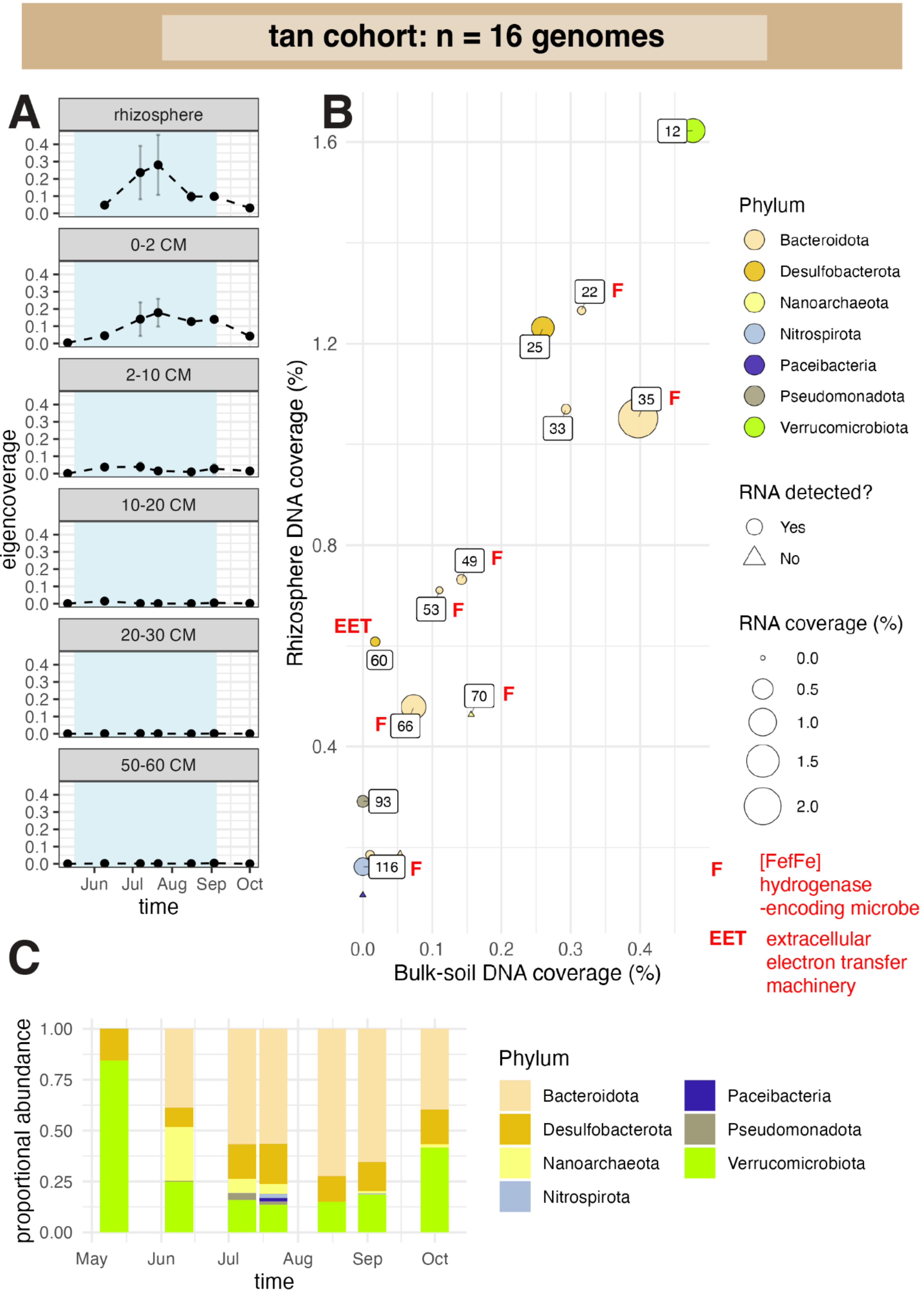

tan cohort: n = 16 genomes
A
rhizosphere
0-2 CM
2-10 CM
10-20 CM
20-30 CM
50-60 CM
eigencoverage
0.0
0.1
0.2
0.3
0.4
Jun
Jul
Aug
Sep
Oct
time
B
Rhizosphere DNA coverage (%)
Bulk-soil DNA coverage (%)
1.6
1.2
0.8
0.4
0.0
0.1
0.2
0.3
0.4
12
22
25
33
35
49
53
60
66
70
93
116
F
EET
Phylum
Bacteroidota
Desulfobacterota
Nanoarchaeota
Nitrospirota
Paceibacteria
Pseudomonadota
Verrucomicrobiota
RNA detected?
Yes
No
RNA coverage (%)
0.0
0.5
1.0
1.5
2.0
F [FefFe] hydrogenase -encoding microbe
EET extracellular electron transfer machinery
C
proportional abundance
0.00
0.25
0.50
0.75
1.00
May
Jun
Jul
Aug
Sep
Oct
time
Phylum
Bacteroidota
Desulfobacterota
Nanoarchaeota
Nitrospirota
Paceibacteria
Pseudomonadota
Verrucomicrobiota

**Fig. S32. Tan cohort abundance, activity, and taxonomic composition. (A)** Mean abundance profiles of cohort genomes across time and soil compartments. Error bars show standard errors across biological replicates. Blue shading indicates flooded time points. **(B)** Genome abundance in the rhizosphere compared with bulk soil, colored by phylum. Genomes with detectable RNA are shown as circles and scaled by whole-genome RNA coverage estimated with CoverM. Genomes without detectable RNA are shown as triangles. Genomes are numbered by decreasing abundance; full genome names are provided in Data S1. **(C)** Phylum-level cohort composition over time across soil compartments.

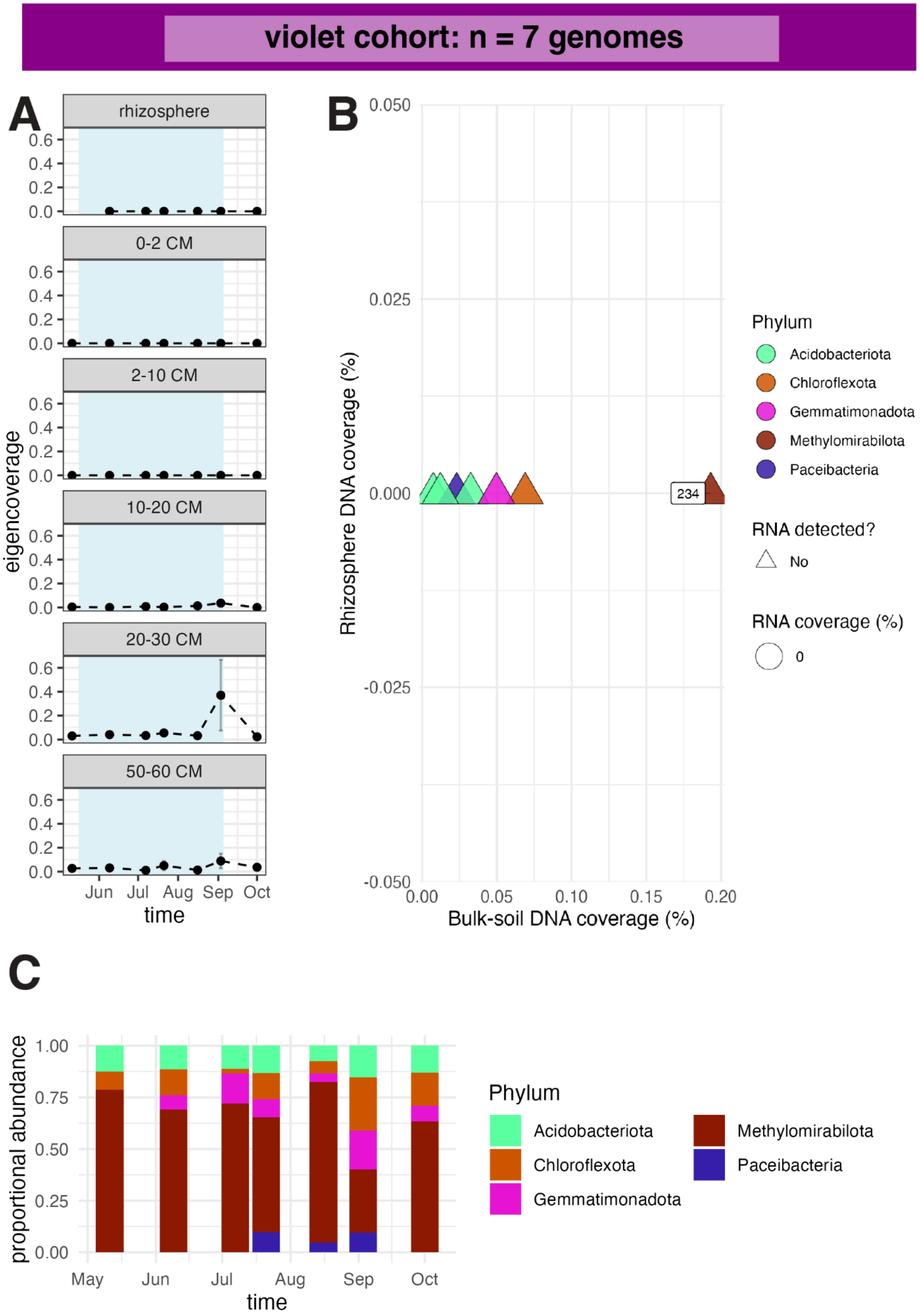
violet cohort: n = 7 genomes
A
rhizosphere
0-2 CM
2-10 CM
10-20 CM
20-30 CM
50-60 CM
0.6
0.4
0.2
0.0
eigencoverage
Jun
Jul
Aug
Sep
Oct
time
B
0.050
0.025
0.000
-0.025
-0.050
Rhizosphere DNA coverage (%)
234
0.00
0.05
0.10
0.15
0.20
Bulk-soil DNA coverage (%)
Phylum
Acidobacteriota
Chloroflexota
Gemmatimonadota
Methylomirabilota
Paceibacteria
RNA detected?
No
RNA coverage (%)
0
C
1.00
0.75
0.50
0.25
0.00
proportional abundance
May
Jun
Jul
Aug
Sep
Oct
time
Phylum
Acidobacteriota
Chloroflexota
Gemmatimonadota
Methylomirabilota
Paceibacteria

**Fig. S33. Violet cohort abundance, activity, and taxonomic composition. (A)** Mean abundance profiles of cohort genomes across time and soil compartments. Error bars show standard errors across biological replicates. Blue shading indicates flooded time points. **(B)** Genome abundance in the rhizosphere compared with bulk soil, colored by phylum. Genomes with detectable RNA are shown as circles and scaled by whole-genome RNA coverage estimated with CoverM. Genomes without detectable RNA are shown as triangles. Genomes are numbered by decreasing abundance; full genome names are provided in Data S1. **(C)** Phylum-level cohort composition over time across soil compartments.

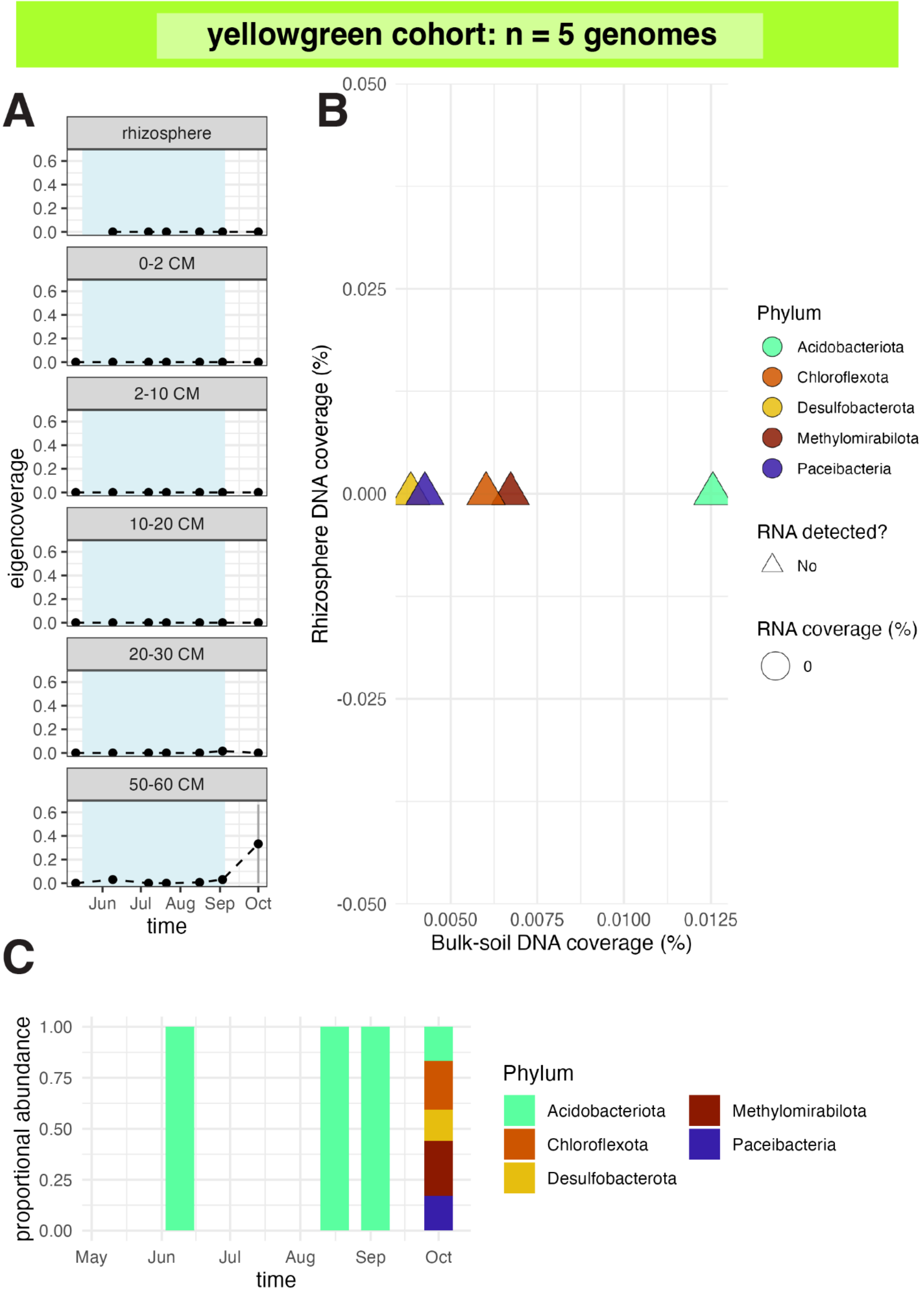

yellowgreen cohort: n = 5 genomes
A
rhizosphere
0-2 CM
2-10 CM
10-20 CM
20-30 CM
50-60 CM
eigencoverage
0.6
0.4
0.2
0.0
Jun Jul Aug Sep Oct
time
B
Rhizosphere DNA coverage (%)
0.050
0.025
0.000
-0.025
-0.050
0.0050 0.0075 0.0100 0.0125
Bulk-soil DNA coverage (%)
Phylum
Acidobacteriota
Chloroflexota
Desulfobacterota
Methylomirabilota
Paceibacteria
RNA detected?
No
RNA coverage (%)
0
C
proportional abundance
1.00
0.75
0.50
0.25
0.00
May Jun Jul Aug Sep Oct
time
Phylum
Acidobacteriota
Chloroflexota
Desulfobacterota
Methylomirabilota
Paceibacteria

**Fig. S34. Yellowgreen cohort abundance, activity, and taxonomic composition. (A)** Mean abundance profiles of cohort genomes across time and soil compartments. Error bars show standard errors across biological replicates. Blue shading indicates flooded time points. **(B)** Genome abundance in the rhizosphere compared with bulk soil, colored by phylum. Genomes with detectable RNA are shown as circles and scaled by whole-genome RNA coverage estimated with CoverM. Genomes without detectable RNA are shown as triangles. Genomes are numbered by decreasing abundance; full genome names are provided in Data S1. **(C)** Phylum-level cohort composition over time across soil compartments.

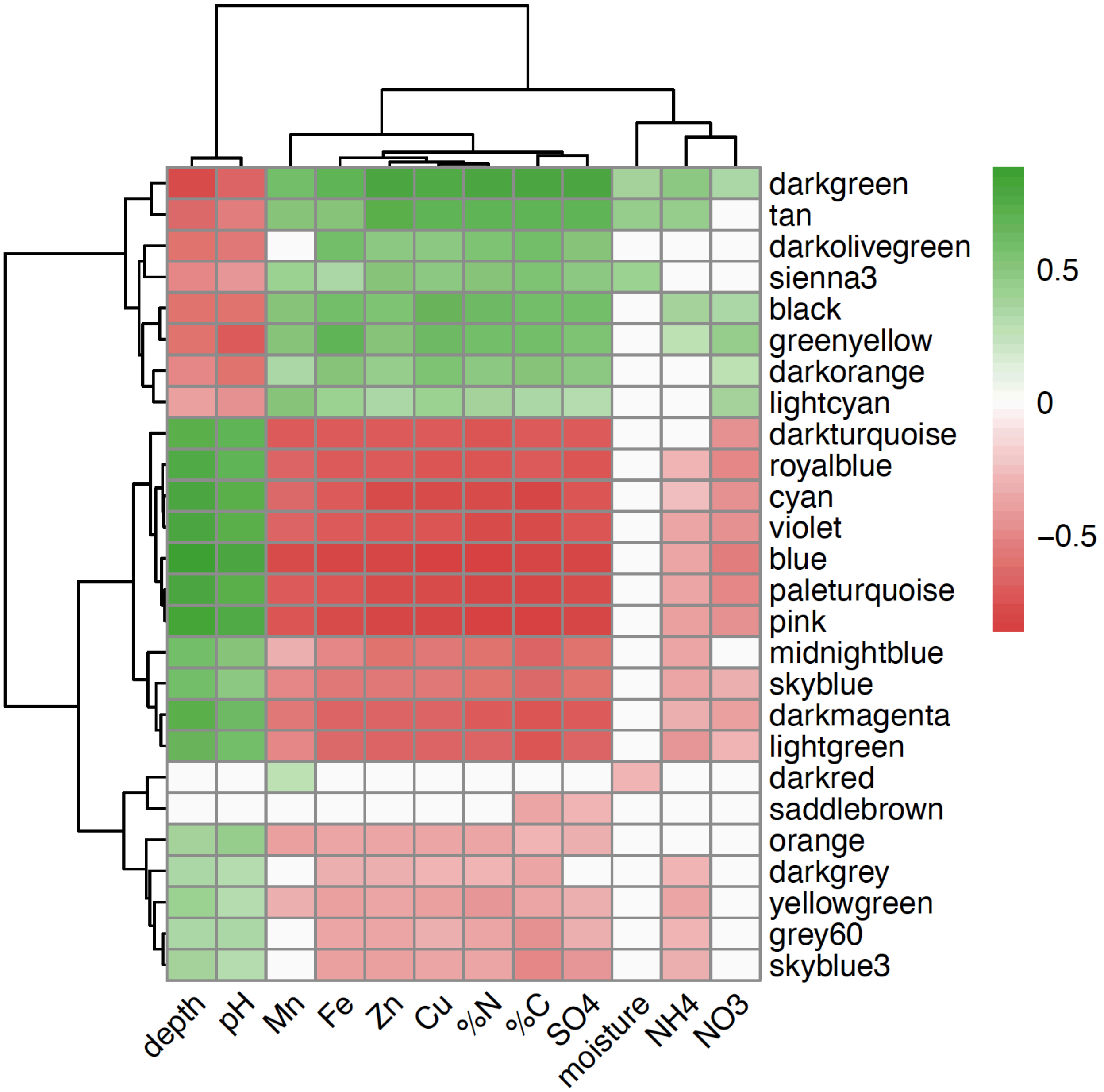


**Fig. S35. Heatmap of significant correlations between genome cohort average abundance patterns (eigenvalues) and environmental metadata.** Spearman rank correlations were calculated between WGCNA genome cohort eigenvalues (rows) and soil physicochemical and field variables (columns), with false discovery rate (FDR) correction applied across all tests (adjusted $p < 0.05$). Tile color denotes correlation strength and direction (green, positive; red, negative; white, near zero). Rows and columns were hierarchically clustered using Euclidean distance and complete linkage.

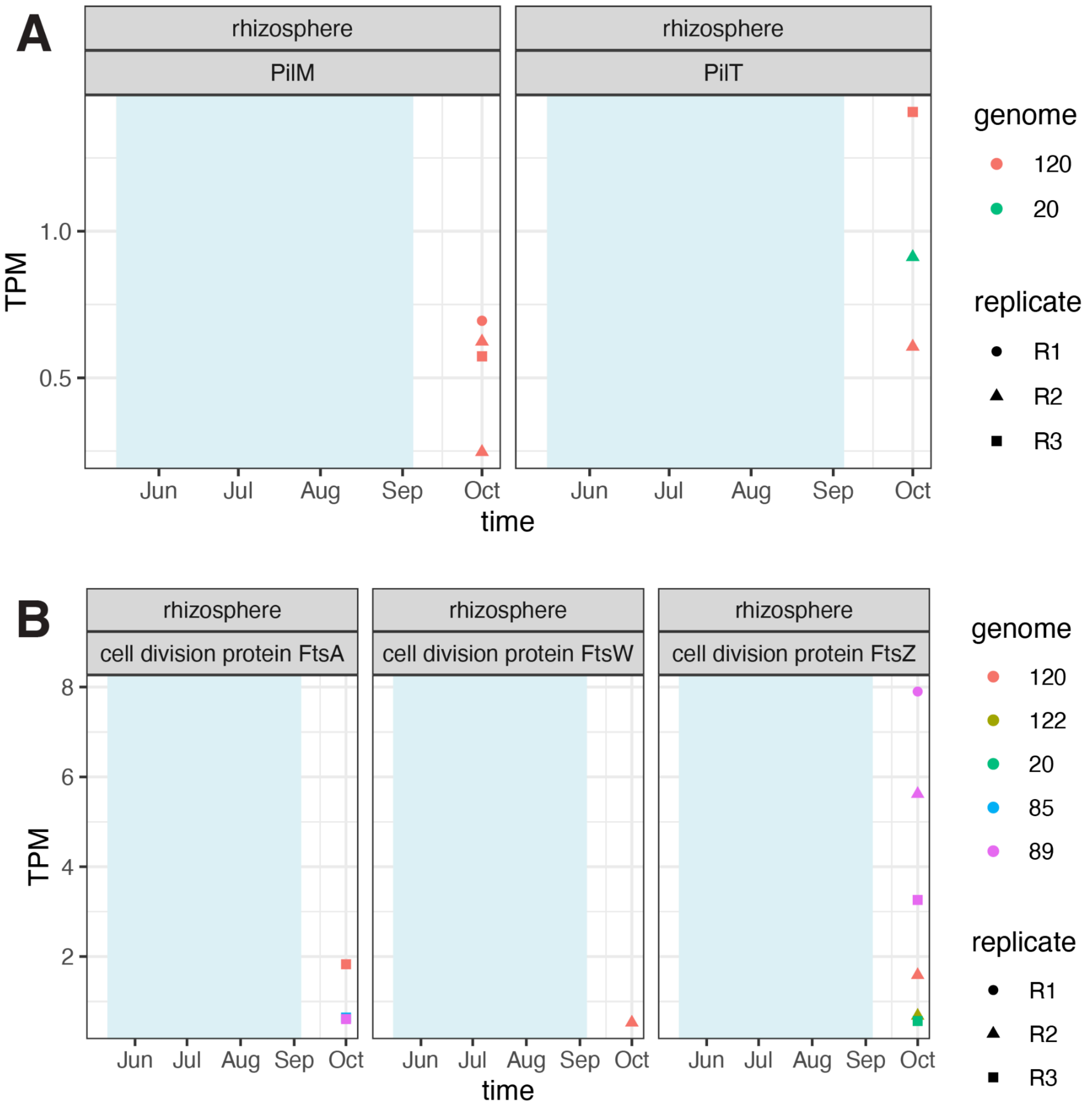


**Fig. S36. Darkorange cohort CPR expressed type IV pili (A) and cell division machinery (B) in the rhizosphere after drydown.** Expression of type IV pili and cell division machinery from darkorange cohort genomes (TPM= transcripts per million). Full genome information is in Data S2.

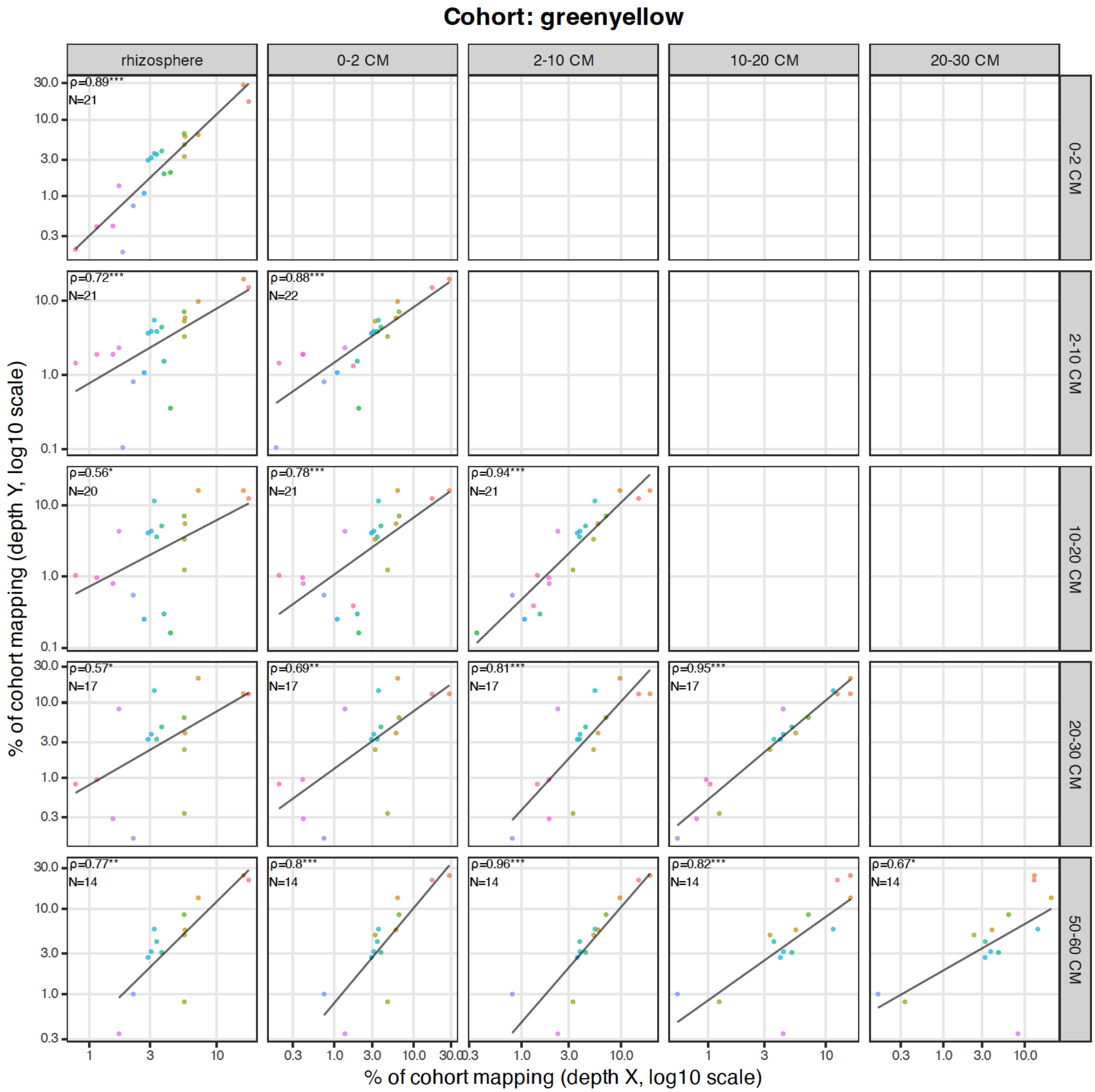


**Fig. S37. Pairwise comparisons of within-cohort genome abundance for greenyellow cohort across soil compartments.** Genome coverage was averaged across replicates and normalized within each cohort and depth to represent percent of cohort coverage. Points represent genomes and are colored by genome; axes are shown on log scales. Spearman's rank correlation coefficients (ρ) and sample sizes (N) are annotated per depth pair, with significance assessed using Benjamini–Hochberg FDR correction.

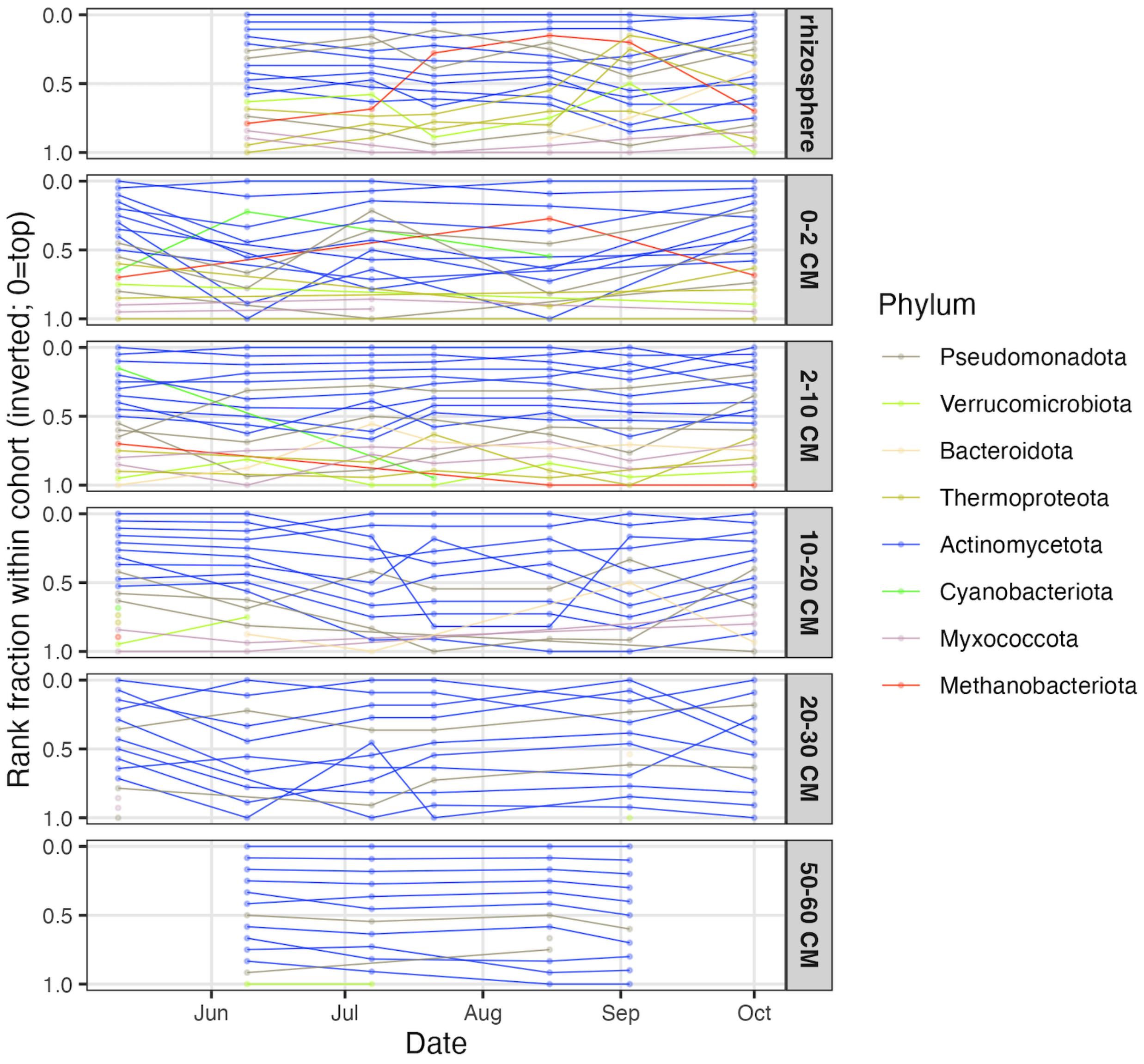


**Extended Data Fig. S38: Genome rank trajectories within greenyellow cohort across soil compartments.** Each line represents an individual genome assigned to the greenyellow cohort. At each sampling date and soil compartment, genomes were ranked by their relative contribution to total greenyellow cohort abundance, with lower values indicating higher rank (0 = most abundant genome). Colors indicate genome phylum. Panels show seasonal rank trajectories separately for the rhizosphere and bulk soil depth intervals.

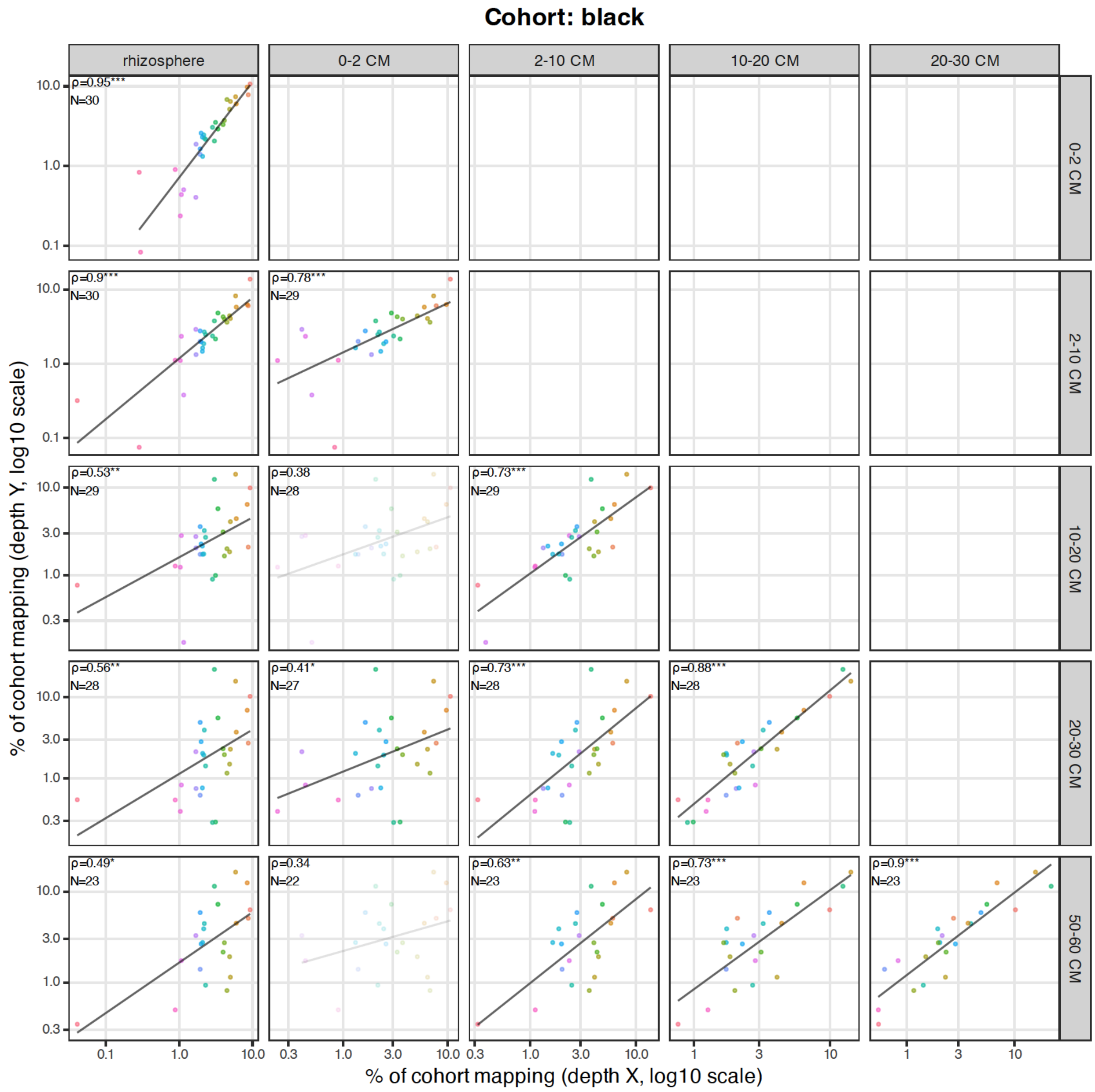


**Fig. S39. Pairwise comparisons of within-cohort genome abundance for black cohort across soil compartments.** Genome coverage was averaged across replicates and normalized within each cohort and depth to represent percent of cohort coverage. Points represent genomes and are colored by genome; axes are shown on log scales. Spearman's rank correlation coefficients (ρ) and sample sizes (N) are annotated per depth pair, with significance assessed using Benjamini–Hochberg FDR correction. Non-significant panels are visually de-emphasized.

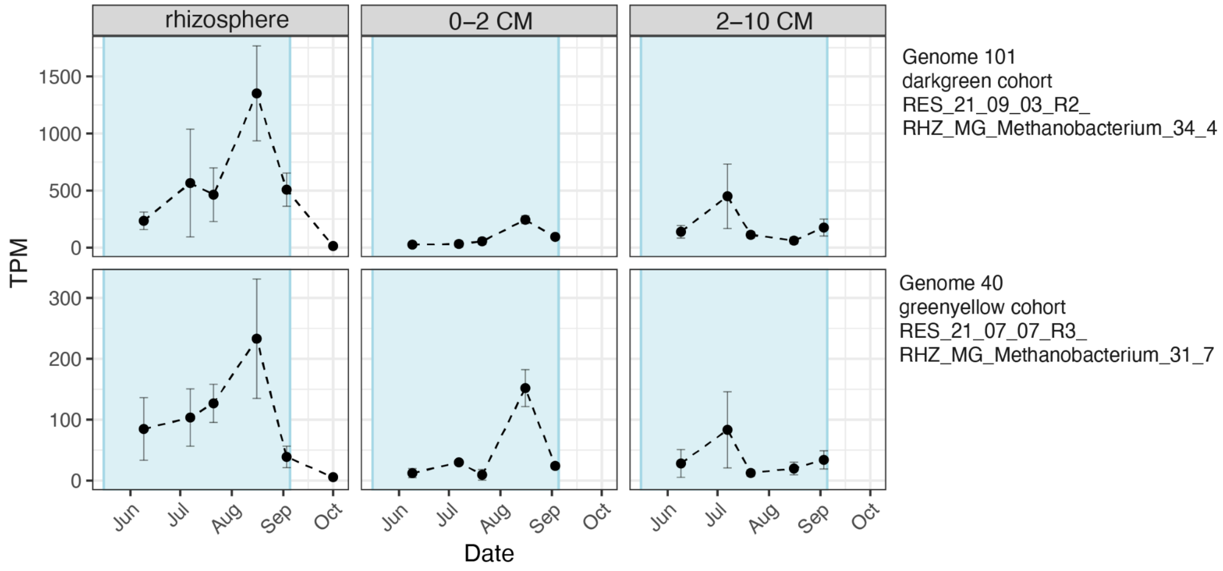


**Fig. S40. Seasonal Methyl-coenzyme M expression patterns are conserved across methanogen genomes.** Summed expression across all mcr subunits within the two recovered methanogen genomes. Expression was averaged across replicates and error bars represent standard error.

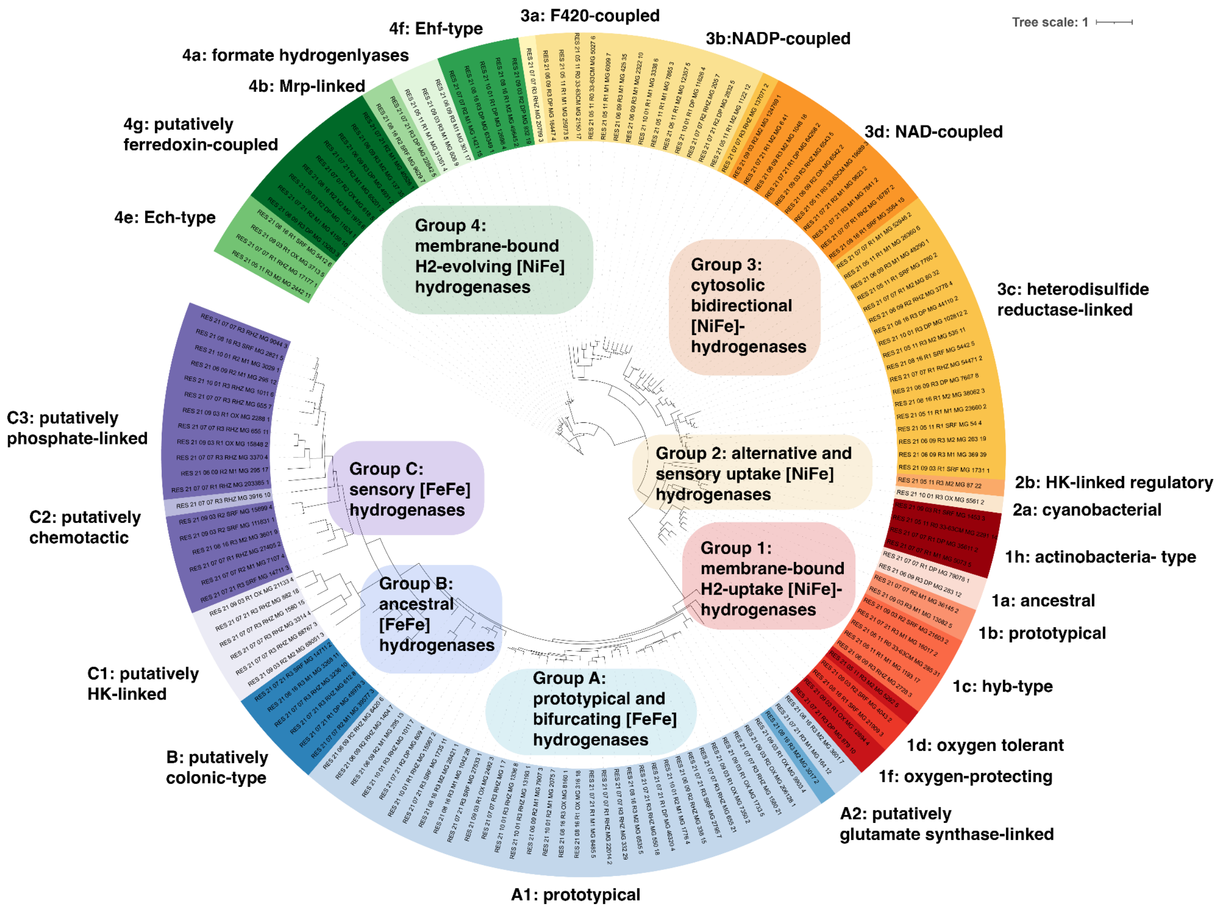


**Fig. S41. Maximum-likelihood phylogeny of hydrogenases detected across genomes, colored by hydrogenase functional group.** Hydrogenases were identified using HydDB HMMs[1], and classes were determined by constructing Maximum-likelihood phylogenies with HydDB references (see methods).

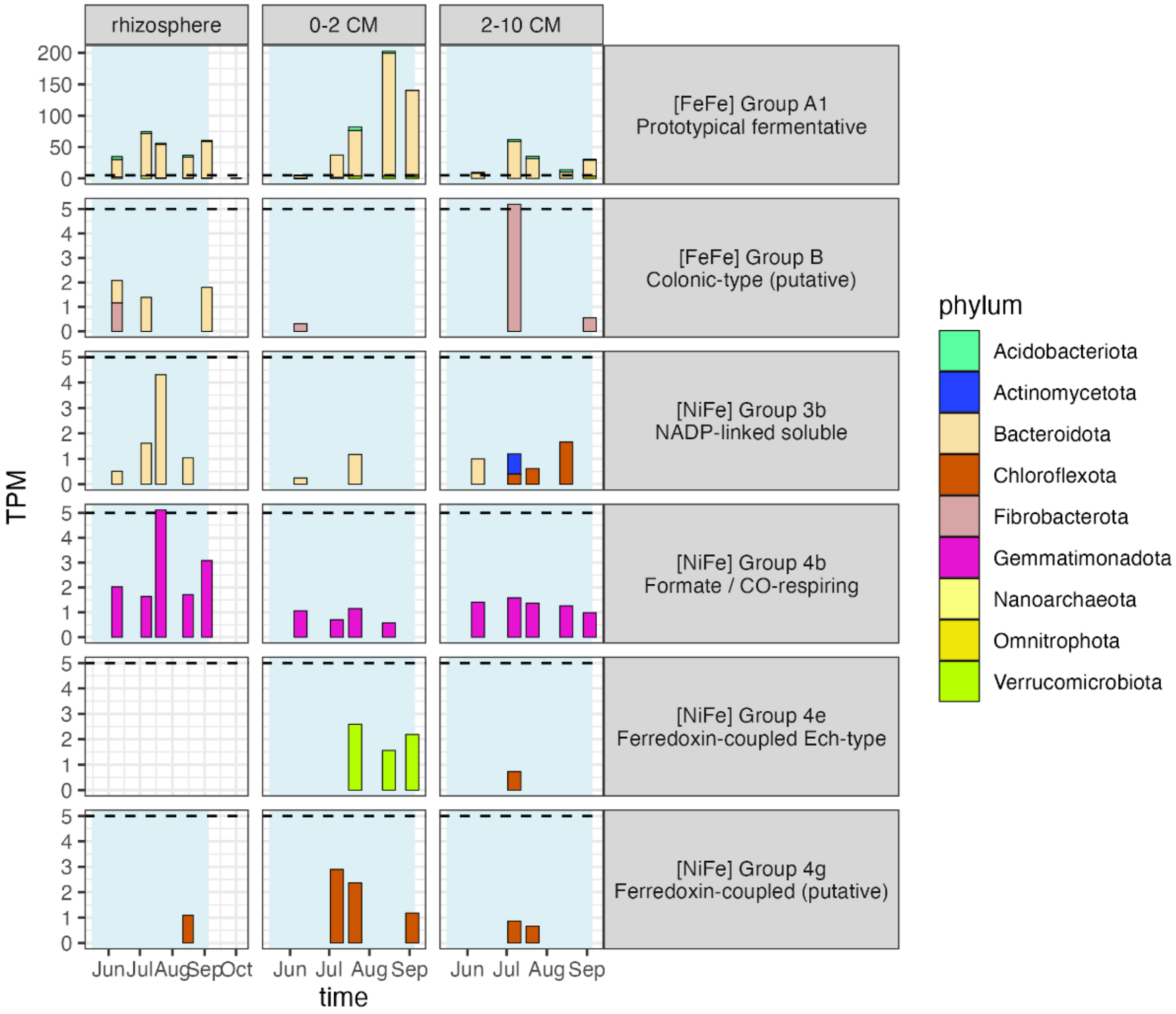


**Fig. S42. $H_2$-evolving hydrogenase expression by soil compartment, colored by genome taxonomy.** Barcharts showing expression over time of hydrogenase genes in the rhizosphere (first column row), 0-2 cm bulk soil (middle column) and 2-10 cm bulk soil (right column). Expression is separated by hydrogenase type (rows) and colored by genome phylum. Units are transcripts per million (TPM). Y-axis differ; horizontal dashed line is plotted at 5 TPM for perspective.

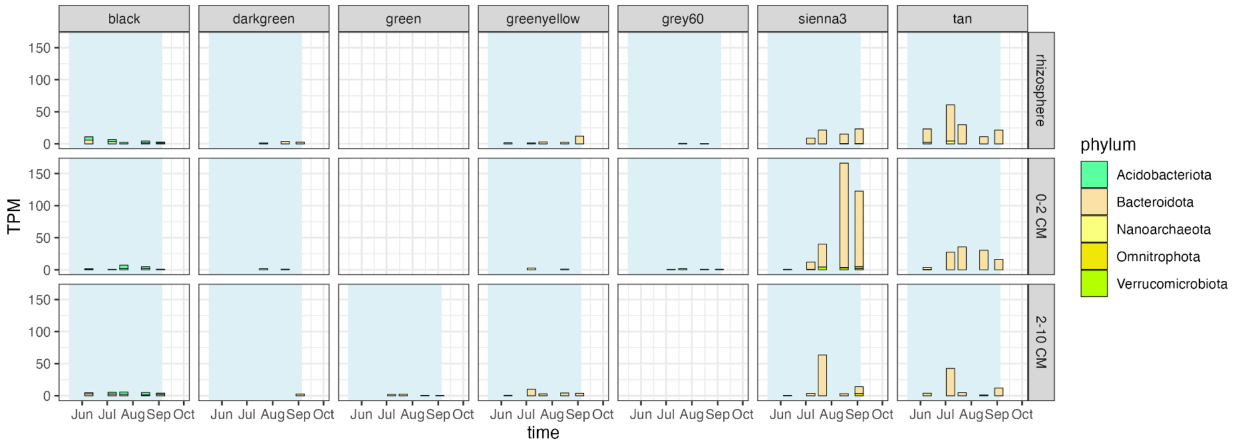


**Fig. S43. Fermentative hydrogenase expression by soil compartment and cohort, colored by genome taxonomy.** Barcharts showing expression over time of fermentative hydrogenase genes in the rhizosphere (top row), 0-2 cm bulk soil (middle row) and 2-10 cm bulk soil (bottom row). Expression is separated by genome cohort (columns) and colored by genome phylum. Units are transcripts per million (TPM).

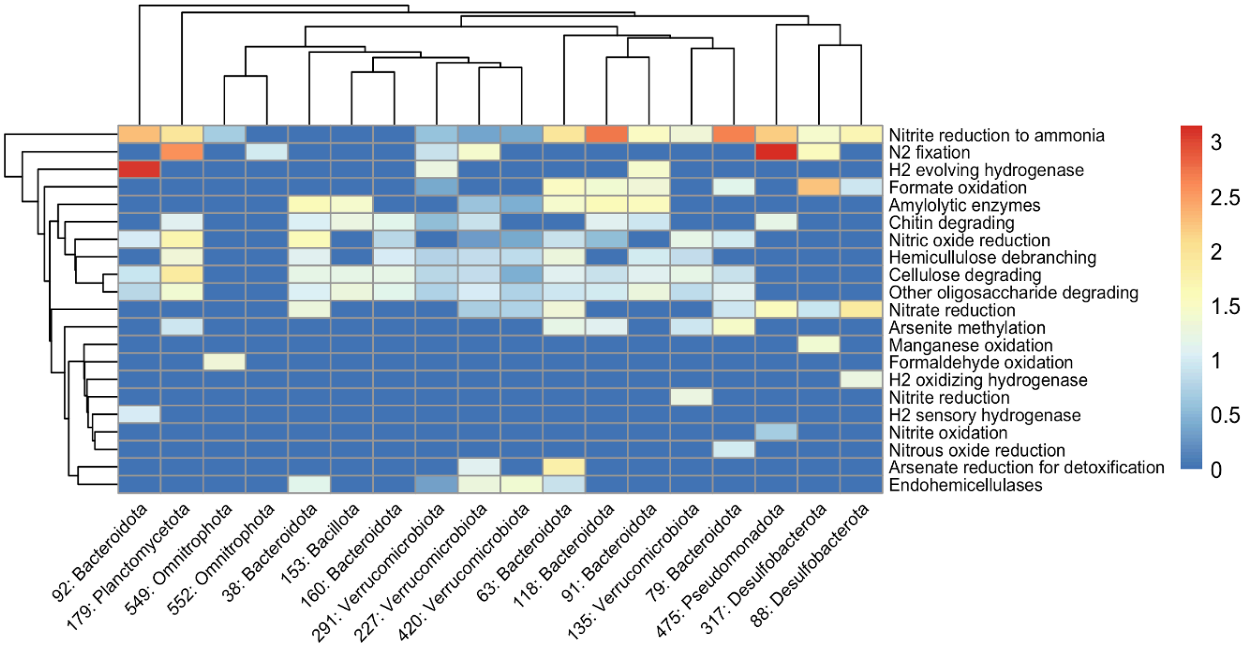


**Fig. S44. Expression of sienna3 cohort genes that may contribute to hydrogen cycling.** Columns are labeled by genome ID: Phylum (full details in Data S2); rows are gene functional annotation. Heatmap is colored by logTPM (warm colors are highly expressed, cool colors have low expression).

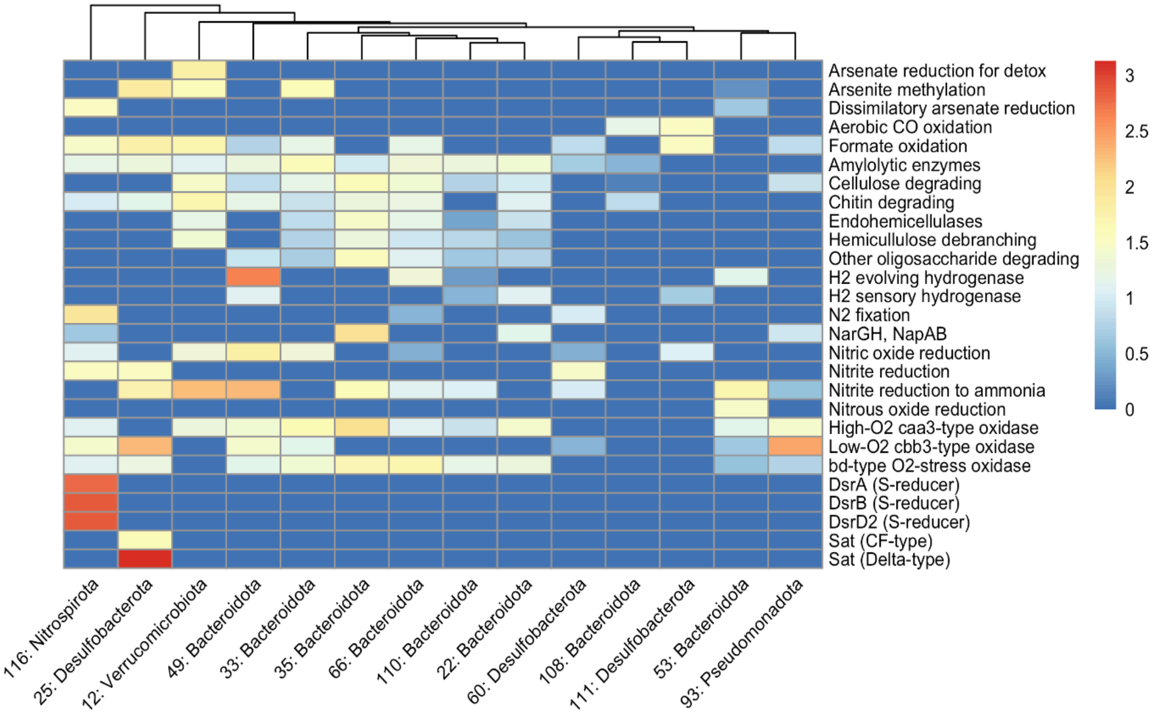

**Fig. S45. Expression of tan cohort genes that may contribute to hydrogen cycling.** Columns are labeled by genome ID: Phylum (full details in Data S2); rows are gene functional annotation. Heatmap is colored by logTPM (warm colors are highly expressed, cool colors have low expression).

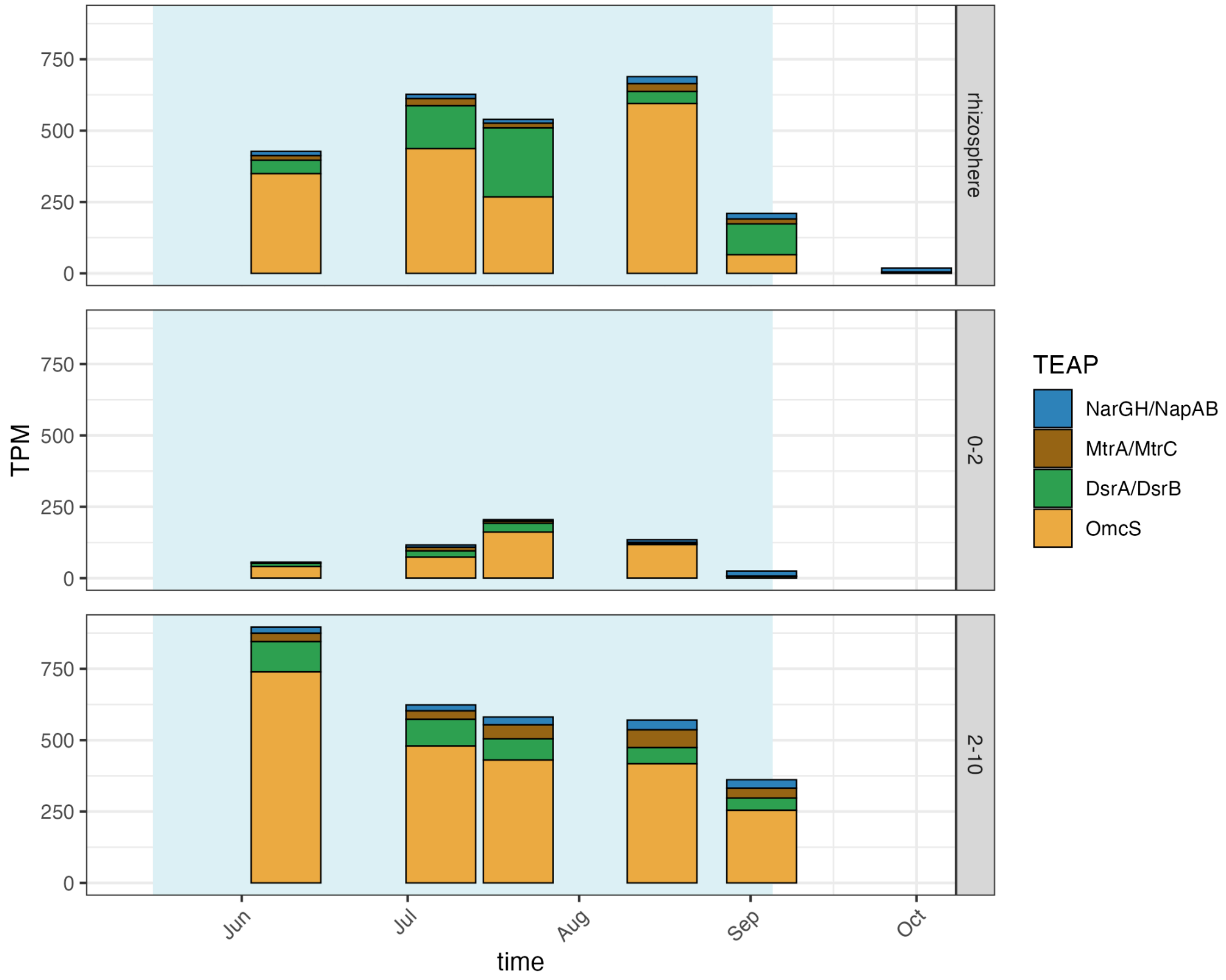


**Fig. S46. Summed TEAP expression over time by soil compartment from genome set.** TPM= transcripts per million. The blue rectangle represents flooded timepoints.

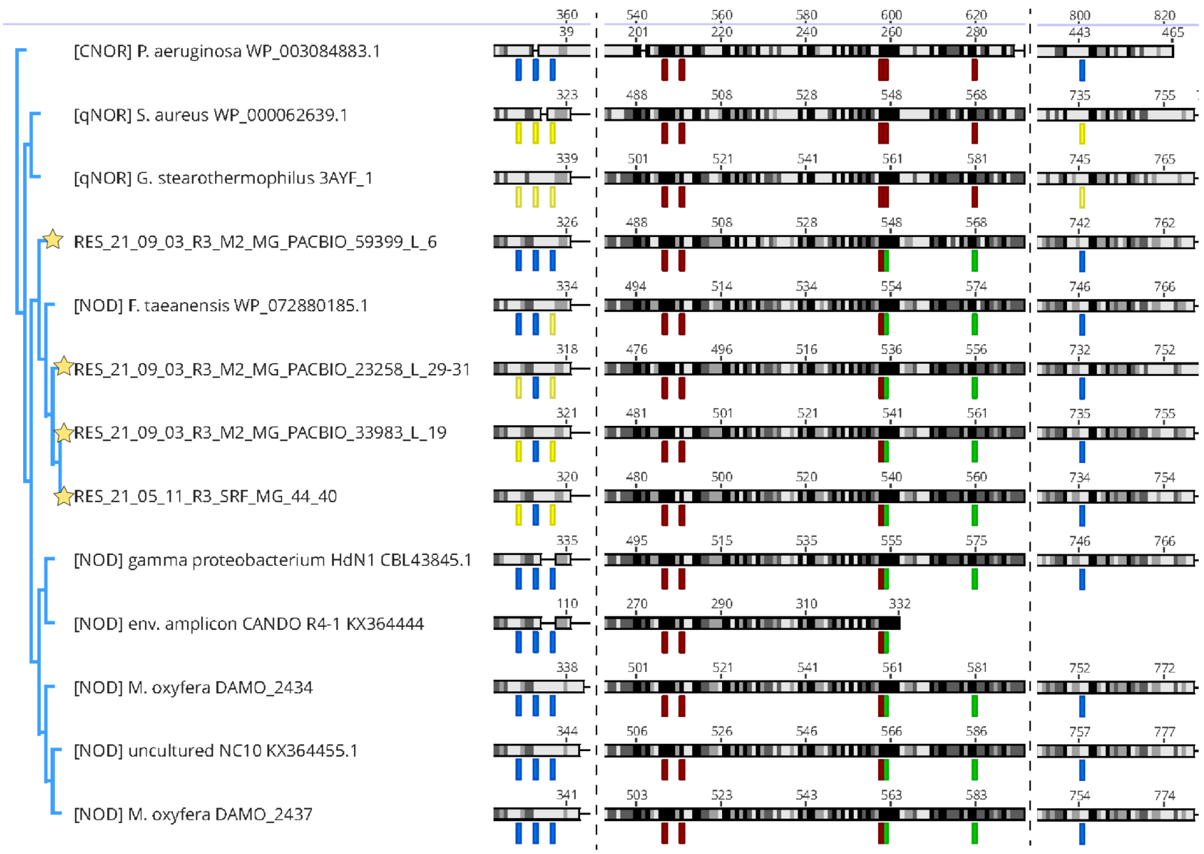


**Fig. S47. Phylogeny of putative NODs compared to qNORs.** Quinol-dependent nitric oxide reductases (qNORs) reduce nitric oxide during denitrification and are the closest characterized homologs of nitric oxide dismutases (NODs). Putative NODs from this study cluster with reference NOD sequences. Stars indicate proteins recovered in this study. Vertical dashed lines denote truncated regions removed for ease of viewing. Yellow bars indicate conserved qNOR quinol-binding residues; blue bars indicate substitutions at these positions. Like reference NODs, the putative NODs from this study have disrupted quinol-binding residues. Red bars indicate conserved qNOR active-site residues; green bars indicate substitutions at these positions. The putative NODs from this study share the same active-site residue changes as reference NODs. Putative NOD sequences from this study are as follows:

> RES_21_09_03_M2_MG_PACBIO_59399_L_6
MNIIKWIDQRKHWWKVFAVIFTVSIAVVGYIGYKTYQYAPPIADFVDEQGQPVFSAGSITSGQKIFLKYGLM
DYGSYLGDGGMRGPDFTAEALNLTARWMQQYYTKQQAEPAASERDARLQEQSLLARIQFELKENRYDEAT
NAVTVNAAQTFAFEQLVKFNAQRFGQGGDLAQKEVFKTKDYISDADEIRDLSAFFYWGGWFCAAQRPGY
DYSYTHNWPYDPLAGNIPHGGLVLWSAIGTLVVILSLGVIFYYYGKMDREAILQAQSAKMPPLATSELLDTF
KPTPTQRACYKFFAVAAVLFGIQVLAGLLAISDFVNLFAGLGIEITEWIPVTVSRAWHTQISILWIAMCWFAAT
IWVLPLICRPEPSGQRAWIETLFWMLVVAAVGGVVGMPLAVHGLLDFGDLWRWLGIQGWEFMQIGRFYQY
LLYAAFVVWLIITARGVWPVLKQKQTWSLPNWMVYSIIGIILMFSAGFVAGPETNFVIADFWRWCTIHMWV
EAFFEVFTTIIVAYFLYLMGFVSHVSAARVVYLSAVLFLGSGLIGISHNFYWNAKSIETVALGGVLSSLQVAPL
VLLTVEAWRFRHLPQSSLAQLKRRHGERGVFGLAEPFLFLVAVNFWNFMGAGVLGFMINLPIVNYYQHGTY
LTVNHAHAALFGVYGNLAIAAMLFCGRWVIGPERWNPRLLRTAFWSLNIGVSLMVVMDLFPVGVHQLIAV

MNDGYAFSRSQEYLSGSVFQTLTWLRGFGVVIFILGGVFPLLWFMVSRWFALKEPQAAAEAFVVPPSVLAV
AAFGSSNGNGEIFVGEPELVEQKI*
> RES_21_09_03_R3_M2_MG_PACBIO_23258_L_29-31
MEQKKGFIHFLMNPKKWWIPLTFIFVVSITGVGLIGYETYYEAPPIPDFINDSGEDVFTADEILEGQATFQRLA
LMDYGSMFGDGANRGPDFTAQALHQITTFITDYYANEPDLSKLAFLGVKEQVKRELRRNRYEQYTNQVKL
TDAQEYASQRLTDFYITYFTTTKFQQSLKKSYISNHEEIKKLSAFFFWGAWVCSTERPGKNYSYTHNWPFDP
DAGNLPGTPVIFWTIIGSLGLVIGLGIVLFYHGRLEKLNDRTFLDNSGPLMTENGVTQFVPTRSQRATYKFFF
AAMLVFLFQVIAGIITVHDFVNFTIFFGYDLRDLFPITISRGWHVQLSLLWISACWIGASLFITPFMVKESKNH
VSYVNTLFWLVVILVAGSVIGIYIGPMGLTKYWYWLGNQGWEYVEPGKLWQGLLFLIFALWAITLYRALRP
ALSLKQPWALPNWLVYTTVSILILLISGFIATPKTNFVIADFWRWCVIHMWAEAFFEVFTTVLVGYFMVLMG
LVSKESATRVIYLATLLFLGSGLLGISHNFYWNAKPVATMALGSVFSTLQVIPLVLLTLEAWRFQRLPKLAME
SMTTKNGTHDITTFGLQDVFLFLVGVNFWNFFGAGVMGFIINLPLANYYEHGTYLTVNHGHAALMGVYGN
LSLAAILFCLKLLTDSKAWNPSLVRLSFWSINIGLMLMVLLDLFPAGILQFQAVVDNGLWFARSNEFIEQETF
QTLTWLRIIGGAVFTLGGVLPLVYVVIKASAHLKNVTHKKLALEISQEEVLL*
> RES_21_09_03_R3_M2_MG_PACBIO_33983_L_19
MAKRNFIGYLLNPKNWWLPLLVVFIISIAGVTMIGIHTYTEAPPIPNYVSEKNETVFSKEDVLNGQAVFQKY
ALMEYGSMFGDGANRGPDYTAEALHYVAQYMNDYYHSQLKADANTDLLRKGVSEHVKSEIKNNRYAKK
NNGVLLSDAQVFATGELIKFYHEKFTNAASPGAFKPSGYITDADEIRSLTAFFFWGAWVCGVERPGEHYSYT
HNWPYDPSAGNTPASAIILWSIIGSLGLIFGLGLVMYYHGKLEKLDDSAYTTNARPFMSRGEIKKFQPDAIQR
STYKFFYVAILLFAVQVLAGILTVHDFVGLVNFFGFNISEPLPITVTRSWHVQLSLLWISACWIGASFFMMAL
VSPKQSRNQVTLINTIFWLTILLVAGSFAGMLLGPTGIIGKNWYWLGHQGWEYVEIGKLWQILLGIVFIIWAIT
IYRGIKPVMKLKQPWALPNWLVYTTFSIILLLISGFIATPKTNFVIADFWRWMVVHMWAEAFFEVFTTVLIGY
FMVMMGLVSKQAAIRVIYLATLLFLGSGLLGISHNFYWNAKPVGTMALGAVFSTLQVIPLVLLTLEAWRFSK
LPKTLENNNKVNGDLNKQFGFSEVFLFLVAVNFWNFFGAGVLGFIINLPIANYYEHGTYLTVNHGHAALMG
VYGNLALAAVLFCCQLLFKADWWRPRIIKTAFWSINIGLLLMVFLDLFPAGIWQFKTVTEEGLWYARSHTFI
ESAGFQTLTWLRIVGGSIFTVGGVIPLVWFITSRRKGLKKKDANMIVSSINEEKITASLVQY*
> RES_21_05_11_R3_SRF_MG_44_40
MANRNFISYLLNPKNWWLPLLIIFVVSITGVTMIGIHTYTEAPPIPNYISGKNETVFSKEEVLKGQAIFQKYAL
MEYGSMFGDGANRGPDYTAEALHHVTQYMNDYYNSKLNDAANELMRKGVGEQVKSEIKTNRHSRENNN
VALSDAQVFAATELIKFYNKKFTDPSFAGSFKPSGYITNADELRSLTSFFFWGAWVCGVERPGEHYSYTHNW
PYDPAAGNTPSAAIILWSIIGALGLILGLGFVLYYHGKLEKLDDNVYTNNAQPFMSRGEIKKFQPDEVQRVT
YKFFYVAIILFAVQVLAGILTVHDFVGFINFFGFNISEPLPITVTRSWHVQLSILWISACWIGASFFMMSLVSPK
QSSSQVKLINTIFWLCVVLVAGSLAGILLGPKGIIGKNWYWLGHQGWEYVEPGKIWQGILGIIFIIWAITLYRG
IKPVMKLKQPWALPNWLVYATFSIIFLLISGFIATPKTNFVIADFWRWMVVHMWAEAFFEVFTTVLIGYFMV
LMGLVSKQAATRVIYLATLLFLGSGLLGISHNFYWNAKPVFTMALGSVFSTLQVVPLVLLTLEAWRFSKLPK
VLETNNRINGDLNKRFGFSEVFLFLVAVNFWNFFGAGVLGFIINLPIANYYEHGTYLTVNHGHAALMGVYG
NLALAAVLFCCQLLFKAEWWKPRVVKTAFWSINIGLLLMVFLDLFPAGIWQFKTVTENGLWFARSHSFIESS
GFQTLTWLRIVGGSIFTVGGVIPLVWFITTRRKGLKEKIIEIKTVQKEYQIVSAIAADY*

#### Supplementary Tables

| Genome | Gene ID | TPM | Psortb prediction | Heme-binding domains | closest functional annotation |
|---|---|---|---|---|---|
| 447 | RES_21_05_11_R0_33-63CM_MG_783_22 | 4.22 | unknown | 5 | formate-dependent nitrite reductase (BLASTN, 68% identity, e-value 2 e-132) |

| | | | | | |
|---|---|---|---|---|---|
| 666 | RES_21_08_16_R1_M1_MG_1041_3 | 5.72 | Extracellular (9.04) | 5 | |
| 666 | RES_21_08_16_R1_M1_MG_657_7 | 0.29 | Extracellular (8.82) | 25 | DFE_0462 (40) |
| 167 | RES_21_09_03_R3_M2_MG_156341_1 | 4.08 | Extracellular (8.82) | 8 | OmcZ (76.9) |
| 167 | RES_21_09_03_R3_M2_MG_2232_5 | 0.25 | Extracellular (4.28) | 8 | OmcZ (76.9) |
| 167 | RES_21_09_03_R3_M2_MG_4091_1 | 1.44 | unknown | 5 | formate-dependent nitrite reductase (BLASTN, 73.2% identity, e-value 5.6 e-137) |
| 167 | RES_21_09_03_R3_M2_MG_4501_10 | 1.09 | unknown | 8 | |
| 167 | RES_21_09_03_R3_M2_MG_698_3 | 1.20 | Extracellular (9.04) | 12 | DFE_0449 (30) |
| 167 | RES_21_09_03_R3_M2_MG_698_4 | 3.74 | unknown | 5 | Best foldseek hit is *A. veneficus* nanowire (8E5G; E-value 7.44e-7) |

**Table S1. Characterization of expressed *M. nitroreducens* MHCs.** For psortb predictions, scores are given in parentheses. The functions of these proteins are unknown, but best functional predictions are given when available. For HMMs, the detection thresholds are given in parenthesis. HMMs were called using metabolic-G. TPM = transcripts per million. Genome 447 is RES_21_05_11_R0_33-63CM_MG_Methanoperedens_41_50, genome 666 is RES_21_08_16_R1_M1_MG_Candidatus_Methanoperedens_nitroreducens_46_7 and genome 167 is RES_21_09_03_R3_M2_MG_Candidatus_Methanoperedens_nitroreducens_44_69. Full genome information is in Data S1.

| Date | Application / event | Rate |
|---|---|---|
| 5/11/2021 | Nitrogen fertilizer applied as aqueous ammonia | 150 lbs/acre |
| 5/13/2021 | M206 seed applied | 150 lbs/acre |
| 5/14/2021 | Started water | — |

| | | |
|---|---|---|
| 5/16/2021 | Field was flooded | — |
| 5/23/2021 | LAMBDA-CY AG insecticide applied | 3.8 oz/acre |
| 5/25/2021 | BUTTE herbicide (Gowan) applied | 7.5 lbs/acre |
| 6/10/2021 | Phosphorus fertilizer applied | 40.5 lbs/acre |
| 6/10/2021 | Potassium fertilizer applied | 35 lbs/acre |
| 6/10/2021 | Zinc fertilizer applied | 15 lbs/acre |
| 6/25/2021 | SUPERWHAM! 41.2 % propanil herbicide applied | 1.5 gal/acre |
| 6/25/2021 | Grandstand R herbicide applied | 4 oz/acre |
| 8/4/2021 | Quadris fungicide (Syngenta) applied | 12 oz/acre |
| 9/5/2021 | Drained water | — |

**Table S2. Field treatments.** Date, description, and application rate of field treatments across the 2021 growing season.

### Description of Supplementary datasets

#### Description of the data and file structure

The dataset is divided into nine main data tables (Data S1–S10), each described below. All files are provided as .xlsx, .csv, or .tsv tables and are linked to the analytical framework outlined in the main text and supplementary methods of the publication. Large files are best explored in R or Python.

**Data S1:** Environmental metadata

A) Particle size
B) $CH_4$, $N_2O$, $CO_2$, and $NH_3$ fluxes
C) Soil chemistry

**Data S2:** Genome descriptions, cohort membership, and statistics.
**Data S3:** Expression of TEAP markers and *pmoB* from 95% ID dereplicated protein set
**Data S4:** Genome and 99% ID vsearch cluster membership of ribosomal protein S3 sequences, as well as cluster coverage (length- and library- normalized coverage of member sequence with the longest scaffold, summed across replicates and time points), and best NCBI NR taxonomic hit of cluster centroid.
**Data S5:** Enrichment and depletion of Kegg Orthologies (KOs) across genome cohorts
**Data S6:** Total expression of most highly expressed annotated genes from the most active cohorts (tan, sienna3, darkgreen, greenyellow, black, and green).
**Data S7:** Time-resolved expression of hydrogenases from rice paddy genomes
**Data S8:** Time-resolved expression of TEAP markers from genome set
**Data S9**: iRep estimated genome replication rates.
**Data S10**: Description of PacBio HiFi genomes from deep sequencing of sample RES_21_09_03_R3_M2
